# Rapid peripheral reprogramming of myelinated afferents drives human hyperalgesia

**DOI:** 10.64898/2026.02.05.703527

**Authors:** O. Bouchatta, A. Marshall, S. H. Lindström, J. Kupari, H. Manouze, A. Barakat, H. Yu, I. Szczot, J. Johansson, R. Saager, M. Larsson, W. Luo, A. G. Marshall, P. Ernfors, H. Olausson, S. S. Nagi, NIH PRECISION Human Pain Network

**Affiliations:** Department of Biomedical and Clinical Sciences, Linköping University, Linköping, Sweden; Pain Research Institute, Institute of Life Course and Medical Sciences, University of Liverpool, Liverpool, United Kingdom; Department of Medical Biochemistry and Biophysics, Division of Molecular Neurobiology, Karolinska Institute, Stockholm, Sweden; Department of Pharmacology and Toxicology, Faculty of Pharmacy, Assiut University, Assiut, Egypt; Department of Neuroscience, Perelman School of Medicine, University of Pennsylvania, Philadelphia, USA; Department of Biomedical Engineering, Linköping University, Linköping, Sweden

## Abstract

A small burn can render a large area of skin painfully tender. This widespread hyperalgesia protects injured tissue and is typically attributed to altered spinal cord mechanisms. Whether peripheral sensory afferents directly interact to contribute to hyperalgesia remains unclear. Using single-unit microneurography, we recorded cutaneous afferents before and after inducing localized TRPV1-mediated inflammatory flare. This triggered minutes-scale reweighting of transcriptomically defined TRPV1^−^ afferents, with divergent effects among myelinated (Aβ-range) classes: tactile receptors showed reduced responsiveness, whereas mechano-nociceptors underwent sensitization. These changes paralleled diminished tactile sensitivity and intensified mechanical pain. Recruitment of mechanically silent branches in Aβ-range mechano-nociceptors produced wide-field amplification of peripheral nociceptive signaling beyond the inflamed site. These findings suggest that rapid peripheral crosstalk from TRPV1^+^ afferents reprograms TRPV1^−^ Aβ-afferents and drives human hyperalgesia.

## Introduction

Painful tenderness after tissue injury is a multi-level protective response that limits further damage during healing. Early work by Lewis distinguished two key components: a local flare surrounding the injured skin, and a broader zone of hyperalgesia extending beyond the site of injury (*1*). The flare is well established as an axon-reflex response mediated by unmyelinated (C) fibers, whereas the mechanisms generating injury-evoked hyperalgesia have long been debated, with peripheral and central origins proposed (*1, 2*). A conceptual distinction later emerged between *primary* hyperalgesia at the site of injury, which was attributed to sensitization of peripheral nociceptor endings, and *secondary* hyperalgesia in surrounding tissue, which was attributed to amplification of pain signaling within dorsal horn circuits of the spinal cord (*3, 4*), while implicitly treating peripheral afferent encoding outside the injury site as unchanged.

Microneurography in humans refined this framework by revealing the peripheral basis of primary heat hyperalgesia. Following intradermal administration of capsaicin, C-mechanoheat (CMH) nociceptors demonstrate lowered heat thresholds within the injury zone, paralleling behavioral sensitization. In contrast, despite the robust emergence of mechanical hyperalgesia, CMH nociceptors do not exhibit lowered mechanical thresholds or enhanced mechanical firing (*5, 6*). Thus, while peripheral CMH sensitization accounts for primary heat hyperalgesia, the peripheral afferent substrate for human mechanical hyperalgesia remains unresolved.

One longstanding doctrine in somatosensory neuroscience holds that thickly myelinated (Aβ) afferents encode innocuous touch, whereas thinly myelinated (Aδ) and unmyelinated (C) fibers signal pain. We recently challenged this dichotomy by identifying human Aβ-range afferents that encode mechanical pain. Using single-unit microneurography, we found that these neurons require strong mechanical stimulation to activate, respond vigorously when stimulation reaches painful levels, and conduct signals at Aβ velocities – comparable to touch fibers – allowing rapid transmission of pain information (*7*). We termed these neurons ultrafast nociceptors (UFNs). One subset responds to cooling, whereas none respond to heating (*8, 9*). Recent single-cell RNA-sequencing (scRNA-seq) analyses of human dorsal root ganglion (DRG) neurons have identified a molecularly distinct subclass of UFNs, the A-PEP.KIT population, which lacks expression of the heat- and capsaicin-gated ion channel TRPV1 – a molecular signature consistent with its physiological heat insensitivity (*8-10*).

As seen in UFNs, single-cell–resolved classification of human sensory neurons further reveals that tactile Aβ low-threshold mechanoreceptors (LTMRs) similarly lack TRPV1 expression, whereas multiple C-fiber populations show enriched TRPV1 levels (*8, 10*). These data suggest that the inflammatory pain signaling induced by heat or capsaicin is asymmetrically organized across afferent classes. Although TRPV1 expression in skin is not restricted to sensory neurons (*11*), the inflammatory flare in humans is closely linked to neural activation and can be attributed to a specific afferent class, the C mechano-insensitive (CMi) nociceptors (*12*). This signaling could alter the functional state of neighboring TRPV1^−^ A-fiber populations (*13*). Furthermore, peripheral sensory input also appears to undergo reconfiguration during pain states, as indicated by studies using rodent nerve-injury models (*14*). The rodent models demonstrate that A-fiber mechano-nociceptors show gain of function, whereas A-LTMRs exhibit desensitization, consistent with a population-level recalibration of large-fiber signaling (*15*).

These observations, together with the discovery of UFNs and the classification of molecularly defined human afferent classes, prompted us to ask whether a previously unrecognized peripheral mechanism could explain mechanical hyperalgesia in humans. Specifically, we aimed to determine whether acute inflammation – evoked by heat or capsaicin – may engage local interactions between TRPV1^+^ and TRPV1^−^ afferents within human skin. In the present study, we combined single-unit microneurography with noxious heating or topical capsaicin to elicit a localized inflammatory flare. We also quantified the accompanying hemodynamic responses and used psychophysical testing to assess the perceptual consequences. Tracking the same identified afferent fibers before and after inflammation confirmed that distinct classes of Aβ afferents undergo rapid and coordinated changes in responsiveness that parallel the development of the inflammatory flare and its perceptual correlates. These findings demonstrate that the induction of hyperalgesia involves a previously unrecognized peripheral mechanism, in which rapid crosstalk from TRPV1^+^ afferents reprograms TRPV1^−^ myelinated afferents, challenging the current view that widespread hyperalgesia after tissue injury arises from central sensitization.

## Results

We recorded single identified afferents from the radial or peroneal nerves of healthy participants using microneurography (Fig. 1A). Mechanically responsive units were classified as myelinated (A) or unmyelinated (C), low- or high-threshold mechanoreceptors (LTMRs or HTMRs). Consistent with prior work, we identified UFNs with conduction velocities overlapping those of Aβ-LTMRs (Fig. 1B). In addition, CMi nociceptors, which are not accessible with mechanical search, were recorded using an electrical search/latency-tracking approach. Where possible, physiological afferent classes were aligned with corresponding human DRG transcriptomic populations.

**Fig. 1.**
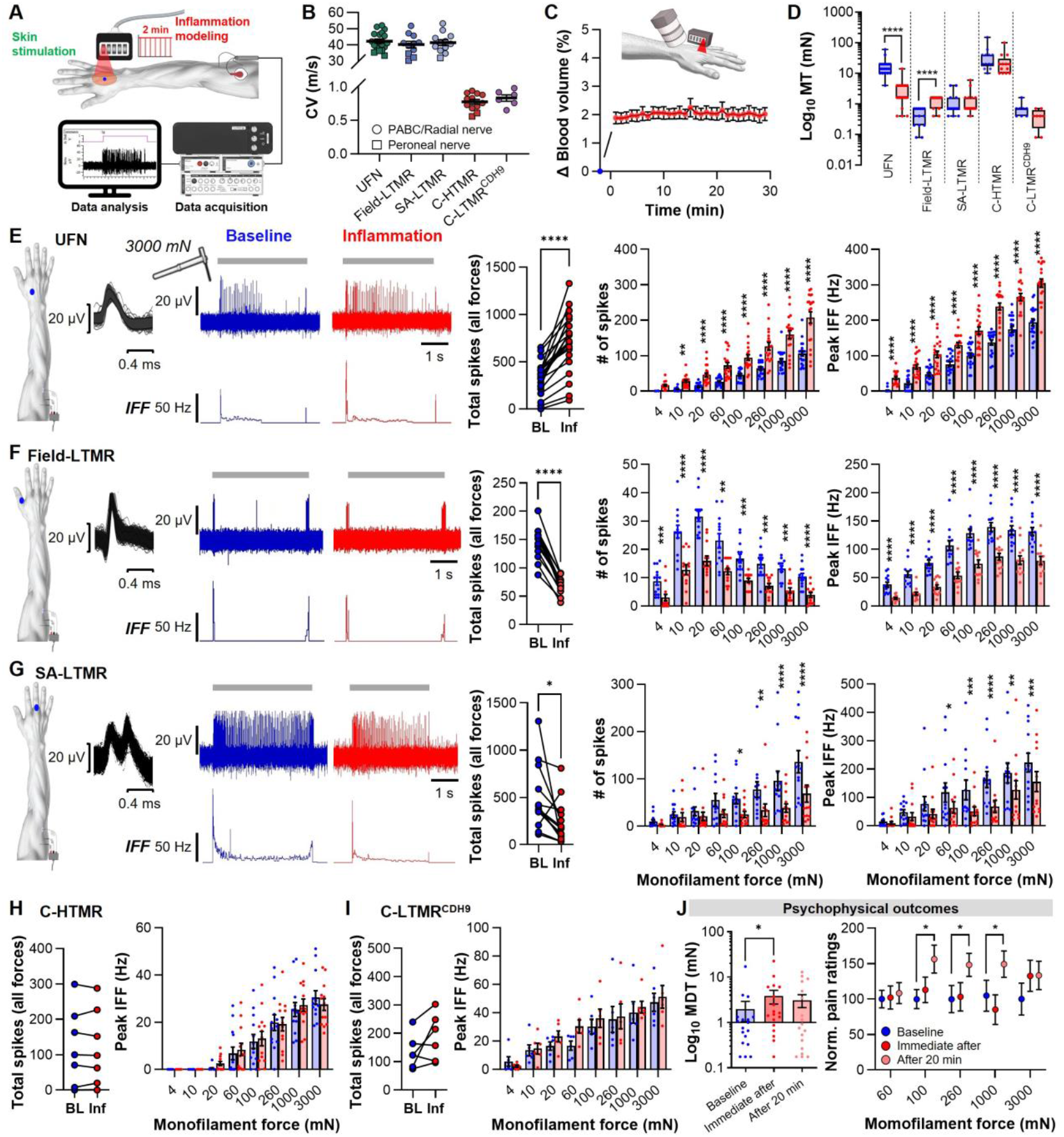
Heat-induced inflammation sensitized UFNs and desensitized Aβ-LTMRs, in alignment with perceptual shifts. (**A**) Schematic of microneurography setup used to record single mechanosensory afferents (radial nerve shown) in awake human participants. After characterizing the afferent’s receptive field (RF, blue dot), the thermode was placed onto the RF to deliver the 2-min 50 °C heat-induced inflammation protocol. (**B**) Conduction velocities (CVs) of identified afferents. UFN CVs (n = 19) did not differ from Aβ-LTMRs (Field-LTMR: q = 1.65, p = 0.7678, n = 11; SA-LTMR: q = 0.73, p = 0.9851, n = 12; RM one-way ANOVA with Tukey’s test). PABC: posterior antebrachial cutaneous. (**C**) Spatial frequency domain imaging (SFDI) assessed cutaneous blood volume following heat-induced inflammation protocol (as in A) delivered to the dorsal hand. Heating induced a sustained increase in blood volume that persisted for ≥30 min. (**D**) Mechanical thresholds (log_10_ mN) before (blue) and after (red) heat-induced inflammation. UFNs showed robust sensitization (t = 6.58, p < 0.0001; n = 20; Paired t-test), whereas Field-LTMRs were desensitized (t = 8.52, p < 0.0001; n = 14). No significant threshold changes were observed for SA-LTMRs (t = 2.01, p = 0.3482; n = 14), C-HTMRs (t = 2.58, p = 0.1484; n = 13), or C-LTMR^CDH9^ (t = 2.34, p = 0.1189; n = 8). Plotted as individual values with median and interquartile range. (**E–G**) Characterization of responses to graded punctate indentation (4-3000 mN) at baseline (blue) and following heat-induced inflammation (red) across afferent subtypes. Far left: RF location (blue dot) and superimposed spike activity during RF stimulation (black traces) of each representative unit. Representative traces show raw spike trains (above) and instantaneous firing frequency (IFF) plots (below) in response to a 3000-mN indentation lasting approximately 5 seconds (grey bars). Quantitative analysis of all units is presented as *total spike count across all forces* (individual units) and with *responsiveness to each force* (number of spikes or peak IFF). (**E**) UFNs exhibited robust increases in total spike count (t = 7.63, p < 0.0001; Paired t-test) and responsiveness across forces (number of spikes: F_(7,113)_ = 21.71, p < 0.0001; peak IFF: F_(7,113)_= 18.66, p < 0.0001; RM two-way ANOVA). (**F**) Field-LTMRs showed robust decreases in total spike counts (t = 14.05, p < 0.001) and responsiveness across forces (number of spikes: F_(3560,4272)_ = 8.61, p < 0.0001; peak IFF: F_(7,84)_ = 12.71, p < 0.0001). (**G**) SA-LTMRs showed modest reductions in total spike counts (t = 2.22, p < 0.05) and responsiveness across forces (number of spikes: F_(7,84)_ = 3.95, p < 0.001; peak IFF: F_(7,84)_ = 3.40, p = 0.003). (**H**–**I**) C-HTMRs and C-LTMR^CDH9^ showed no change in total spike counts or peak IFF across forces (all non-significant). Asterisks (where shown) on the force-response data (E-I) indicate significant post hoc comparisons (Sidak’s test). (**J**) Psychophysics paralleled afferent reweighting: Mechanical detection thresholds (MDT) doubled immediately after inflammation (F_(1695,2881)_ = 3.48, q = 2.83, p = 0.0210; n = 18; RM one-way ANOVA with Dunnett’s test), reflecting reduced tactile sensitivity, whereas pain ratings to noxious monofilaments were elevated 20 min post-inflammation (F_(5392,8627)_ = 1.68; 100 mN: q = 4.5, p = 0.0141; 260 mN: q = 3.83, p = 0.0287; 1000 mN: q = 3.78, p = 0.0413; RM two-way ANOVA with Tukey’s test). All data shown as individual values with mean + SEM, unless noted. *p < 0.05, **p < 0.01, ***p < 0.001, ****p < 0.0001.

Transcriptomic profiling has identified a UFN subset distinguished by KIT expression. These UFN^A-PEP.KIT^ afferents co-express the mechanically gated PIEZO2 and cold-gated TRPM8 ion channels, innervate human hair follicles, and respond to hair pull, skin indentation, and cooling (*8, 9*). In contrast, UFN^KIT−^ afferents represent a molecularly heterogeneous population that responds to punctate indentation but lacks hair-pull and cooling sensitivity (*8, 9*). UFNs lacking cooling sensitivity (UFN^KIT–^) are not yet molecularly resolved (*8-10*). Rapidly adapting Aβ hair-follicle (HF) LTMRs align with the Aβ-LTMR.CCKAR population (henceforth referred to as Aβ-LTMR(HF)^CCKAR^), while C-LTMRs correspond to the C-LTMR.CDH9 population (henceforth referred to as C-LTMR^CDH9^) (*10*). Transcriptomic identities distinguishing Aβ Field- and slowly adapting (SA-) LTMRs remain unresolved in current human DRG datasets (*10*); therefore, they are referred to using established physiological nomenclature (*16*). Because the C-HTMR population recorded here was functionally heterogeneous, comprising mechano-heat, mechano-cool, and polymodal units, we retained the broad physiological classification.

### Cutaneous heating produced sustained inflammation, shifted mechanical responsiveness of UFNs and LTMRs, and elicited perceptual changes

Microneurography recordings were performed at baseline and following a heat-induced inflammation protocol (Fig 1A). Heating for two minutes at 50°C generated a persistent vascular flare and increased local blood volume lasting at least 30 min, confirming a sustained inflammatory response (Fig. 1C and Fig. S1). In contrast, the increased skin temperature declined rapidly toward baseline after cessation of heating (Fig. S1).

Inflammation induced distinct, class-specific changes in mechanical sensitivity. Thresholds decreased in UFNs, indicating sensitization, whereas thresholds increased in Field-LTMRs. Mechanical thresholds were unchanged in slowly adapting (SA-) LTMRs, C-LTMR^CDH9^, and C-HTMRs (Fig. 1D). UFNs also exhibited increased firing to punctate indentation across innocuous and noxious forces (Fig. 1E). For both mechanical thresholds and graded punctate indentation, UFN^A-PEP.KIT^ and UFN^KIT−^ afferents showed indistinguishable baseline responses and inflammatory modulation and were therefore pooled. In contrast to UFNs, Field- and SA-LTMR responses were attenuated (Fig. 1F-G), whereas C-LTMR^CDH9^ and C-HTMR response profiles were unaffected by inflammation (Fig. 1H-I and Fig. S2). Finally, prolonged after-discharges were observed in CMi nociceptors (Fig. S3). The changes in Aβ-LTMRs and UFNs corresponded to observed perceptual shifts, as tactile detection thresholds nearly doubled, reflecting reduced tactile sensitivity, while mechanical pain ratings increased after inflammation (Fig. 1J). No spontaneous pain was reported at baseline. While the heat-induced inflammation protocol was painful (Fig. S4B), after inflammation, spontaneous pain was only reported by 3 out of 18 participants (using a 0-100 numerical rating scale; immediately: NRS 3-50/100; 20 min: NRS 1-15/100).

Similar patterns were observed for mechanical sensitivity to hair-pull and brushing. During hair-pull stimulation, the UFN^A-PEP.KIT^ afferents showed strong baseline sensitivity, which increased after inflammation, while after inflammation UFN^KIT−^ afferents developed a de novo but significantly weaker hair-pull response (F_(1000, 6000)_ = 86.76, p < 0.0001; RM two-way ANOVA). Field-LTMRs remained unresponsive under both conditions. SA-LTMRs showed reduced activity after inflammation, while C-LTMR^CDH9^ and C-HTMR responses were unchanged (Fig. S5). In brushing assays, inflammation induced de novo sensitivity to low-intensity (soft-brush) stroking and enhanced sensitivity to high-intensity (coarse-brush) stroking in UFNs, while attenuating brushing responses in Field-LTMRs, SA-LTMRs, and C-HTMRs. By contrast, the Aβ-LTMR(HF)^CCKAR^ and C-LTMR^CDH9^ types maintained stable responsiveness to brushing (Fig. S6). Brush-evoked pain to single low-intensity stroking was uncommon during psychophysical testing (2/18 participants; NRS 6–48/100).

Together, these data showed that heat-induced inflammation rapidly reconfigured mechanical signaling in myelinated afferents, as characterized by sensitization of UFNs and desensitization of Aβ-LTMR subtypes, whereas CMi nociceptors exhibited prolonged after-discharge consistent with sustained activation during neurogenic inflammation.

### Heat-induced inflammation enhanced cold responsiveness in UFNs and impaired thermal discrimination

Heat-induced inflammation markedly altered cold responsiveness in both UFN classes. UFN^A-PEP.KIT^ afferents exhibited enhanced cooling responses (Fig. 2A), whereas UFN^KIT−^ afferents developed a de novo but significantly weaker cooling sensitivity (Fig. 2B; F_(3,36)_ = 54.16, p < 0.0001; RM one-way ANOVA). The C-HTMRs, C-LTMR^CDH9^, and SA-LTMRs all displayed baseline cooling responses; however, clear modulation following inflammation was observed only in SA-LTMRs, which showed reduced cooling-evoked activity (Fig. 2C–E). Field-LTMRs remained cold-insensitive across both conditions (Fig. 2F).

**Fig. 2.**
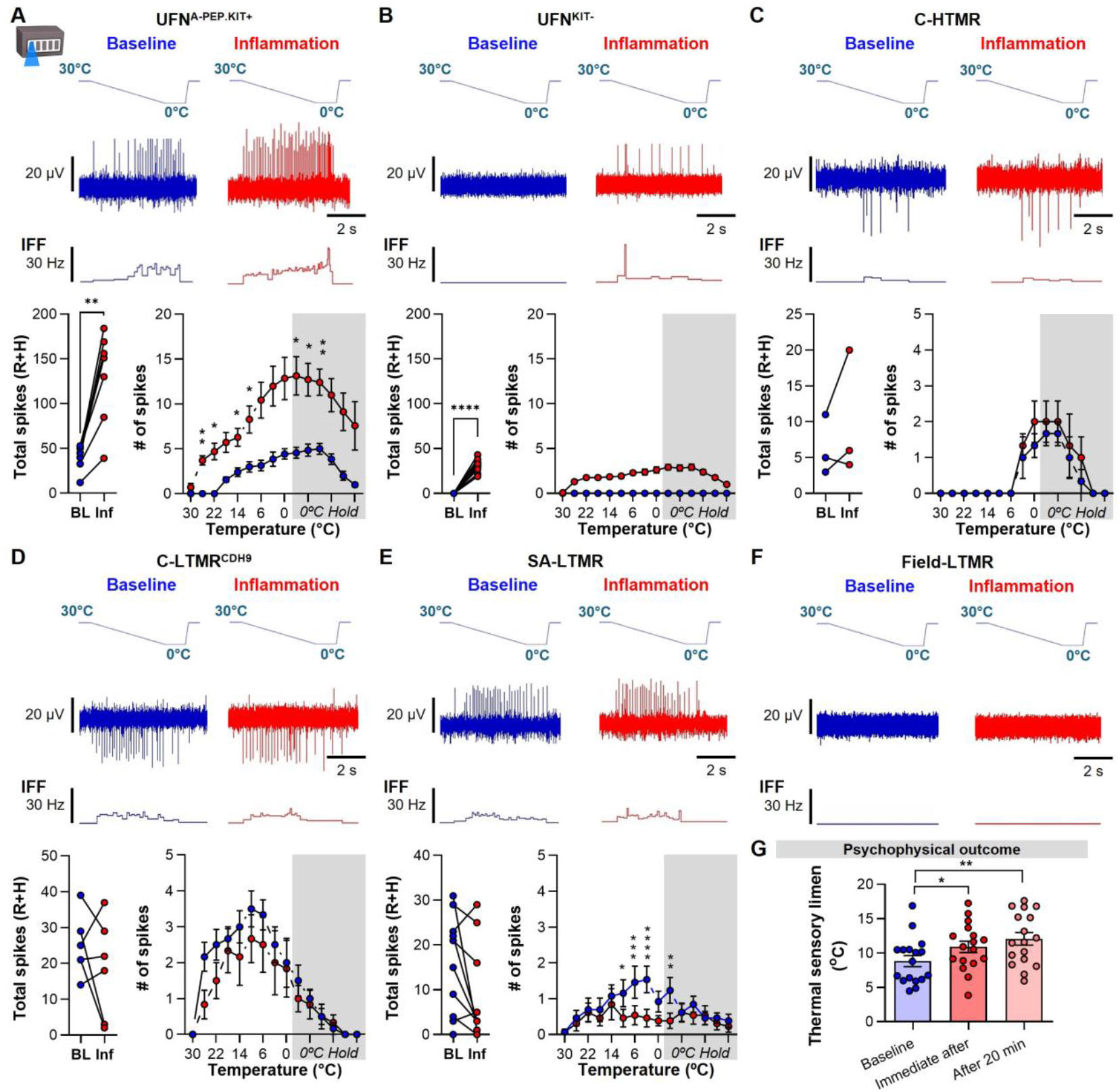
Heat-induced inflammation reshaped UFN and SA-LTMR cold sensitivity and impaired thermal discrimination. (**A**–**F**) Characterization of cooling responses (30 °C → 0 °C; 5 s hold) at baseline (blue) and following heat-induced inflammation (red) across afferent subtypes. Top: Ramp-and-Hold (R+H) cooling stimulus plotted above representative traces showing raw spike trains (above), and IFF plots (below). Bottom: Quantitative analysis of all units is presented as *total spike count across R+H phases* (individual units) and *temperature-response curves* (spike output resolved in 1-s bins). (**A**) UFN^A-PEP.KIT^ exhibited enhanced cooling sensitivity (t = 5.93, p = 0.001, n = 7; Paired t-test), with a significant upward shift in the temperature-response function (F_(2946,1767)_ = 3.97, p = 0.0255; RM two-way ANOVA). (**B**) UFN^KIT−^ acquired a de novo cooling response after inflammation (t = 15.00, p < 0.0001, n = 13), with a significant upward shift in the temperature-response function (F_(3830,4596)_ = 13.40, p < 0.0001). (**C-D**) C-HTMR (n = 3) and C-LTMR^CDH9^ (n = 6) showed no change (p > 0.05) in cooling-evoked responsiveness following inflammation, either in total spikes counts or in the temperature-response function. (**E**) Inflammation reduced cooling-evoked activity in SA-LTMRs (t = 2.04, p = 0.0629; n = 13), with a significant downward shift in the temperature**-**response function ≤10 °C (F_(14,168)_ = 2.78, p = 0.0010). (**F**) Field-LTMR remained cold-insensitive under both baseline and inflammation conditions (n = 13). Asterisks (where shown) on the temperature-response curves (A-G) indicate significant post hoc comparisons (Sidak’s test). (**G**) Thermal sensory limen (the minimum temperature difference needed to reliably distinguish warm from cool) increased after inflammation (F_(1586,2538)_ = 7.9; baseline *vs*. immediately after: q = 3.09, p = 0.0129; baseline *vs*. after 20 min: q = 4.28, p = 0.0011; RM one-way ANOVA with Dunnett’s test). All data shown as individual values with mean + SEM, unless noted. *p < 0.05, **p < 0.01, ***p < 0.001, ****p < 0.0001.

Where heating was applied as a test stimulus (30–50 °C ramp with 5-s hold), no heat-evoked responses were observed in A-mechanoreceptors at baseline or after inflammation, whereas heat-evoked responses in C-mechanoreceptors were either unchanged or enhanced (Fig. S7).

Psychophysical testing revealed no changes in cold detection and pain thresholds. However, significant warm hypoesthesia and heat hyperalgesia were observed immediately after inflammation (Fig. S4). Further, inflammation produced a deficit in thermal discrimination, as assessed using the thermal sensory limen task, which measures the ability to detect alternating cool and warm stimuli (Fig. 2G).

Taken together, these findings showed that heat-induced inflammation heightened cold sensitivity in UFN^A-PEP.KIT^ afferents, unmasked a new cooling response in UFN^KIT−^ afferents, and was accompanied by impaired thermal discrimination. In contrast, UFNs remained unresponsive to heating under both baseline and following heat-induced inflammation.

### Heat-induced inflammation reconfigured afferent signaling across modalities and classes

Heat-induced inflammation drove broad, modality-spanning shifts in sensory signaling across afferent classes. Fig. 3A summarizes afferent activity evoked by brushing, graded punctate indentation, graded hair pull, and thermal stimulation (cooling and heating) for each afferent class at baseline and in the same recordings after the induction of inflammation. To visualize shifts in response profiles, we segregated pain-sensing (UFN^A-PEP.KIT^, UFN^KIT−^, C-HTMR) and touch-sensing (Field-LTMR, SA-LTMR, C-LTMR^CDH9^) afferents and generated UMAPs for each grouping. Following inflammation, the UFN^A-PEP.KIT^, UFN^KIT−^, Field-LTMRs and SA-LTMRs exhibited clear shifts in their UMAP positions relative to baseline, consistent with their modality-specific changes in responsiveness, whereas C-fiber populations showed limited displacement (Fig. 3B).

**Fig. 3.**
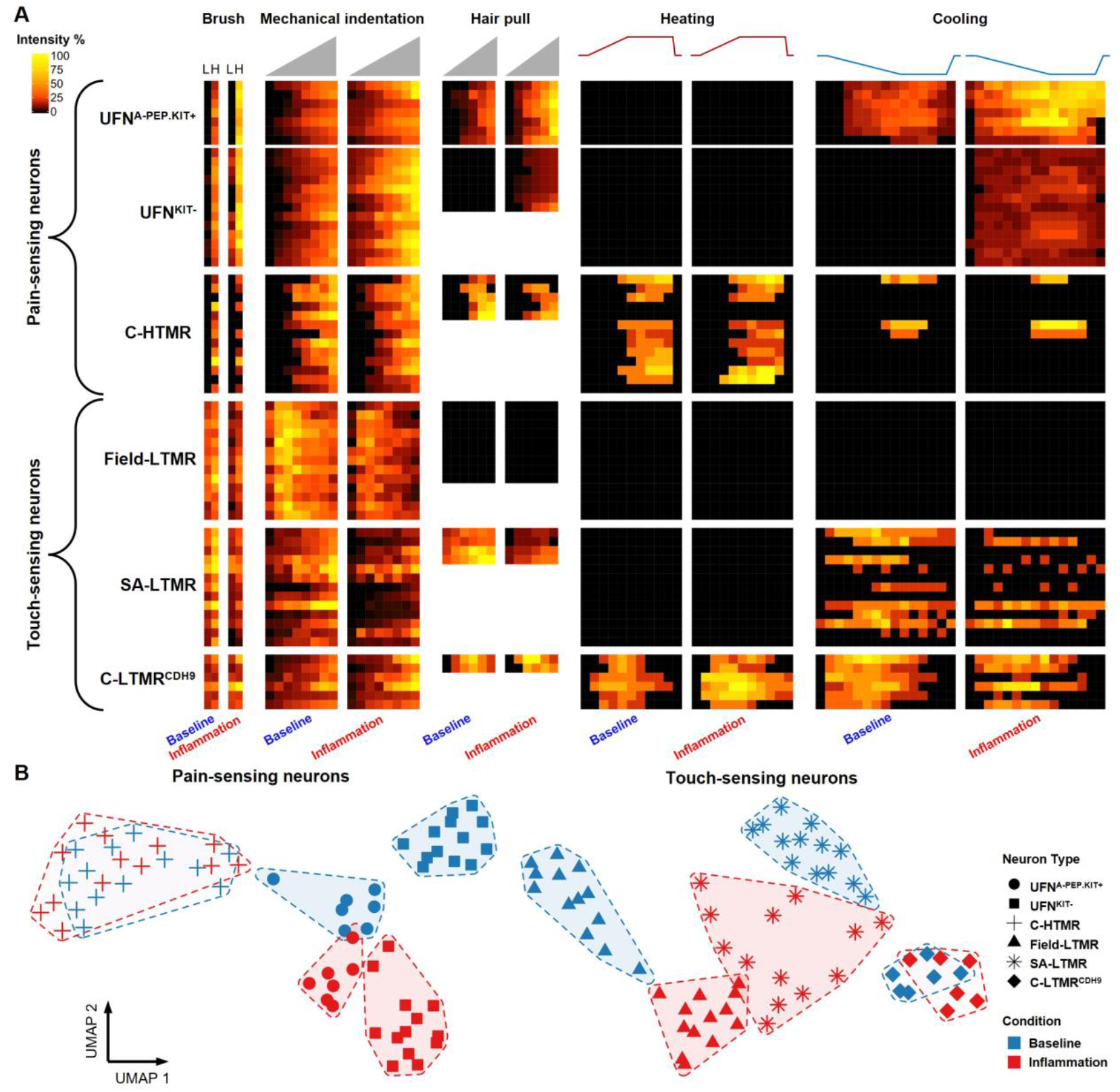
Multimodal response profiling revealed inflammation-related shifts concentrated in UFNs and Aβ-LTMRs. (**A**) Heatmaps summarizing stimulus-evoked activity across afferent classes at baseline (blue) and during heat-induced inflammation (red). Stimulus modalities span low (L) / high (H) intensity brushing, graded punctate indentation (4, 10, 20, 60, 100, 260, 1000 and 3000 mN), graded hair pull (10, 20, 50, 100, 200 and 400 mN) and thermal stimuli (cooling to 0°C and heating to 50°C with R+H protocol, as depicted). Responses are normalized within modality (color scale, % Intensity). (**B**) UMAP projections based upon response profiles after segregation into pain-sensing neurons (left; UFN^A-PEP.KIT^, UFN^KIT−^, C-HTMR) and touch-sensing neurons (right; Field-LTMR, SA-LTMR, C-LTMR^CDH9^). Units formed distinct groups when visualized by type (shape) and experimental condition (baseline, blue vs. inflammation, red), revealing inflammation-related shifts in the modality tuning of Field-LTMRs, SA-LTMRs and both UFN types. In contrast, C-HTMR and C-LTMR^CDH9^ groupings remained comparatively stable.

### Heat-induced inflammation expanded the UFN receptive fields and the area of mechanical hyperalgesia

Heat-induced inflammation produced a pronounced expansion of UFN receptive fields (RFs), as reflected by a many-fold increase in the cutaneous territory responsive to punctate indentation, which coincided with the rapid emergence of vascular flare zones (Fig. 4A–D). RF expansion was not concentric and instead displayed a proximal bias. Responses within the expanded RF could be evoked at the same reduced mechanical thresholds observed at the primary site. The UFN^A-PEP.KIT^ and UFN^KIT−^ afferents showed similar RF expansion and inflammatory modulation and were therefore pooled. In contrast to UFNs, most C-HTMRs showed reductions in RF size, while effects in other afferent classes were subtle and variable (Fig. 4C). Psychophysical testing revealed an enlargement of the skin area in which punctate mechanical stimulation was perceived as painful, with hyperalgesia emerging immediately and persisting for at least 20 min (Fig. 4E).

**Fig. 4.**
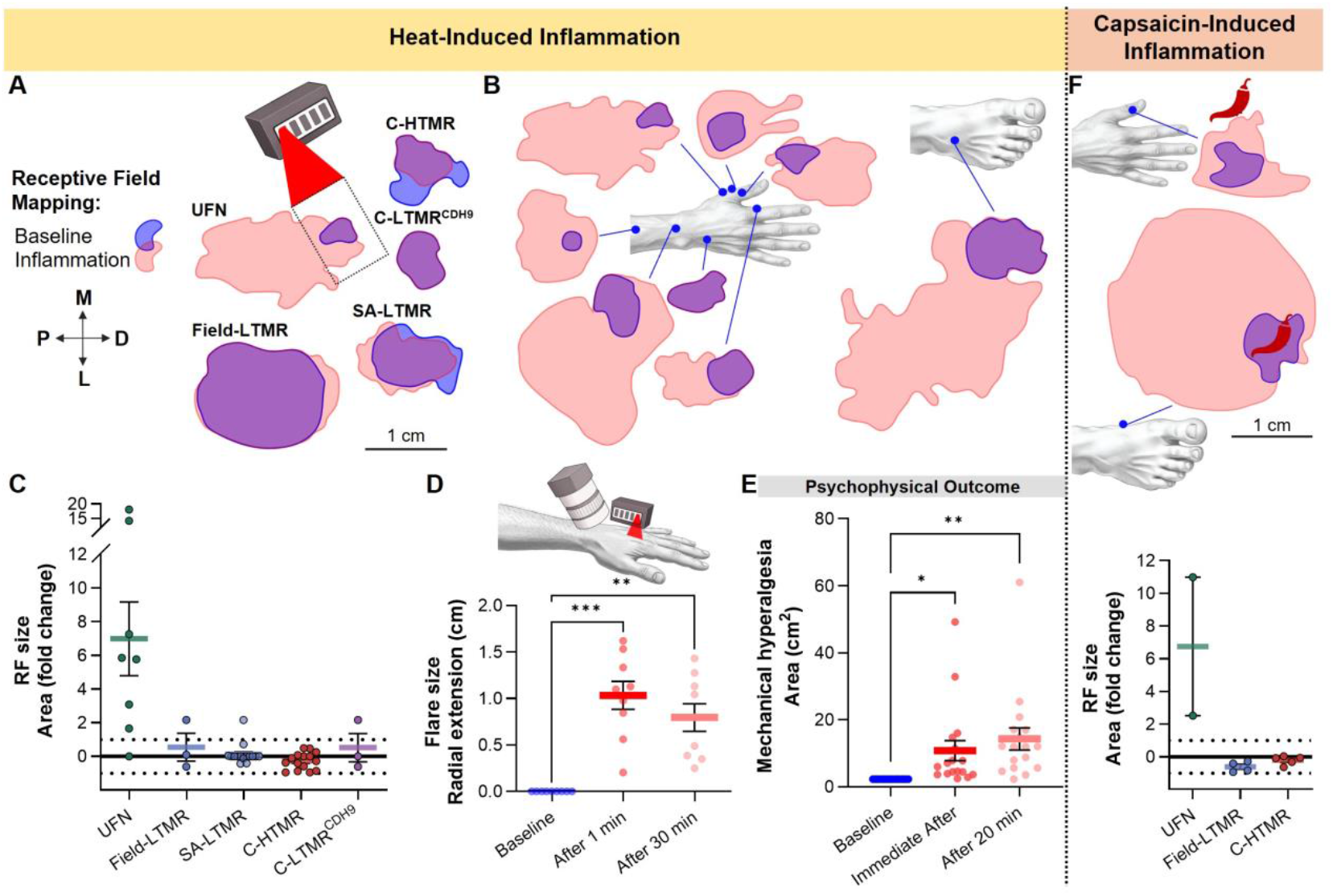
Heat-induced inflammation evoked expansion of UFN receptive fields and secondary hyperalgesia. (**A**) Example RF maps for multiple afferent classes (UFN, Field-LTMR, C-HTMR, C-LTMR^CDH9^, and SA-LTMR). Boundaries of the mechanical RFs are shown for baseline (blue) and post-inflammation (red) with overlap (purple). Rectangular thermode footprint is plotted as a dashed outline over the RF of the example UFN. Plotted with matched scale and orientation (P, proximal; M, medial; D, distal; L, lateral). (**B**) Spatial organization and location of all recorded UFN RFs (relative scaling). (**C**) Fold-change in RF area following heat-induced inflammation. UFNs exhibited robust RF enlargement (n = 8). In contrast, most C-HTMRs RFs were reduced (9/16), while a subset expanded (3/16) or remained unchanged (4/16). Field-LTMR (n = 3), and C-LTMR^CDH9^ (n = 3) showed variable changes, while SA-LTMR (n = 15) were typically unchanged. (**D**) SFDI measurements of the radial extent of erythema revealed flare development following heat-induced inflammation (F_(1427,1182)_ = 34.10, p < 0.0001, n = 9; RM one-way ANOVA). (**E**) An area of punctate mechanical hyperalgesia developed immediately after heat-induced inflammation and increased further at 20 min (F_(1431,2289)_ = 9.99, p = 0.0019; n = 18; RM one-way ANOVA). Asterisks (where shown, in D,E) indicate significant post hoc comparisons (Dunnett’s test). (**F**) Topical capsaicin induced similar changes in RF size. Top: Location and spatial expansion of 2 UFN RF maps after capsaicin-induced inflammation 5 mm outside of RF or overlapping the RF (chili icons). Bottom: Fold-change in RF area following capsaicin-induced inflammation. UFNs (n = 2) exhibited robust RF enlargement, while Field-LTMRs (n = 4), and C-HTMRs (n = 5) frequently showed RF reduction. All data plotted as individual values and mean ± SEM. *p < 0.05, **p < 0.01, ***p < 0.001.

We assessed whether TRPV1 activation alone was sufficient to drive UFN sensitization by applying topical capsaicin within the baseline RF. Despite lacking TRPV1 expression (*8*) and exhibiting no heat-evoked activity (Fig. 3A and Fig. S7A), the UFNs displayed a robust expansion of RF size and marked increases in mechanical sensitivity across multiple punctate force levels following capsaicin application (Fig. 4F and Fig. S8A-B). Notably, applying capsaicin 5 mm outside the original RF also resulted in RF expansion and enhanced responsiveness to both mechanical and cooling stimuli (Fig. 4F and Fig. S8C-D). In contrast, Field-LTMRs desensitized following capsaicin application, as observed after heat-induced inflammation, exhibiting elevated mechanical thresholds and reduced responses across punctate force levels, accompanied in this case by a consistent reduction in RF size (Fig. 4F and Fig. S8E–F).

Together, these findings showed that heat-evoked inflammation or topical TRPV1 activation produced a strong, many-fold enlargement of UFN RFs and an expansion of the area of mechanical pain. This effect was selective for UFNs, as RFs in other afferent classes were unchanged or showed reductions. These parallel changes underscore a close correspondence between UFN sensitization and the spatial spread of hyperalgesia in humans.

### Heat-induced inflammation redistributes mechanoreceptor drive via local signaling mechanisms

We assessed how acute inflammation alters the balance of peripheral mechanoreceptor signaling by quantifying the relative contributions of Aβ-LTMRs (including Field- and SA-LTMRs), C-LTMR^CDH9^, C-HTMRs, and UFNs to mechanical encoding across brushing, indentation, and hair-pull stimuli, with each stimulus examined at low and high intensities. At baseline, Aβ-LTMRs dominated the encoding of low-intensity mechanical stimuli, whereas UFNs contributed disproportionately to high-intensity stimuli, particularly during indentation and hair pull. Heat-induced inflammation produced a robust redistribution of afferent contributions across modalities (Fig. 5A). UFNs accounted for a significantly larger fraction of mechanoreceptor signaling after inflammation compared with baseline across stimulus types and intensities (beta regression model odds ratio: 4.3, p < 0.001), with large shifts observed for both low- and high-intensity stimuli. Conversely, the contribution of Aβ-LTMRs declined, consistent with a population-level reweighting of peripheral afferent drive.

**Fig. 5.**
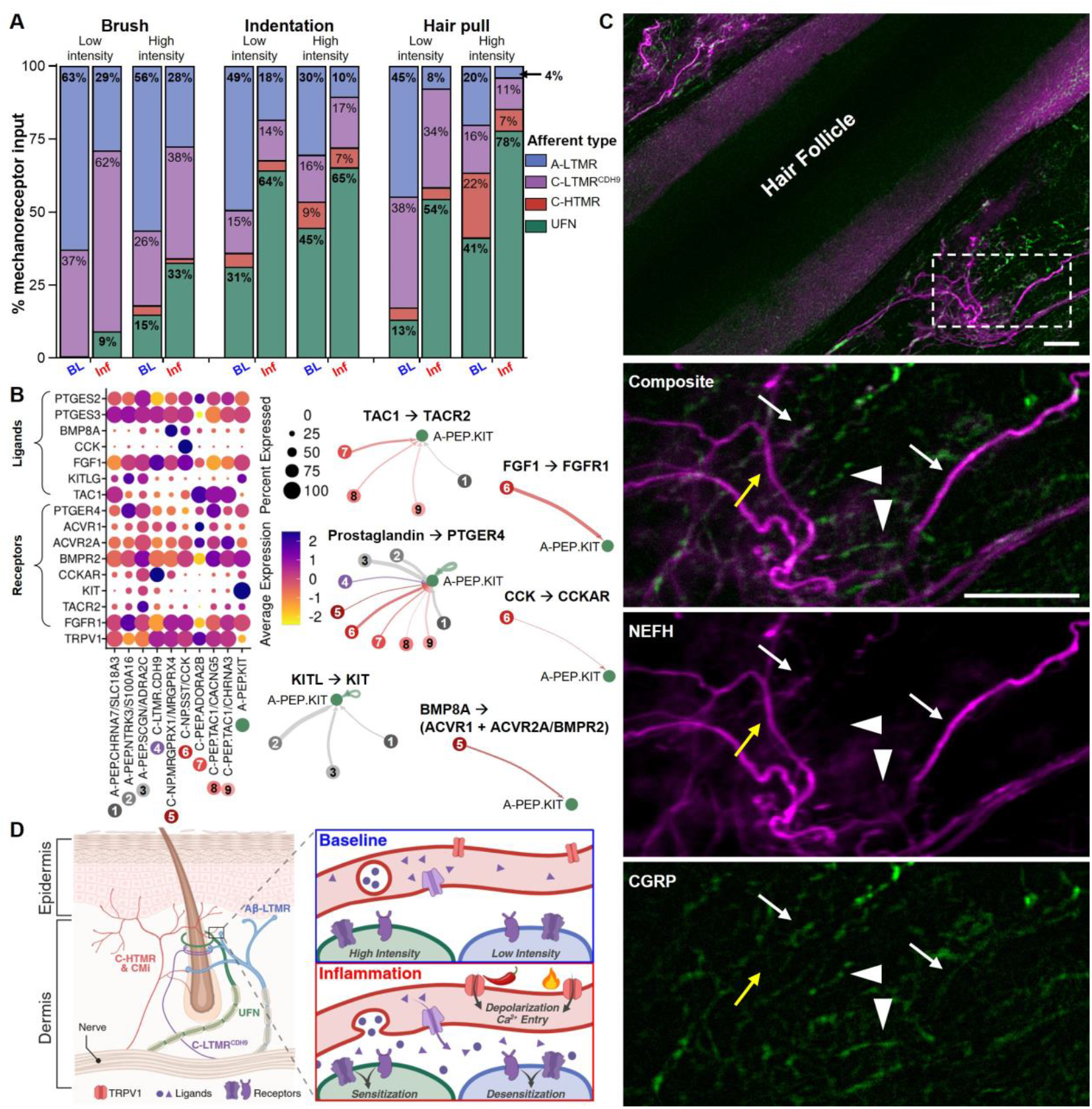
Heat-induced inflammation reweights peripheral mechanoreceptor drive via local signaling mechanisms. (**A**) Relative contribution of Aβ-LTMRs (Field + SA), C-LTMR^CDH9^, C-HTMRs, and UFNs to mechanical encoding at baseline and after heat-induced inflammation. The contribution of each afferent class to a stimulus modality (subdivided into low/high intensity - brush: soft vs coarse; indentation: 4–60 mN vs 100–3000 mN; hair pull: 10–50 mN vs 100–400 mN) was calculated from spike counts, normalized by the number of recorded units in that class, as a fraction of total spike counts for all afferent classes. Heat-induced inflammation increased the proportional UFN drive (green) and reduced Aβ-LTMR drive (blue) across modalities and intensities. (**B**) Predicted ligand-receptor (L-R) interactions from TRPV1^+^ DRG populations to UFN^A-PEP.KIT^ neurons. Left: Dot plot shows gene expression level (heatmap) and percent of cells expressing (dot size) for selected ligands and receptors. Right: Circle plots highlight predicted L-R pathways with established nociceptive roles. Lines illustrate inferred interaction strength (thickness) and direction of signaling (arrowheads; towards UFN^A-PEP.KIT^). (**C**) Spatial organization of cutaneous afferent subtypes in human hairy skin. Representative confocal images show immunofluorescence of CGRP and NEFH. Afferent fibers were identified by co-expression patterns: C-fibers are CGRP^+^/NEFH^-^, UFN are CGRP^+^/NEFH^+^, and Aβ-LTMRs NEFH^+^/CGRP^−^. Large composite image shows dense innervation of the perifollicular region. Higher-magnification views highlight the close proximity of CGRP^+^ C-fibers (white arrowheads) to NEFH^+^/CGRP^−^ Aβ-LTMRs (yellow arrow) and to NEFH^+^/CGRP^+^ UFNs (white arrows), suggesting potential anatomical substrates for local sensory interaction. Scale bars 50 µm. (**D**) Working model illustrating the local signaling mechanisms by which TRPV1^+^ C-fibers mediate the reweighting of UFN and Aβ-LTMR afferents during heat/capsaicin-induced inflammation. During inflammation, TRPV1 channels on C-fibers (red terminal) open enabling Ca^2+^-dependent synthesis and release of inflammatory mediators (FGF, prostaglandins or tachykinins). Whether these ligands act on neighboring TRPV1^−^ Aβ-afferents to produce *sensitization* (as for UFNs) or *desensitization* (as for Aβ-LTMRs) depends on the compliment of expressed receptors.

To identify the candidate signaling pathways capable of supporting afferent-specific reweighting during inflammation, we examined ligand-receptor interactions using published and newly generated scRNA-seq datasets from molecularly defined human DRG neuron populations (*10*) (Fig. 5B, Fig. S9, and Table S1). We found several mediators associated with inflammatory signaling – including prostaglandins, bone morphogenetic proteins, fibroblast growth factors, tachykinins, and KIT ligand – in TRPV1-expressing DRG neuron populations. The UFN^A-PEP.KIT^ neurons expressed the cognate receptors (PTGER3/4, ACVR1, FGFR1, TACR2, and KIT), indicating their capacity to respond to locally released inflammatory mediators. These analyses indicated potential molecular pathways through which TRPV1-mediated inflammation could influence neighboring TRPV1-negative UFN^A-PEP.KIT^ afferents within inflamed skin. Because transcriptomic identities corresponding to Field- and SA-LTMRs remain incompletely resolved in current human DRG datasets, similar ligand-receptor analyses for these populations were not feasible.

We assessed the anatomical organization of sensory afferents in human hairy skin by performing double immunofluorescence staining for CGRP and NEFH (Fig. 5C). This labeling distinguished CGRP^+^ C-fibers, NEFH^+^ A-LTMRs lacking CGRP expression, and UFNs co-expressing CGRP and NEFH. Confocal imaging revealed a co-localization of these afferent subtypes within the perifollicular region. The CGRP^+^ C-fibers were frequently observed in close proximity to NEFH^+^ Aβ-LTMRs and NEFH^+^/CGRP^+^ UFNs (Fig. 5C). The close spatial intermingling of these terminals provides an anatomical substrate through which activity in C-nociceptive fibers could rapidly influence neighboring myelinated afferents during inflammation.

Integrating our electrophysiological, transcriptomic, and anatomical observations, we outlined a local signaling framework that links TRPV1 activation of human peptidergic afferents to crosstalk with neighboring TRPV1^−^ Aβ afferents in the skin, leading to changes in mechanoreceptor coding (Fig. 5D). Under baseline conditions, UFNs preferentially encode high-intensity mechanical input, whereas Aβ-LTMRs dominate low-intensity encoding. During inflammation, UFNs showed robust sensitization, consistent with ligand–receptor analyses identifying potential inflammatory mediators released by TRPV1-expressing neurons and receptors expressed by UFNs. In parallel, our functional data demonstrated a marked attenuation of Aβ-LTMR input during inflammation, although the mechanisms underlying this suppression remain unresolved. Together, these processes result in a redistribution of peripheral mechanoreceptor contributions toward nociceptive signaling during inflammation.

## Discussion

Our findings show that acute heat-induced inflammation produces a rapid recalibration of myelinated afferent signaling, which is mediated indirectly by TRPV1 activation in other sensory afferents. This rebalancing is marked by a gain of function in UFNs and attenuated function in selective Aβ-LTMRs, indicating an early and dynamic peripheral contribution to pain hypersensitivity that shapes the afferent drive entering central circuits.

The gain-of-function response in UFNs is not intrinsic to the cell type: We draw this conclusion because these neurons remained heat- and capsaicin-insensitive before and after inflammation, consistent with their lack of TRPV1 expression. Instead, our data supports an indirect mechanism mediated by TRPV1^+^ fibers, echoing Lewis’s early proposal of neurogenic inflammation (*1*). In humans, “silent” CMi nociceptors are preferentially recruited by capsaicin (*17*) and constitute the principal drivers of neurogenic flare (*12*). Gating of TRPV1 by capsaicin or heat results in impulses that travel antidromically down collateral branches (the axon reflex) and cause the release of neuropeptides such as substance P, neurokinin A, and CGRP from peripheral nerve endings. We propose that the release of inflammatory mediators, including neuropeptides and growth factors, from peripheral TRPV1-positive afferents sensitizes adjacent UFNs.

Analysis of the human DRG single-cell transcriptomics revealed several candidate TRPV1^+^ afferent fiber interactions that could drive the sensitization of UFN^A-PEP.KIT^. These include several ligand/receptors that have previously been implicated in pain: PGE2, FGF1 (*18, 19*), BMP8A (*20, 21*), CCK (*22*), and KIT (*23*) produced by TRPV1^+^ afferent fibers in interactions with their cognate receptors expressed by UFN^A-PEP.KIT^. However, the most notable interaction was that between TAC1 and TACR2. TAC1 encodes substance P and neurokinin A (among other related peptides), which interact with TACR1 and TACR2 receptors, respectively (*24*). Substance P and neurokinin A, along with CGRP, cause strong arteriolar vasodilation (i.e., the flare), and intracutaneous injection of substance P, is accompanied by a painful sensation (*25*). Previous work in mice showing that TACR2 activation increases sensory neuron excitability (*26*) and that tachykinin expression from the TAC1 gene is required for A-fiber nociceptor sensitization in nerve-injury models (*27*) supports a paracrine mechanism for inter-afferent modulation. Non-neuronal inflammatory sources set into motion by the axon reflex (*28, 29*) may further contribute to this local signaling milieu.

These types of cross-afferent interactions are anatomically plausible. UFN^A-PEP.KIT^ endings are closely associated with human hair follicles (*8, 9*) and likely form circumferential endings akin to their mouse counterparts (*30*). Human hair follicles are also densely innervated by TRPV1^+^ C-nociceptors that drive flare responses. This close spatial arrangement provides a substrate for rapid, spatially confined inter-afferent signaling during neurogenic inflammation.

Sensitized UFNs exhibited lowered mechanical thresholds and recruitment of previously silent receptive-field branches. This expansion provides a peripheral neural correlate consistent with mechanical allodynia, thereby allowing innocuous forces to activate nociceptive signaling across enlarged skin territories. Previous work has demonstrated that human C-fiber nociceptors possess mechano-insensitive but electrically excitable subfields that can be unmasked using mustard oil or capsaicin (*31*). Our findings extend this principle to UFNs and reveal a markedly greater relative expansion of receptive fields following inflammation. Importantly, unlike C-fibers – whose newly unmasked regions remain in the noxious threshold – UFN thresholds after inflammation fall within the tactile range, enabling normally innocuous stimulation to engage nociceptive pathways across both original and newly recruited territories. This combination of tactile-range thresholds and large-scale receptive-field expansion positions UFNs as major drivers of widespread punctate allodynia and hyperalgesia.

The finding that capsaicin applied outside the original receptive field induces a comparable gain of function reinforces a distributed peripheral mechanism: TRPV1^+^ C-fiber activity can quickly modulate neighboring Aβ-afferent excitability. Classical explanations of secondary hyperalgesia emphasize central sensitization supported by ongoing peripheral drive, the tuning of which is presumed to remain stable. Our data prompt a reappraisal of this assumption, showing that during acute inflammation UFNs rapidly acquire reduced mechanical thresholds and expanded receptive fields via recruitment of silent branches. As a result, mechanical stimulation over a broader skin area, and at what were previously subthreshold intensities, increases nociceptive input at its peripheral origin.

Recent mouse work using in vivo calcium imaging of DRG neurons argues that tactile allodynia can arise without qualitative changes in peripheral encoding (*32*), with spontaneous nociceptor activity interacting with preserved A-LTMR input to bias central interpretation. Our data, by recording action potentials of peripheral afferents, extend this framework by showing that peripheral input is not uniformly preserved in humans. Importantly, Aβ-LTMRs become functionally down-weighted, whereas UFNs gain function. This reweighting of fast-conducting A-fiber channels is particularly consequential given their rapid conduction, extensive central arborizations (*33*) and ability to modulate protective reflex circuits (*34*). From a gate-control perspective, simultaneous increases in nociceptive drive and decreases in tactile input constitute a bidirectional shift that favors pain transmission. This pattern parallels the reduction in Aβ-LTMR mechanical sensitivity associated with fibromyalgia (*35*) and suggests that tactile down-weighting may be a shared feature of acute inflammatory and chronic pain states.

CMi nociceptors have been implicated in neuropathic pain and fibromyalgia (*36, 37*), where their contribution is often attributed to gain of function or spontaneous activity. In contrast, the UFNs in our study exhibited gain of function without spontaneous firing and in response to TRPV1-mediated inflammatory signaling despite their own lack of TRPV1. The CMi nociceptors, by comparison, displayed prolonged after-discharge following heat-induced inflammation. This suggests a distinct modulatory role in which extended C-fiber activity broadens the temporal window for influencing neighboring myelinated afferents through local crosstalk mechanisms (Fig. 5B-D).

At the spinal level, TRPV1^+^ and TRPV1^−^ afferent terminals can be found in close proximity, but this arrangement is unlikely to account for the rapid peripheral changes in UFNs and Aβ-LTMRs observed here. The timescale and selectivity of these effects are difficult to reconcile with known spinal anatomy or with centrally mediated (antidromic) modulation of peripheral afferent activity. Instead, inflammatory hypersensitivity at the spinal level is associated with shifts in excitatory-inhibitory ensemble activity, including reduced inhibitory interneuron recruitment and increased excitatory drive, which are changes that emerge over days to weeks (*38*). Viewed alongside our findings, this suggests that similar reweighting principles may operate across the somatosensory system, with a redistribution of sensory drive occurring rapidly (within minutes) in the periphery before longer-timescale adaptations unfold centrally.

Finally, our data challenge rigid labeled-line interpretations. Although peripheral neurons show modality selectivity under baseline conditions, we demonstrate a profound state dependence, as acute injury induces functional polymodality in UFNs, including de novo sensitivity to innocuous touch and mild cooling. This plasticity complements the distributed processing observed in central circuits (*39*), underscoring that the boundary between painful and nonpainful is dynamically reconstructed across multiple levels of the system.

Our identification of UFNs as major contributors to widespread hyperalgesia expands the canonical C-fiber–centric view of human pain mechanisms. This finding may prove to be of value to clinicians and to patients living with hyperalgesia, because these patients experience the limited efficacy of currently dispensed centrally acting analgesics and the dangers of addictive opioid therapies. Targeting peripheral gain- and loss-of-function mechanisms within molecularly defined A-fiber populations may provide novel non-opioid therapeutic avenues. Therefore, whether similar peripheral reweighting contributes to chronic pain remains an important question for future studies. More broadly, our results highlight the importance and power of real-time interrogation of human sensory afferents combined with transcriptomic insight, as this strategy can reveal rapid, state-dependent shifts in encoding that may otherwise be invisible when using approaches lacking temporal resolution.

Several limitations should be considered when interpreting these findings. First, although our data support a model in which TRPV1^+^ afferents drive local inter-afferent signaling in the skin, TRPV1 expression is not restricted to sensory neurons (*11*). While the rapid, afferent-class-specific changes observed here argue for a neural mechanism, contributions from non-neuronal cells are not precluded. Second, electrophysiological recordings, psychophysical testing, and flare measurements were, necessarily, performed in separate participant cohorts, limiting direct within-subject correlations. Third, transcriptomic mapping of human afferents remains incomplete, particularly for Aβ Field- and SA-LTMR populations, limiting molecular inference for these classes. Finally, our study focuses on acute heat-induced inflammation; whether similar peripheral reweighting operates in chronic pain states remains to be determined.

## Acknowledgments

We acknowledge the Microscopy Core Facility at the Faculty of Medicine and Health Sciences, Linköping University and V. Loitto for providing assistance in confocal imaging. We appreciate the help of S. Amezcua with human skin biopsies and D. Persson and L. Medling with participant recruitment. We thank U. Alam for access to resources and regulatory support for the latency-tracking experiments. We also extend our thanks to W. Moore and K. Ng for assisting in some of the microneurography data collection.

## Author contributions

Conceptualization: SSN, HO, OB

Formal analysis: OB, SHL, AM, JK, HM, AB, HY, IS, JJ, AGM

Funding acquisition: SSN, HO, PE, WL, AGM

Investigation: OB, SHL, AM, AGM, SSN, HM, JJ

Project administration: OB, SHL, SSN

Supervision: RS, ML, WL, AGM, PE, HO, SSN

Visualization: OB, SHL, HM, AB, IS

Writing – original draft: OB, SSN, HO

Writing – review & editing: all authors

## Data, code, and materials availability

All data supporting the findings of this study are available in the manuscript and Supplementary Materials. Upon acceptance, source data, analysis code, and any related materials will be deposited in a publicly accessible repository with a citable DOI.

## Supplementary Materials

Materials and Methods

Figs. S1 to S9

Table S1

References (7-9, 16, 40-57)

