## Supplementary Materials for "Rapid peripheral reprogramming of myelinated afferents drives human hyperalgesia"

**The PDF file includes:**

Materials and Methods
Figs. S1 to S9
Table S1

### Materials and Methods

#### 1. Human participants

Heat-induced inflammation was assessed using psychophysical, neurophysiological, and histological approaches. Psychophysical testing was performed in 18 healthy volunteers (8 males, 10 females; 19–62 years). Skin imaging was conducted in a separate cohort of 10 healthy controls (7 males, 3 females; 30–40 years). Ultrasound-guided microneurography was carried out in 75 participants (41 males, 34 females; 21–60 years). For immunostaining of cutaneous afferents, distal leg skin biopsies were obtained from an additional six healthy controls (5 males, 1 female; 32–45 years). Participants were recruited through university and social media advertisements. The exclusion criteria comprised neurological or musculoskeletal disorders, skin diseases, diabetes, and recent use of analgesic or psychoactive medication. All participants provided written informed consent before the start of the experiment. All procedures were approved by the Swedish Ethical Review Authority (2020-04426 (amended 2024-00772-02), 2016-43331) and Southwest-Frenchay Research Ethics Committee of University of Liverpool (20/SW/0138) and were conducted in accordance with the Declaration of Helsinki.

#### 2. Psychophysical assessment

A 1.2 × 1.8 cm rectangular test area was marked on the dorsolateral forearm, corresponding to Contacts 4 and 5 of a Peltier thermode (QST.Lab, Strasbourg, France). All assessments were performed within this homotopic zone, except for mapping of secondary (heterotopic) hyperalgesia. Participants were blinded to the stimulation area. Pain intensity was rated using a numeric rating scale (NRS; 0–100). The full protocol required approximately 12 min. Mechanical detection thresholds were assessed using von Frey monofilaments (0.25–512 mN; OptiHair2, MRC Systems, Germany), applied perpendicularly until buckling, with five subthreshold and five suprathreshold values obtained. Mechanical pain sensitivity was evaluated using nylon monofilaments (4–3000 mN; Aesthesio, Bioseb, France) applied in pseudo-random order (~1 s per stimulus), with pain ratings recorded after each stimulus. Secondary hyperalgesia was mapped using a 128 mN pinprick stimulator (MRC Systems, Germany) applied along eight radially oriented tracks toward the test site in ~1 cm increments. Participants indicated a clear increase in sharpness or pain, and corresponding distances (proximal–distal, medial–lateral, and diagonal) were measured. Thermal detection thresholds, thermal sensory limen, and thermal pain thresholds were assessed using a contact thermode (T09, QST.Lab, France) positioned over the test area (baseline 32 °C; ramp rate 1 °C/s), with two repetitions per measure. Thermal pain ratings were obtained using ramp-and-hold stimuli (4 °C/s ramp, 5 s hold) to 0 °C and 50 °C, with two repetitions per temperature. Following baseline testing, heat-induced inflammation was evoked using twelve ramp-and-hold heat stimuli (30–50 °C; 100 °C/s ramps; 10.2 s duration) delivered via two adjacent stimulation zones (each 7.4 × 12.2 mm) within the test area. Pain ratings were collected after each trial. The full psychophysical protocol was repeated immediately and 20 min after heat-induced inflammation.

#### 3. Spatial frequency domain imaging (SFDI)

Heat-induced inflammation was assessed by measuring changes in tissue blood volume using an in-house quantitative multi-spectral Spatial Frequency Domain Imaging (SFDI) system (*40*). The hand was immobilized using a vacuum pillow, such that the relatively flat surface of dorsal hand (indicated in Fig. S1) was centered in the camera's field of view. Peltier thermode placement was marked on the skin (~2 cm bracket). The test area (proximal to this bracket) included the area of direct heating (ROI; 7.4 x 12.2 mm) from Contact 5 of the thermode and several cm of surrounding skin. Measurements were calibrated with a silicone phantom of known optical properties. Images were acquired at 520, 536, 556, and 626 nm at ~1 min intervals before (baseline) and for 30 min after heat-induced inflammation (as described above). By illuminating tissue with a sequence of sinusoidal intensity patterns, optical absorption can be differentiated and quantified from the effects of volumetric light scattering within tissue. Optical absorption data were used to estimate oxyhemoglobin, deoxyhemoglobin (*41*), and melanin concentrations by least-squares fitting in MATLAB R2024a. An epidermal thickness of 75 μm was assumed (*42*), with melanin confined to the epidermis, a layered model of light transport was employed (*43*) and hemoglobin localized to blood vessels (mean radius 100 μm) (*44*). This layered model isolation of dermal hemoglobin underlying epidermal melanin has been validated in vivo across different skin pigmentations (*45, 46*). Total blood volume was calculated as the sum of oxy- and deoxyhemoglobin and expressed as tissue volume percentage. Flare extension was quantified at 1, 5, and 30 min after heat-induced inflammation as the distance from the center of the heated area to the 50% isocontour between baseline and post-inflammation blood volume within the ROI. One participant was excluded due to atypical baseline blood volume, yielding data from nine participants. To exclude residual heat as the sole contributor to blood-volume changes, thermal imaging was performed at baseline and up to 5 min after heat-induced inflammation in six volunteers. Each temperature measurement was relative to a nearby non-skin constant temperature surface and then change from baseline was calculated.

**4. In Vivo electrophysiological recordings of human sensory afferents**

Single unit axonal recordings (microneurography) were obtained from the posterior antebrachial cutaneous, radial, or superficial peroneal nerves in awake participants seated comfortably with the tested limb supported. Room temperature was maintained at 22 °C, and participants acclimated before testing. Microneurography was performed using established protocol (*7, 8*). Under real-time ultrasound guidance (LOGIQ P9; GE Healthcare), an insulated tungsten microelectrode (FHC Inc.) was inserted into the target nerve, with a nearby uninsulated reference electrode. Neural signals were amplified using a high impedance headstage (MLT185) and low noise amplifier (FE185 Neuro Amp EX; ADInstruments). Neural activity was sampled at 20 kHz, and acquired using a PowerLab 16/35 system (hardware PL3516/P; ADInstruments) and LabChart Pro v8.1.5 (ADInstruments, Oxford, UK). Upon entering a fascicle and once a stable intraneural position was achieved, two protocols were used to search for cutaneous mechano-sensitive and -insensitive afferents.

- **Protocol 1: Recording mechano-sensitive afferents**

Mechanosensitive afferents were identified using gentle brushing and classified as low or high threshold mechanoreceptors according to standard criteria (*7, 16, 47*). Mechanical thresholds were determined using Semmes–Weinstein monofilaments. Receptive-field (RF) mapping was performed using forces at least four times each unit's pre-inflammation mechanical threshold. The same force was applied before and after inflammation. Following inflammation, we confirmed that the expanded RF responded to the updated mechanical threshold. Low and high intensity brushing stimuli were applied at ~3 cm/s (three repetitions), and force–response relationships were assessed using monofilaments delivering 4–3000 mN. Individual hairs within the RF were stimulated using a handheld hairpuller as previously described (*9*). Thermal sensitivity was tested, using a single contact of the Peltier thermode (7.4 × 12.2 mm; QST.Lab), with ramp-and-hold (4 °C/s ramp; 5 s hold) cooling (30-0 °C) and heating (30-50 °C) stimuli (three repetitions each). Heat-induced inflammation was elicited using the same protocol as in the psychophysical experiments. In microneurography, heating was applied via a single 7.4 × 12.2 mm thermode contact positioned over the mapped receptive field, which was sufficient given the spatially confined receptive fields of individual units. Chemically-induced inflammation was tested in separate experiments by topical application of capsaicin (0.075%) on/near the RF for 5 min, followed by removal of residual compound. Units were continuously monitored for spontaneous activity, and all test stimuli were repeated after inflammation. All responses were confined to the defined RF and reproducible across repeated trials. Conduction velocity (CV) was estimated using latencies to electrically or mechanically evoked spikes and measurement of the distance between the RF and recording electrode.

- **Protocol 2: Recording mechano-insensitive C-afferents**

Mechano-insensitive C-afferents (CMi) were identified using transcutaneous electrical stimulation (2–3 mm steel electrode tip). After C-fibers were detected, sterile electro-acupuncture needles (Harmony Medical, 13 mm × 0.22 mm) were inserted intradermally immediately distal to the receptive field and advanced to within 5 mm of the electrically defined site. Electrical pulses (0.5 ms; 0–20 mA; Digitimer DS7) were delivered at 0.125–0.5 Hz at ~2× threshold, within participant tolerance. The stimulation protocol comprised 2–6 min at 0.25 Hz, a 2-min pause, followed by activity-dependent slowing (ADS): 20 stimuli at 0.125 Hz, 20 at 0.25 Hz, 30 at 0.5 Hz, and a final continuous 0.25 Hz block (*48*). Mechanical responsiveness was assessed during 0.25 Hz stimulation using the marking technique (*49*) and monofilaments (64–512 mN) were applied within a ~1 cm region surrounding the electrical stimulation site. For thermal testing the QST thermode was placed at the electrical stimulation site, and baseline data was collected at 30 °C for ≥30 s followed by the heat-induced inflammation protocol above; 0.25 Hz stimulation continued for up to 25 min. The following parameters were calculated: *(i)* CV in m/s (distance between the recording electrode and electrical stimulation site divided by the unconditioned latency after the 3 minute pause), *(ii)* ADS in the stimulation protocols (final latency during 0.5 Hz expressed as a percentage of the initial latency at the start of the protocol), *(iii)* Percentage slowing of CV change to 0.25 Hz stimulation from rest (after a 3 minutes pause), and *(iv)* Recovery of CV after ADS stimulation protocol was calculated as the absolute change in latency from the final latency to stimulation at 0.5 Hz to the latency after 10 further stimulations at 0.25 Hz expressed as a percentage of the final latency at 0.5 Hz. Units were classified as CM or CMi according to ADS-based criteria (*48*).

Neural data from Protocol 1 were analyzed using LabChart and Spike2, whereas data from Protocol 2 were processed using SpikeSpy. Spikes were detected using a ≥2:1 signal to noise ratio and confirmed by waveform template matching.

**5. Skin biopsy and immunostaining**

Human skin biopsies were obtained from the distal lower leg, approximately 10 cm proximal to the lateral malleolus, using a 3-mm circular punch under local anesthesia in accordance with US and European clinical guidelines (*50*). Tissue samples were fixed in 4% paraformaldehyde (PFA; Polysciences Europe GmbH) at 4 °C overnight, followed by cryoprotection in 30% sucrose at 4 °C for an additional 24 h. Biopsies were then embedded in optimum cutting temperature (OCT) compound, and 100-µm sections were prepared using a cryostat (CM1950, Leica Biosystems Inc.). Free-floating sections were processed for immunohistochemistry, stained, and optically cleared with RapiClear 1.47 (Sunjin Lab, Hsinchu City, Taiwan) according to the manufacturer's instructions. Triple immunolabelling was performed using the following primary antibodies: Rabbit anti-PGP 9.5 (1:1000; ThermoFisher), Chicken anti-α-NEFH (1:2000; ThermoFisher), and Guinea pig anti-CGRP (1:250; Synaptic Systems). Sections were incubated with primary antibodies for 5 days at 4 °C. Corresponding Alexa Fluor–conjugated secondary antibodies (Alexa

488, 568, or 647 IgG; 1:200; Invitrogen) were applied for 3 days at 4 °C. Skin sections were cleared with RapiClear 1.47 at RT overnight and then mounted on slide with fresh RapiClear reagent. Images were acquired using a spinning disk confocal microscope (Nikon Eclipse Ji) equipped with a 20× objective.

**6. Ligand-Receptor pair analysis**

Ligand–receptor (L–R) interactions among human dorsal root ganglion (DRG) neuron subtypes were inferred using CellChat (v2.1) following the recommended workflow for single cell RNA seq datasets as described in the developers' online tutorial (https://github.com/jinworks/CellChat) (*51*). As input, we used a subset of the Bhuiyan et al. (*10*) dataset restricted to neuronal populations exhibiting high TRPV1 expression, together with the A PEP.KIT neuron type. To focus on paracrine and small molecule communication, the analysis was limited to the "Secreted Signaling" and "Non-protein Signaling" categories within the CellChat database. All TRPV1 expressing neuron types were designated as signal sending ("source") populations, and A PEP.KIT neurons were defined as the signal receiving ("target") population. CellChat was used to compute the probability of L–R interactions and to statistically evaluate communication likelihoods across all source–target pairs. Using a significance threshold of $p < 0.05$, we identified 462 putative L–R interactions between individual TRPV1+ neuron types and A-PEP.KIT neurons. These initial interactions were subsequently curated by filtering based on interaction strength and biological plausibility, incorporating established knowledge of sensory neuron subtype function and innervation patterns within the skin.

**7. Data analysis and visualization**

**7.1. Heatmap**

For heatmap visualization, a total of 65 units were analyzed (7 UFN$^{A\text{-}PEP.KIT+}$, 13 UFN$^{KIT}$, 13 C-HTMR, 13 Field-LTMR, 13 SA-LTMR, and 6 C-LTMR$^{CDH9}$ units). Each unit was tested with different stimulus modalities (brush, mechanical indentation, hair pull, heating, and cooling) at varying intensity levels under two experimental conditions: Baseline and Inflammation. For the hair-pull stimulus modality, the receptive field of some units (6 UFN$^{KIT-}$, 8 C-HTMR, 3 Field-LTMR, 9 SA-LTMR, and 4 C-LTMR$^{CDH9}$ units) lacked a hair follicle in the receptive field, and therefore, we were not able to record a response. For all stimulus modalities, the spike count was used to quantify the neural response. Data were normalized for each stimulus modality of each afferent type and scaled to 100% using the following formula:

$$x' = x / \max(x) * 100\%$$

Heatmap displays normalized response values (0–100%) with units as rows and stimulus modalities as columns. The color scale reflects the magnitude of the normalized response. Heatmap was generated using R (version 4.3.2). The following packages were used for data pre-processing and visualization: readxl (*52*), tidyverse (*53*), ComplexHeatmap (*54*), circlize (*54*), and pBrackets (*55*).

**7.2. UMAP visualization**

Data used for the UMAP visualization included spike count data from all stimulus modalities and peak frequency data from brush, mechanical indentation, and hair pull stimulus modalities. No normalization was applied. Hair pull missing data were imputed using the median calculated separately for each combination of pulling force, afferent type, and experimental condition. The data were split into two categories: pain-sensing neurons and touch-sensing neurons. The pain-sensing group included UFN$^{A\text{-}PEP.KIT+}$, UFN$^{KIT-}$, and C-HTMR units, whereas the touch-sensing group included Field-LTMR, SA-LTMR, and C-LTMR$^{CDH9}$ units. To reduce the dimensionality, Principal Component Analysis (PCA) was first performed using the base R function prcomp with centering (center = TRUE) and scaling (scale. = TRUE) enabled. These settings normalize the data to have a mean of zero and a standard deviation of one, thus ensuring that all features contribute equally to the PCA. Next, the first 15 principal components were selected. Uniform Manifold Approximation and Projection (UMAP) was performed on these selected principal components using the umap package version 0.2.10.0 with default configuration settings (*56*). UMAP was applied to further reduce the dimensionality in a nonlinear manner, preserving local data structure and relationships that may be missed by linear methods such as PCA, thereby improving visualization and clustering of our complex dataset.

#### 7.3. Mechanical input analysis

To determine how acute inflammation reshapes the mechanical sensory input, we quantified the relative contribution of each afferent type to the total sensory input. Spike counts were first normalized to account for differences in the number of units across afferent types. Specifically, the mean spike count for each afferent type (UFN, C-HTMR, A-LTMR, and C-LTMR$^{CDH9}$) was calculated by dividing the total spike count by the number of units of that class. For each stimulus modality and intensity level, mean spike count values were expressed as a proportion of the total population, providing a measure of the "relative contribution" of each afferent type. The relative contribution of UFN input was analyzed using a Beta Regression model using the betareg package in R (*57*). The model included experimental condition, stimulus modality, and intensity level. Statistical significance was determined via maximum likelihood estimation.

#### 7.4. Statistical analysis

Statistical analyses were conducted using GraphPad Prism (v10.5; GraphPad Software, San Diego, USA). Psychophysical data with normal distributions were analyzed using repeated-measures ANOVA. Mechanical detection thresholds were log10-transformed to achieve normality, and 0.1 was added to all NRS ratings to retain zero values. To compare data collected at baseline with each timepoint post heat-induced inflammation (immediately after and 20 minutes), repeated measures ANOVA with Dunnett's multiple comparison test was used for normally distributed data (CDT, WDT, TSL, MDT) and Friedman's test (Geisser and Greenhouse correction) with Dunn's multiple comparison test for non-parametric data (CPT, HPT). Microneurography data were analyzed using paired t-tests for two-sample comparisons, and repeated-measures ANOVA for multiple comparisons. Two-way repeated-measures ANOVA followed by Tukey post hoc tests was used where appropriate, with F values reported for stimulus × inflammation interactions. Statistical significance was set at $p < 0.05$.

### Supplementary Figures

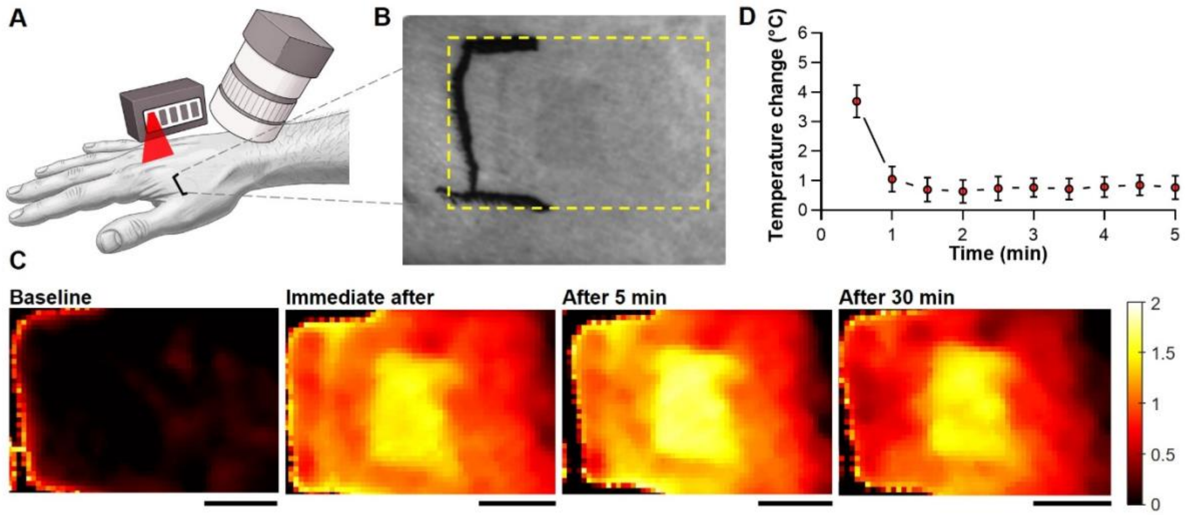


**Fig. S1. Assessment of local blood volume and skin temperature changes following heat-induced inflammation.**

(**A**) Schematic of heat-induced inflammation protocol applied to the dorsal hand using a Peltier thermode. (**B**) Photograph of the region of the hand that was assessed for skin temperature and blood volume changes. Black bracket on the skin marks location of thermode placement. The area of image analysis was proximal (right) to this. (**C**) Blood volume maps obtained by spatial frequency domain imaging (SFDI) at baseline, immediately after, 5 min, and 30 min post-heating. Color scale represents relative blood volume (%). Scale bars: 10 mm. (**D**) Changes in skin temperature (°C, relative to baseline) after heat-induced inflammation. Baseline skin temperature was measured just prior to heating, then skin temperature was assessed every 30 sec for 5 min. A brief elevation was observed at 30 sec and rapidly declined toward baseline (mean ± SEM; n = 6).

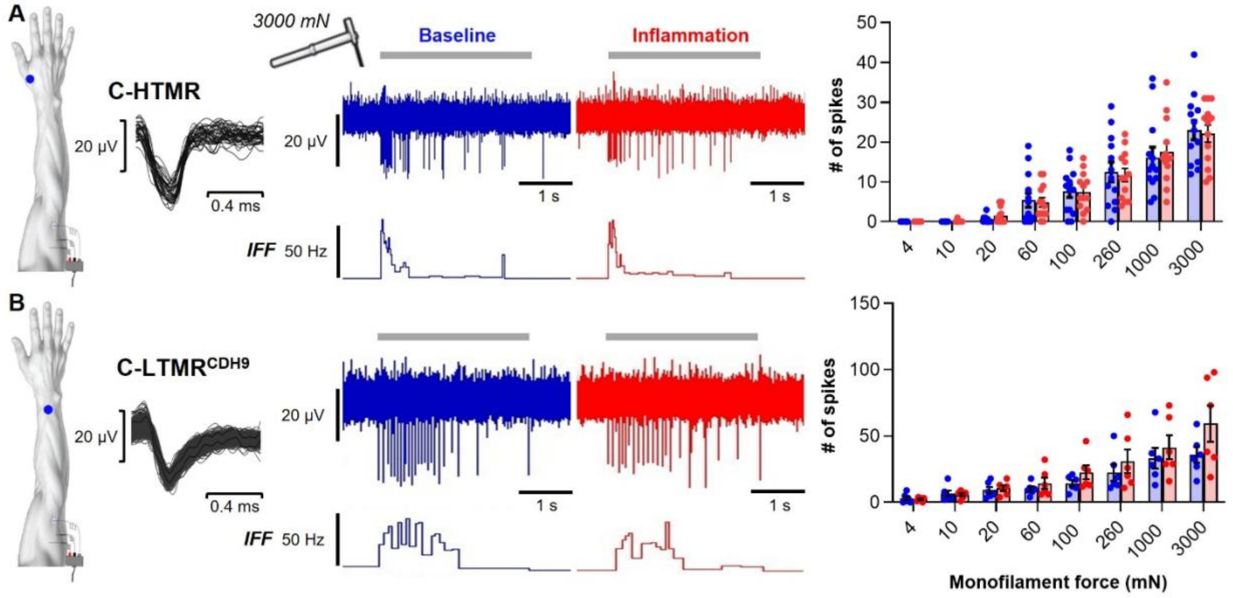


**Fig. S2. C-fiber mechanosensitivity remained stable after inflammation.**

Representative traces and quantitative analyses (plotted as in Fig. 1E-G) for C-HTMR (**A**), C-LTMR$^{CDH9}$ (**B**) during graded punctate indentation (grey bars), at baseline (blue) and following heat-induced inflammation (red). Responses to all forces were unchanged by heat-induced inflammation for both C-fiber types (C-HTMR: $F_{(7,84)} = 0.48$, $p = 0.8465$; $n = 13$; C-LTMR$^{CDH9}$: $F_{(7,35)} = 3.37$, $p = 0.1256$; $n = 6$; RM two-way ANOVA with Sidak's test).

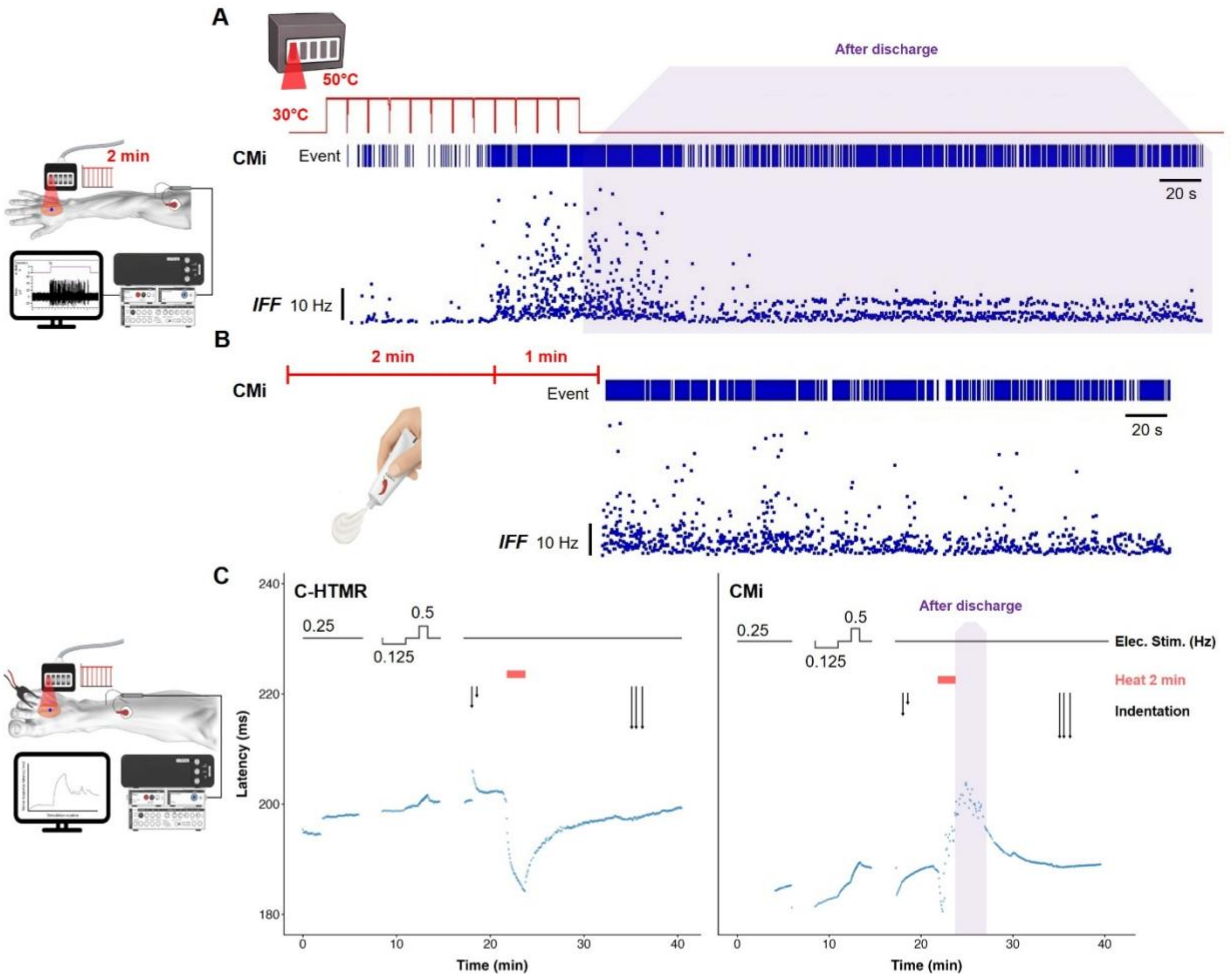


**Fig. S3. Heat/capsaicin-induced inflammation unmasked firing in CMi nociceptors.**

(**A**) CMi responses during and after the heat-induced inflammation protocol. Spike activity is shown as an event raster (top) and instantaneous firing frequency (IFF; bottom). Heat stimulation evoked delayed activation of the mechanically insensitive nociceptor, with spike activity persisting long after heat stimulation ended (after discharge; purple shading). (**B**) CMi recording following topical capsaicin application, showing robust and sustained activation. (**C**) Latency-raster plots illustrate latency changes in C-fibers during electrical stimulation of the cutaneous receptive field. The stimulation sequence consisted of baseline phase at 0.25 Hz, followed by an ascending low-frequency ADS protocol (0.125 Hz, 0.25 Hz, 0.5 Hz, and 0.25 Hz), and ending with continuous 0.25 Hz stimulation. 2–3-minute pauses separated phases. Timing of indentation indicated by arrows (short = 128 mN, medium = 256 mN, long =512 mN von Frey filaments). Left: Latency profile of a C-HTMR (CV = 0.7 m/s, ADS = 1.4%, recovery after ADS = 50%) that exhibited a marking response to a 256 mN indentation (at baseline). This afferent displayed heating-induced speeding during the heat-induced inflammation protocol and lost the mechanical marking response afterward. Right: Latency profile of a CMi (ADS = 9.5%, recovery after ADS = 11.8%) that did not exhibit mechanical marking. During heat-induced inflammation, the fiber showed an initial latency decrease consistent with heating-induced speeding, followed by a clear "marking response" characterized by an increase in latency. This increase was accompanied by mixed saw-tooth and rounded baseline deflections, indicative of spontaneous bursting and tonic firing that persisted for several minutes after heating ended (after discharge). Schematics at left illustrate Protocol 1 (top, used for CMi shown in panels A-B) and Protocol 2 (bottom, used for C-HTMR and CMi shown in panel C).

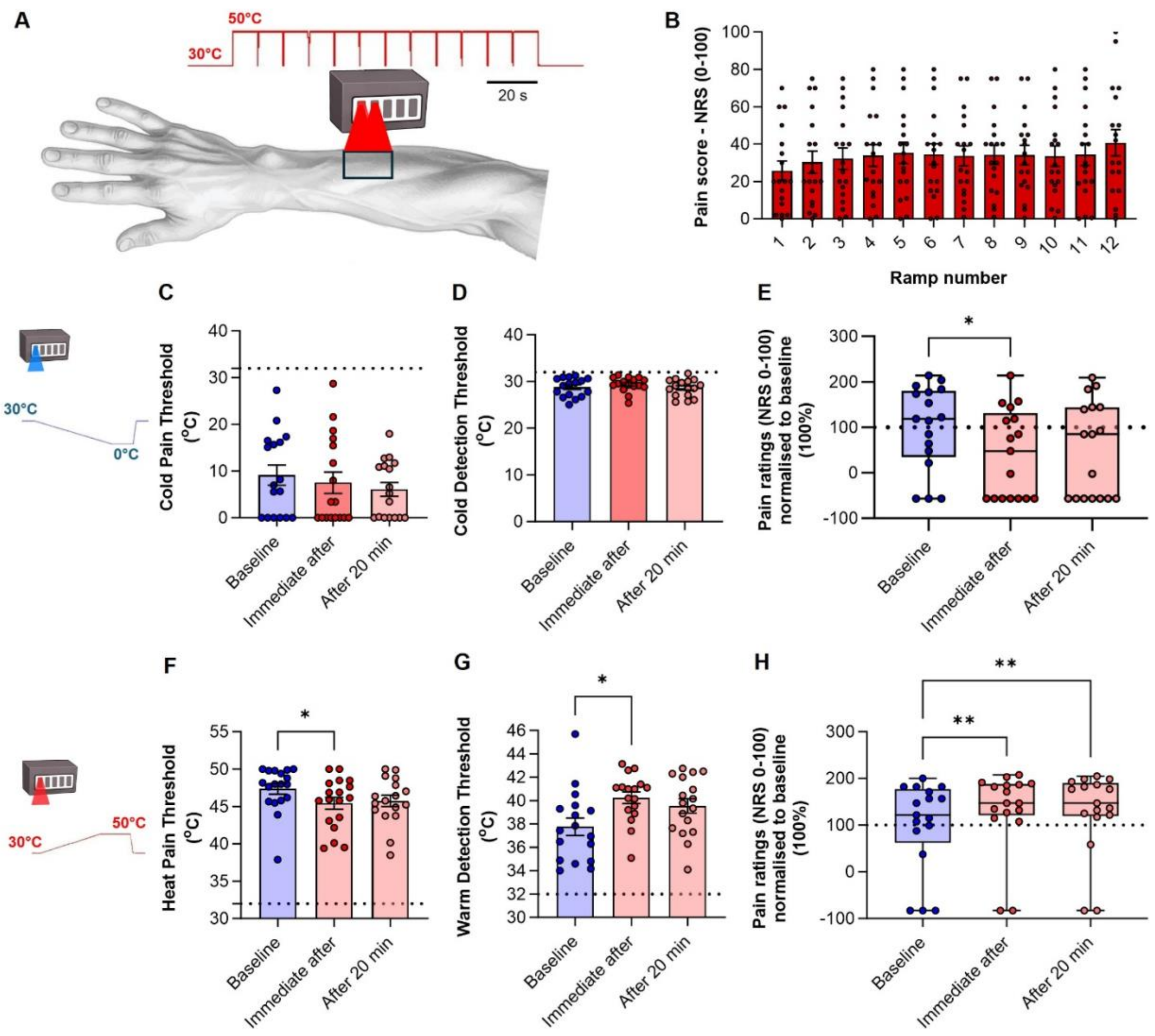


**Fig. S4. Effects of heat-induced inflammation on innocuous and noxious thermal sensitivity.** (**A**) Schematic of the primary test area used for psychophysical assessment and experimental model for heat-induced inflammation. All participants developed transient (< 24 hours) erythema in the primary area with a large proportion (88%) developing localized flare. None of the participants developed blistering. (**B**) Pain ratings during heat-induced inflammation protocol: Pain ratings (numeric rating scale (NRS) from 0 (no pain) to 100 (worst pain imaginable)) for 12 repeats of ramp-and-hold (R+H) heating to 50 °C. Median pain ratings increased from 20.0 (6.5 - 42.5 IQR) during the first R+H to 36.0 (15.0 - 66.3) during the final R+H (Wilcoxon matched-pairs signed rank test $p < 0.001$). (**C**,**F**) Thermal pain thresholds within the primary test area. Cold pain threshold (**C**) did not differ between baseline and inflammation. Heat pain threshold (**F**) was reduced immediately after inflammation ($F_{(1.669, 27.53)} = 5.12$, $p = 0.0170$; baseline: 47.3 ± 0.7 °C *vs.* immediately after: 45.4 ± 3.4 °C; $q = 2.65$, $p = 0.0307$; $n = 17$; RM one-way ANOVA with Dunnett's test). (**D**,**G**) Thermal detection thresholds within the primary test area. Cold detection threshold (**D**) was unchanged, whereas warm detection threshold (**G**) increased immediately after inflammation ($F_{(1.730, 27.69)} = 5.77$, $p = 0.0104$; baseline: 37.7 ± 0.74 °C *vs.* immediately after: 40.2 ± 2.04 °C; $q = 3.04$, $p = 0.0144$; $n = 17$; RM one-way ANOVA with Dunnett's test). (**E**,**H**) Thermal pain sensitivity. Normalized pain ratings to noxious cold (**E**) and heat (**H**) R+H stimulation (n = 18). After inflammation, cold pain ratings were reduced (baseline: median 5.5, IQR 0.8–23.3 *vs.* immediately after: median 1.0, IQR 0–7.8; $p = 0.0327$), while heat pain ratings were increased (baseline: median 11.0, IQR 3.3–40.0 *vs.* immediately after: median 20.0, IQR 11–51.3; $p = 0.0093$; Friedman's test with Dunn's multiple comparison test). Data plotted as individual values with mean ± SEM (B-D,F,G) or median ± IQR/range (E,H). *$p < 0.05$, **$p < 0.01$.

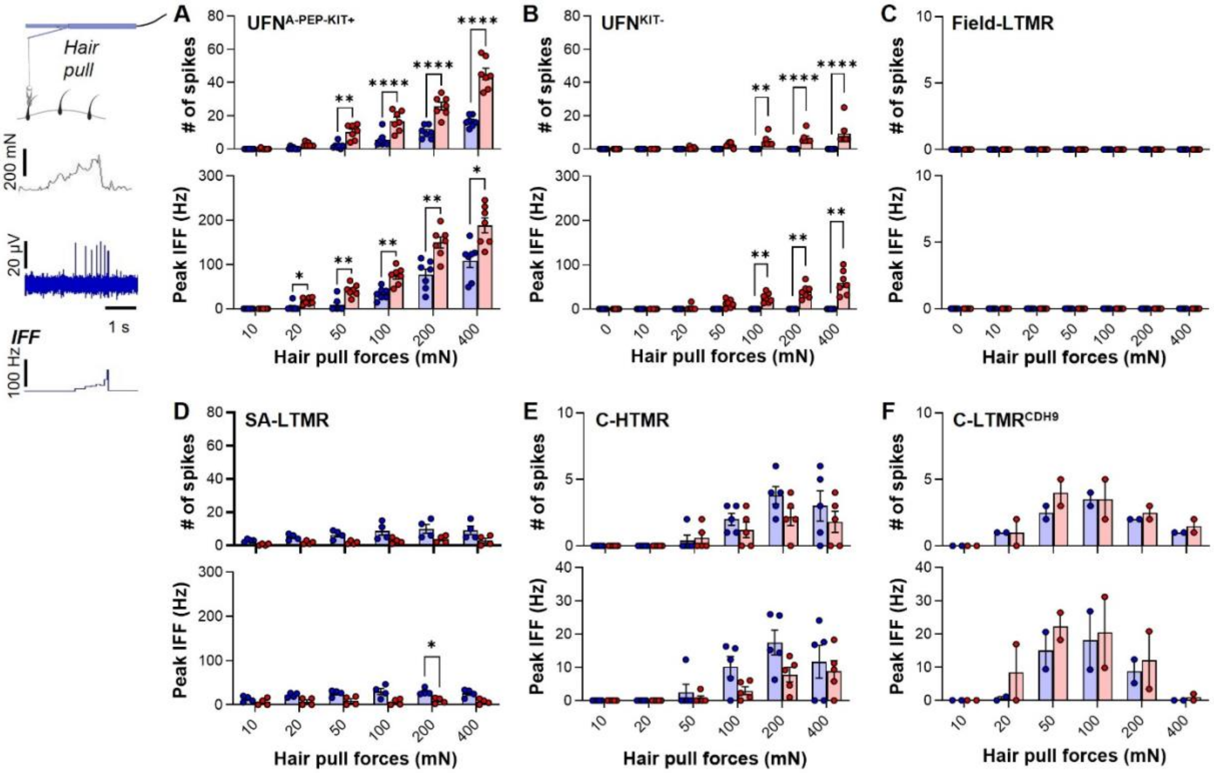


**Fig. S5. Heat-induced inflammation triggered gain-of-function responses in UFNs to hair pulling.**

Inset at top left illustrates the device used to pull a single hair while recording the force applied (~200 mN trace show at top) allowing unit responses to be correlated to the force applied. Representative traces show spike train response (middle), and instantaneous frequency (IFF; bottom). Quantitative data summarize spike counts, and peak firing frequency across force levels (10–400 mN) at baseline (blue) and following heat-induced inflammation (red). Data are presented as individual values and means ± SEM. (**A**) UFN$^{A\text{-}PEP.KIT+}$ afferents exhibited robust hair-pull sensitivity at baseline, which was potentiated by inflammation: spike counts increased across forces ($F_{(1,12)} = 87.12$, $p < 0.0001$; $n = 7$; RM two-way ANOVA), accompanied by elevated peak IFF ($F_{(1,12)} = 25.56$, $p = 0.0003$). (**B**) UFN$^{KIT-}$ afferents showed little or no baseline responsiveness but developed clear hair-pull sensitivity after inflammation (number of spikes: $F_{(1,12)} = 18.70$, $p = 0.0011$; peak IFF: $F_{(1,12)} = 36.82$, $p < 0.0001$; $n = 7$). (**C**) Field-LTMRs remained unresponsive under all testing conditions. (**D**) SA-LTMR responses were attenuated following inflammation (number of spikes: $F_{(1,6)} = 7.18$, $p = 0.0365$; peak IFF: $F_{(1,16)} = 10.86$, $p = 0.0165$; $n = 4$). (**E**) C-HTMR afferents respond to hair-pull at baseline but were not significantly altered by inflammation (number of spikes: $F_{(1,8)} = 1.93$, $p = 0.2019$; peak IFF: $F_{(1,8)} = 3.51$, $p = 0.0976$; $n = 5$). (**F**) Hair-pull responses of C-LTMR$^{CDH9}$ appear to be unaffected by inflammation ($n = 2$). Asterisks (where shown) on the force-response data indicate significant post hoc comparisons (Sidak's test). *$p < 0.05$, **$p < 0.01$, ***$p < 0.001$, ****$p < 0.0001$.

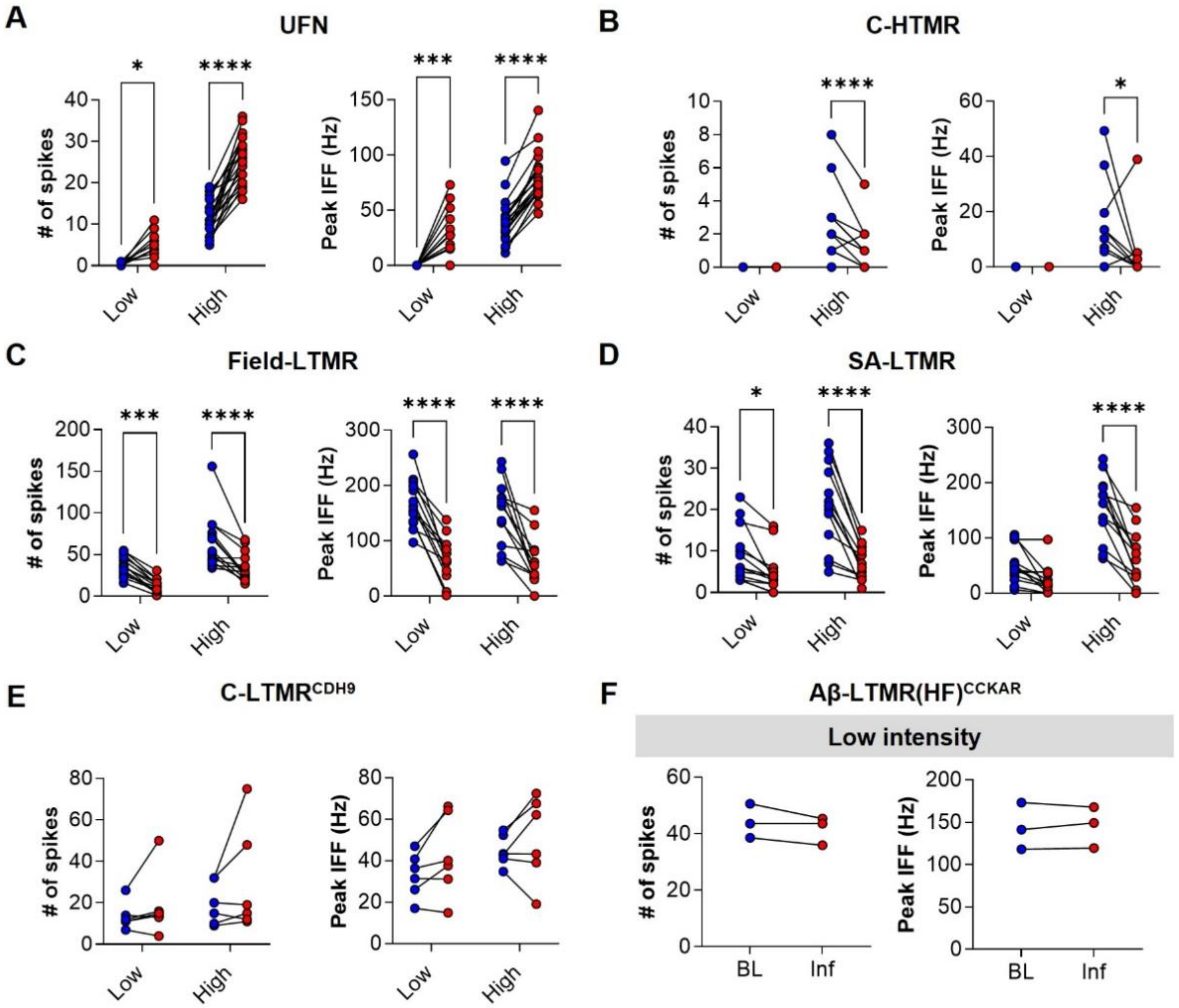


**Fig. S6. Heat-induced inflammation shifted tactile coding by potentiating UFN brush responses and dampening LTMR output.**

Summary quantifications of afferent responses during baseline (blue) and after heat-induced inflammation (red). Stimuli included low- and high-intensity stroking using soft or coarse brush, respectively. Each panel shows number of spikes (left) and peak IFF (right) for individual units in before-and-after plots. (**A**) UFN afferents exhibited robust inflammation-induced facilitation of brush-evoked activity (number of spikes: $F_{(1,38)} = 119.6$, $p < 0.0001$; low intensity: $q = 2.34$, $p = 0.0478$; high intensity: $q = 13.12$, $p < 0.0001$; and peak IFF: $F_{(1,38)} = 96.42$, $p = 0.0007$; low: $q = 4.31$, $p = 0.0002$; high: $q = 9.56$, $p < 0.0001$ $n = 20$; RM two-way ANOVA). (**B**) C-HTMRs displayed reduced responsiveness to high-intensity brushing following inflammation (number of spikes: $F_{(1,24)} = 12.84$, $p = 0.0015$; high intensity: $q = 5.06$, $p < 0.0001$; and peak IFF: $F_{(1,24)} = 2.90$, $p = 0.1014$; high intensity: $q = 2.40$, $p = 0.0475$; $n = 13$). (**C**) Field-LTMRs demonstrated reduced responses across both brush intensities (number of spikes: $F_{(1,24)} = 0.5225$, $p = 0.4767$; low: $q = 4.86$, $p = 0.0001$; high: $q = 5.89$, $p < 0.0001$; and peak IFF: $F_{(1,24)} = 0.38$, $p = 0.5414$; low: $q = 6.49$, $p < 0.0001$; high: $q = 5.61$, $p < 0.0001$; $n = 13$). (**D**) SA-LTMRs also exhibited pronounced attenuation of both intensities of brush-evoked responses following inflammation (number of spikes: $F_{(1,24)} = 14.79$, $p = 0.0008$; low: $q = 2.55$, $p = 0.0345$; high: $q = 7.99$, $p < 0.0001$; and peak IFF: $F_{(1,24)} = 14.22$, $p = 0.0009$; low: $q = 2.38$, $p = 0.05$; high: $q = 7.71$, $p < 0.0001$; $n = 13$). (**E**) C-LTMR$^{CDH9}$ showed no significant inflammation-dependent changes (number of spikes: $F_{(1,10)} = 0.40$, $p = 0.538$; low: $q = 0.90$, $p = 0.6260$; high: $q = 1.80$, $p = 0.1927$; and peak IFF: $F_{(1,10)} = 0.23$, $p = 0.6356$; low: $q = 1.80$, $p = 0.1922$; high: $q = 1.11$, $p = 0.4978$; $n = 6$). Asterisks (where shown) on the force-response data (A-D) indicate significant post hoc comparisons (Sidak's test). (**F**) Aβ-LTMR(HF)$^{CCKAR}$ identified by preferential sensitivity to hair movement, were only tested with low-intensity brushing. They maintained stable responses to low-intensity brushing following inflammation (number of spikes: $t = 1.62$, $p = 0.1271$; peak IFF: $t = 0.34$, $p = 0.7348$; $n = 3$; paired t-test). *$p < 0.05$, **$p < 0.01$, ***$p < 0.001$, ****$p < 0.0001$.

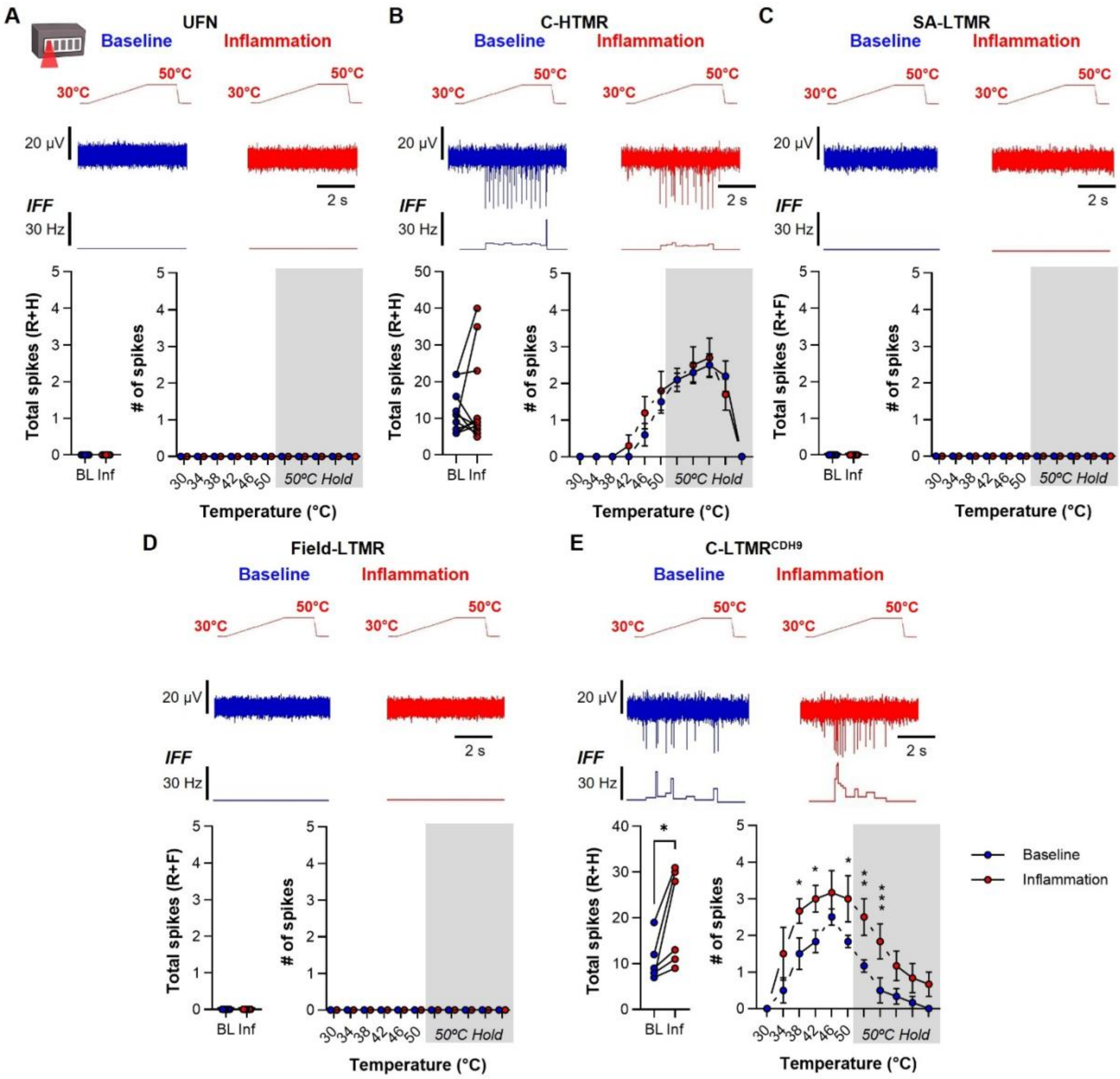


**Fig. S7. Heating test stimuli evoked no responses in UFNs or Aβ-LTMRs at baseline or after inflammation**

Representative recordings and quantitative analyses of neuronal responses to heat stimulation (30 °C → 50 °C; 5 s hold) under baseline conditions (blue) and following heat-induced inflammation (red). Data displayed as in Fig. 2A-F. (**A,C,D**) UFNs (n = 20), SA-LTMRs (n = 13), and Field-LTMRs (n = 13) do not exhibit heat-evoked activity either at baseline or following heat-induced inflammation. (**B**) C-HTMRs (n = 10) displayed similar heat-evoked responses under inflammation condition. Total spike counts (t = 0.89, p = 0.3935; paired t-test), and temperature–response curves (inflammation: $F_{(1,9)} = 0.29$, p = 0.5991; RM two-way ANOVA with Sidak's test) remained unchanged following inflammation. (**E**) C-LTMR$^{CDH9}$ (n = 6) demonstrated a modest increase in overall heat-evoked activity following inflammation (total spike counts: t = 2.97, p =

0.0308). Further, temperature–response curves revealed an increase in firing dynamics during the heat ramp (inflammation: $F_{(1,5)} = 8.87$, $p = 0.0308$).

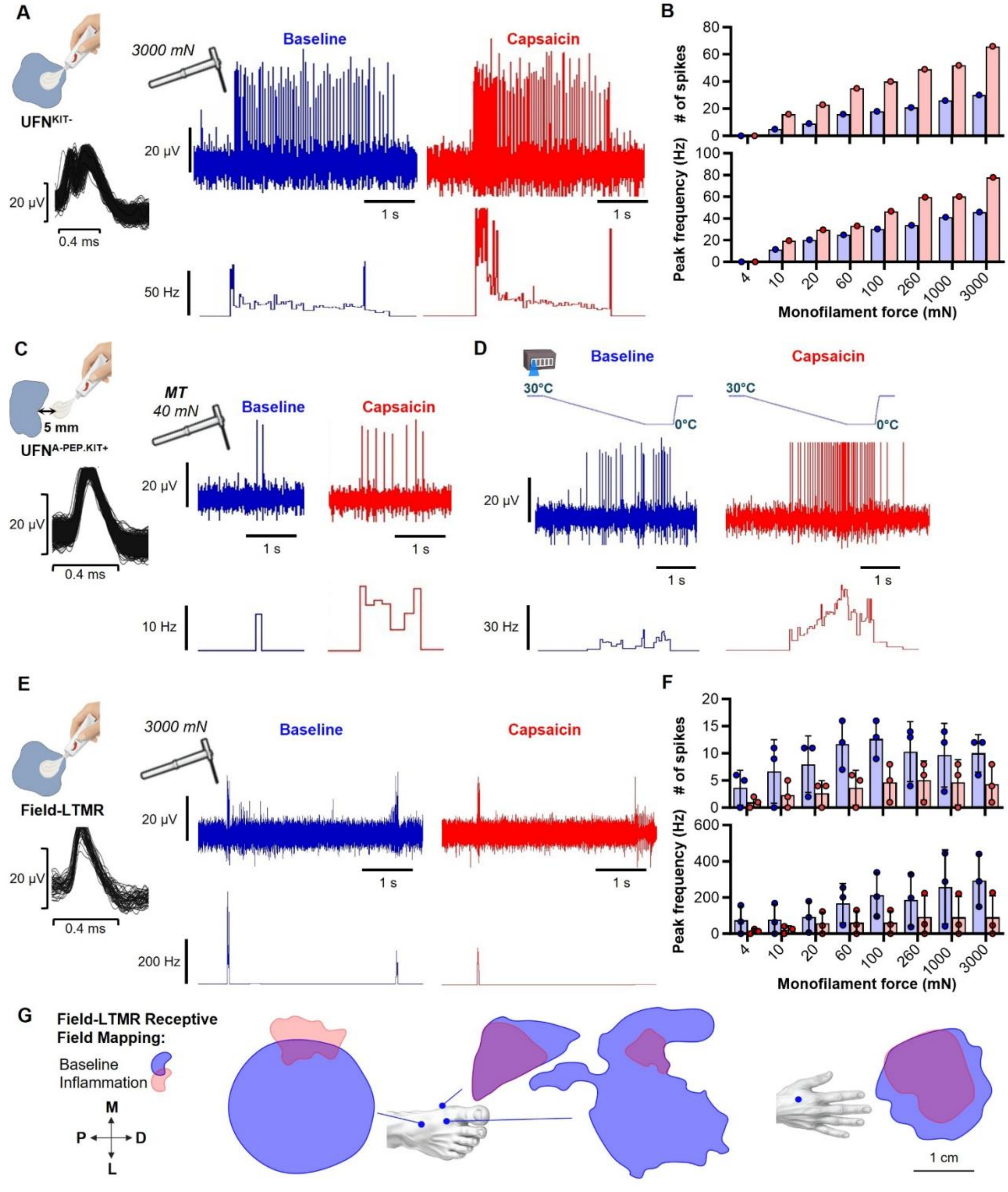


**Fig. S8. Capsaicin-induced inflammation sensitizes UFNs but desensitizes Field-LTMRs.** Topical capsaicin application produced effects similar to heat-evoked inflammation. Two UFNs (UFN$^{KIT-}$, **A**–**B**; UFN$^{A-PEP.KIT}$, **C**–**D**) and four Field-LTMRs were recorded during capsaicin-induced inflammation. Both UFNs exhibited robust sensitization following capsaicin exposure, as reflected by marked reductions in mechanical threshold (from 150 → 20 mN and 40 → 1.6 mN). In contrast, Field-LTMRs showed desensitization, with mechanical thresholds increasing (from 0.07–1.6 mN at baseline to 10–20 mN after capsaicin). (**A**) Representative single-unit UFN recording showing action-potential waveform (left) and responses to punctate indentation (3000 mN) at baseline (blue) and 5 min after capsaicin application inside the receptive field (RF; red). Corresponding instantaneous (IFF) plots demonstrate increased spiking following capsaicin-induced inflammation. (**B**) Quantification of total spike count (top) and peak firing frequency (bottom) evoked by graded forces (4–3000 mN). Capsaicin enhanced firing across a broad force range. (**C**) Example UFN response at the original mechanical threshold (40 mN) before (blue) and 5 min after (red) capsaicin applied 5 mm outside the RF. Increased spiking at the baseline threshold is indicative of mechanical sensitization; IFF shown below. (**D**) Cooling-evoked responses at baseline (blue) and after capsaicin (red), demonstrating enhanced cold sensitivity. (**E**) Representative single-unit Field recording showing action-potential waveform (left) and responses to punctate indentation (3000 mN) at baseline and after capsaicin application inside the RF. Corresponding IFF plots demonstrate decreased spiking following inflammation. (**F**) Quantification of total spike count and peak firing frequency evoked by graded forces (4–3000 mN; 3 units). Capsaicin reduced firing across a broad force range. (**G**) Location and spatial organization (relative scaling) of all recorded Field-LTMR RFs (n=4) before and after capsaicin application. Boundaries of the mechanical RFs are shown for baseline (blue) and post-inflammation (red) with overlap (purple). Capsaicin reduced RF size in Field-LTMRs.

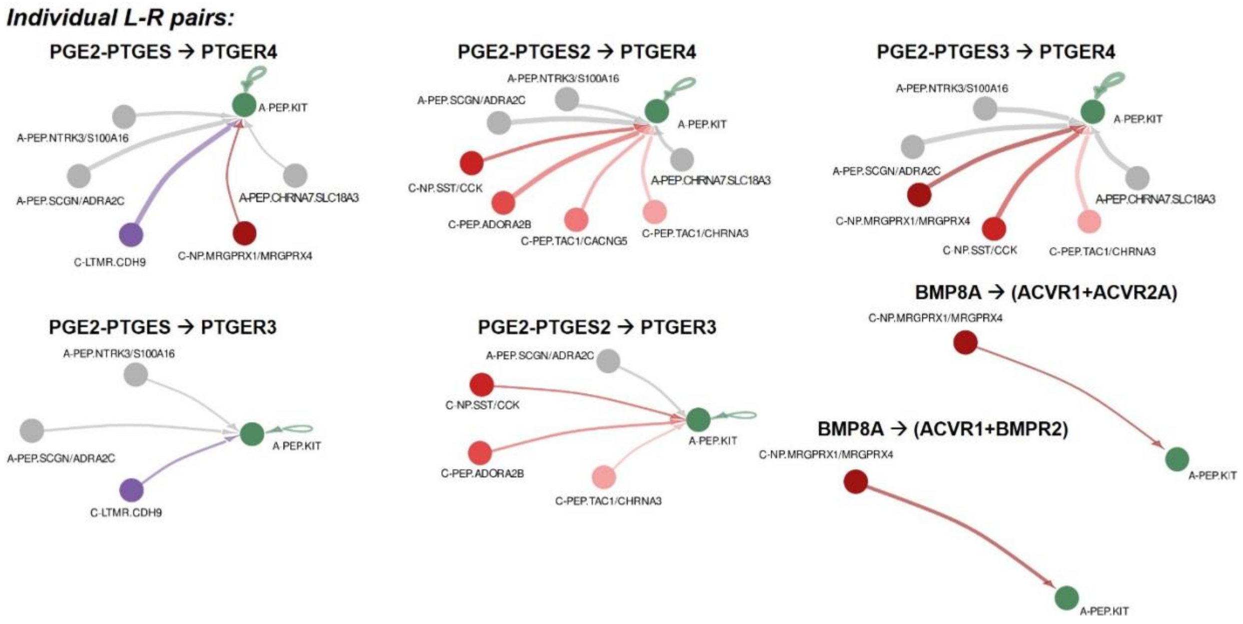


**Fig. S9. Ligand-receptor interaction from TRPV1+ DRG neurons to A-PEP.KIT neurons.** Circle plots showing possible Ligand-receptor (L-R) pairs in the classes "Secreted Signaling" and "Nonprotein Signaling" between TRPV1+ DRG neuron types and the A-PEP.KIT type. The direction of signaling goes towards A-PEP.KIT for each interaction (arrowheads). Thickness of the line represents the strength of possible interaction.

### Table S1. Ligand-receptor interactions.

| source | ligand | receptor | prob | pval | interaction_name | interaction_name_2 | pathway_name | annotation | evidence |
|---|---|---|---|---|---|---|---|---|---|
| C-NP.MRGPRX1/MRGPRX4 | TGFB1 | TGFbR1_R2 | 0.000212375 | 0 | TGFB1_TGFBR1_TGFBR2 | TGFB1 - (TGFBR1+TGFBR2) | TGFb | Secreted Signaling | KEGG: hsa04350 |
| A-PEP.KIT | TGFB1 | ACVR1B_TGFbR2 | 8.47686E-05 | 0.02 | TGFB1_ACVR1B_TGFBR2 | TGFB1 - (ACVR1B+TGFBR2) | TGFb | Secreted Signaling | PMID: 27449815 |
| C-NP.MRGPRX1/MRGPRX4 | TGFB1 | ACVR1B_TGFbR2 | 0.000139318 | 0 | TGFB1_ACVR1B_TGFBR2 | TGFB1 - (ACVR1B+TGFBR2) | TGFb | Secreted Signaling | PMID: 27449815 |
| C-NP.MRGPRX1/MRGPRX4 | TGFB1 | ACVR1_TGFbR | 9.15494E-05 | 0 | TGFB1_ACVR1_TGFBR1 | TGFB1 - (ACVR1+TGFBR1) | TGFb | Secreted Signaling | PMID: 29376829 |
| C-PEP.TAC1/CACNG5 | BMP6 | ACVR1_ACVR2A | 1.35651E-05 | 0 | BMP6_ACVR1_ACVR2A | BMP6 - (ACVR1+ACVR2A) | BMP | Secreted Signaling | KEGG: hsa04350; PMID:26893264 |
| C-PEP.TAC1/CHRNA3 | BMP6 | ACVR1_ACVR2A | 3.92051E-05 | 0 | BMP6_ACVR1_ACVR2A | BMP6 - (ACVR1+ACVR2A) | BMP | Secreted Signaling | KEGG: hsa04350; PMID:26893264 |
| C-PEP.TAC1/CACNG5 | BMP6 | ACVR1_ACVR2B | 4.76609E-06 | 0 | BMP6_ACVR1_ACVR2B | BMP6 - (ACVR1+ACVR2B) | BMP | Secreted Signaling | KEGG: hsa04350; PMID:26893264 |
| C-PEP.TAC1/CHRNA3 | BMP6 | ACVR1_ACVR2B | 1.37751E-05 | 0 | BMP6_ACVR1_ACVR2B | BMP6 - (ACVR1+ACVR2B) | BMP | Secreted Signaling | KEGG: hsa04350; PMID:26893264 |
| C-PEP.TAC1/CACNG5 | BMP6 | ACVR1_BMPR2 | 4.46862E-05 | 0 | BMP6_ACVR1_BMPR2 | BMP6 - (ACVR1+BMPR2) | BMP | Secreted Signaling | KEGG: hsa04350; PMID:26893264 |
| C-PEP.TAC1/CHRNA3 | BMP6 | ACVR1_BMPR2 | 0.000129137 | 0 | BMP6_ACVR1_BMPR2 | BMP6 - (ACVR1+BMPR2) | BMP | Secreted Signaling | KEGG: hsa04350; PMID:26893264 |
| C-PEP.TAC1/CACNG5 | BMP6 | BMPR1A_ACVR2A | 2.13685E-05 | 0 | BMP6_BMPR1A_ACVR2A | BMP6 - (BMPR1A+ACVR2A) | BMP | Secreted Signaling | KEGG: hsa04350; PMID:26893264 |
| C-PEP.TAC1/CHRNA3 | BMP6 | BMPR1A_ACVR2A | 6.17567E-05 | 0 | BMP6_BMPR1A_ACVR2A | BMP6 - (BMPR1A+ACVR2A) | BMP | Secreted Signaling | KEGG: hsa04350; PMID:26893264 |
| C-PEP.TAC1/CACNG5 | BMP6 | BMPR1A_ACVR2B | 7.50788E-06 | 0 | BMP6_BMPR1A_ACVR2B | BMP6 - (BMPR1A+ACVR2B) | BMP | Secreted Signaling | KEGG: hsa04350; PMID:26893264 |
| C-PEP.TAC1/CHRNA3 | BMP6 | BMPR1A_ACVR2B | 2.16993E-05 | 0 | BMP6_BMPR1A_ACVR2B | BMP6 - (BMPR1A+ACVR2B) | BMP | Secreted Signaling | KEGG: hsa04350; PMID:26893264 |
| C-PEP.TAC1/CACNG5 | BMP6 | BMPR1A_BMPR2 | 7.03904E-05 | 0 | BMP6_BMPR1A_BMPR2 | BMP6 - (BMPR1A+BMPR2) | BMP | Secreted Signaling | KEGG: hsa04350; PMID:26893264 |
| C-PEP.TAC1/CHRNA3 | BMP6 | BMPR1A_BMPR2 | 0.000203403 | 0 | BMP6_BMPR1A_BMPR2 | BMP6 - (BMPR1A+BMPR2) | BMP | Secreted Signaling | KEGG: hsa04350; PMID:26893264 |

| | | | | | | | | | |
|---|---|---|---|---|---|---|---|---|---|
| C-NP.MRGPRX1/MRGPRX4 | BMP7 | ACVR1_ACVR2A | 1.65057E-05 | 0.04 | BMP7_ACVR1_ACVR2A | BMP7 - (ACVR1+ACVR2A) | BMP | Secreted Signaling | KEGG: hsa04350; PMID:26893264 |
| C-NP.MRGPRX1/MRGPRX4 | BMP7 | ACVR1_ACVR2B | 5.79931E-06 | 0.04 | BMP7_ACVR1_ACVR2B | BMP7 - (ACVR1+ACVR2B) | BMP | Secreted Signaling | KEGG: hsa04350; PMID:26893264 |
| C-NP.MRGPRX1/MRGPRX4 | BMP7 | ACVR1_BMPR2 | 5.43724E-05 | 0.04 | BMP7_ACVR1_BMPR2 | BMP7 - (ACVR1+BMPR2) | BMP | Secreted Signaling | KEGG: hsa04350; PMID:26893264 |
| C-NP.MRGPRX1/MRGPRX4 | BMP7 | BMPR1A_ACVR2A | 2.60007E-05 | 0 | BMP7_BMPR1A_ACVR2A | BMP7 - (BMPR1A+ACVR2A) | BMP | Secreted Signaling | KEGG: hsa04350; PMID:26893264 |
| C-NP.MRGPRX1/MRGPRX4 | BMP7 | BMPR1A_ACVR2B | 9.13547E-06 | 0.03 | BMP7_BMPR1A_ACVR2B | BMP7 - (BMPR1A+ACVR2B) | BMP | Secreted Signaling | KEGG: hsa04350; PMID:26893264 |
| C-NP.MRGPRX1/MRGPRX4 | BMP7 | BMPR1A_BMPR2 | 8.56471E-05 | 0 | BMP7_BMPR1A_BMPR2 | BMP7 - (BMPR1A+BMPR2) | BMP | Secreted Signaling | KEGG: hsa04350; PMID:26893264 |
| C-NP.MRGPRX1/MRGPRX4 | BMP8A | ACVR1_ACVR2A | 0.000371206 | 0.01 | BMP8A_ACVR1_ACVR2A | BMP8A - (ACVR1+ACVR2A) | BMP | Secreted Signaling | KEGG: hsa04350; PMID:26893264 |
| C-NP.MRGPRX1/MRGPRX4 | BMP8A | ACVR1_BMPR2 | 0.001221136 | 0 | BMP8A_ACVR1_BMPR2 | BMP8A - (ACVR1+BMPR2) | BMP | Secreted Signaling | KEGG: hsa04350; PMID:26893264 |
| C-NP.MRGPRX1/MRGPRX4 | BMP8A | BMPR1A_ACVR2A | 0.000584542 | 0 | BMP8A_BMPR1A_ACVR2A | BMP8A - (BMPR1A+ACVR2A) | BMP | Secreted Signaling | KEGG: hsa04350; PMID:26893264 |
| C-NP.MRGPRX1/MRGPRX4 | BMP8A | BMPR1A_ACVR2B | 0.000205507 | 0 | BMP8A_BMPR1A_ACVR2B | BMP8A - (BMPR1A+ACVR2B) | BMP | Secreted Signaling | KEGG: hsa04350; PMID:26893264 |
| C-NP.MRGPRX1/MRGPRX4 | BMP8A | BMPR1A_BMPR2 | 0.001921354 | 0 | BMP8A_BMPR1A_BMPR2 | BMP8A - (BMPR1A+BMPR2) | BMP | Secreted Signaling | KEGG: hsa04350; PMID:26893264 |
| C-NP.SST/CCK | BMP8A | BMPR1A_BMPR2 | 0.000796678 | 0.02 | BMP8A_BMPR1A_BMPR2 | BMP8A - (BMPR1A+BMPR2) | BMP | Secreted Signaling | KEGG: hsa04350; PMID:26893264 |
| C-LTMR.CDH9 | GDF10 | ACVR1B_ACVR2A | 5.09415E-05 | 0.02 | GDF10_ACVR1B_ACVR2A | GDF10 - (ACVR1B+ACVR2A) | GDF | Secreted Signaling | KEGG: hsa04350 |
| C-NP.SST/CCK | GDF10 | ACVR1B_ACVR2A | 4.23425E-05 | 0 | GDF10_ACVR1B_ACVR2A | GDF10 - (ACVR1B+ACVR2A) | GDF | Secreted Signaling | KEGG: hsa04350 |
| A-PEP.SCGN/ADRA2C | GDF11 | TGFBR1_ACVR2A | 0.005169745 | 0 | GDF11_TGFBR1_ACVR2A | GDF11 - (TGFBR1+ACVR2A) | GDF | Secreted Signaling | KEGG: hsa04350 |
| C-PEP.ADORA2B | GDF11 | TGFBR1_ACVR2A | 0.003932195 | 0.02 | GDF11_TGFBR1_ACVR2A | GDF11 - (TGFBR1+ACVR2A) | GDF | Secreted Signaling | KEGG: hsa04350 |

| | | | | | | | | | |
|---|---|---|---|---|---|---|---|---|---|
| A-PEP.SCGN/ADRA2C | GDF11 | ACVR1B_ACVR2B | 0.001196194 | 0.02 | GDF11_ACVR1B_ACVR2B | GDF11 - (ACVR1B+ACVR2B) | GDF | Secreted Signaling | KEGG: hsa04350 |
| C-PEP.ADORA2B | GDF11 | ACVR1B_ACVR2B | 0.000908976 | 0.04 | GDF11_ACVR1B_ACVR2B | GDF11 - (ACVR1B+ACVR2B) | GDF | Secreted Signaling | KEGG: hsa04350 |
| A-PEP.CHRNA7/SLC18A3 | WNT10B | FZD3_LRP6 | 6.93979E-05 | 0 | WNT10B_FZD3_LRP6 | WNT10B - (FZD3+LRP6) | WNT | Secreted Signaling | KEGG: hsa04310; PMID: 23209147 |
| A-PEP.SCGN/ADRA2C | WNT10B | FZD3_LRP6 | 0.000412868 | 0 | WNT10B_FZD3_LRP6 | WNT10B - (FZD3+LRP6) | WNT | Secreted Signaling | KEGG: hsa04310; PMID: 23209147 |
| C-PEP.TAC1/CACNG5 | WNT10B | FZD3_LRP6 | 0.000217736 | 0 | WNT10B_FZD3_LRP6 | WNT10B - (FZD3+LRP6) | WNT | Secreted Signaling | KEGG: hsa04310; PMID: 23209147 |
| A-PEP.CHRNA7/SLC18A3 | WNT10B | FZD4_LRP6 | 3.21388E-05 | 0 | WNT10B_FZD4_LRP6 | WNT10B - (FZD4+LRP6) | WNT | Secreted Signaling | KEGG: hsa04310; PMID: 23209147 |
| A-PEP.SCGN/ADRA2C | WNT10B | FZD4_LRP6 | 0.000191235 | 0 | WNT10B_FZD4_LRP6 | WNT10B - (FZD4+LRP6) | WNT | Secreted Signaling | KEGG: hsa04310; PMID: 23209147 |
| C-PEP.TAC1/CACNG5 | WNT10B | FZD4_LRP6 | 0.000100846 | 0 | WNT10B_FZD4_LRP6 | WNT10B - (FZD4+LRP6) | WNT | Secreted Signaling | KEGG: hsa04310; PMID: 23209147 |
| A-PEP.CHRNA7/SLC18A3 | WNT10B | FZD7_LRP6 | 5.04389E-06 | 0 | WNT10B_FZD7_LRP6 | WNT10B - (FZD7+LRP6) | WNT | Secreted Signaling | KEGG: hsa04310; PMID: 23209147 |
| A-PEP.SCGN/ADRA2C | WNT10B | FZD7_LRP6 | 3.00162E-05 | 0.01 | WNT10B_FZD7_LRP6 | WNT10B - (FZD7+LRP6) | WNT | Secreted Signaling | KEGG: hsa04310; PMID: 23209147 |
| A-PEP.CHRNA7/SLC18A3 | WNT10B | FZD8_LRP6 | 1.21148E-05 | 0 | WNT10B_FZD8_LRP6 | WNT10B - (FZD8+LRP6) | WNT | Secreted Signaling | KEGG: hsa04310; PMID: 23209147 |
| A-PEP.SCGN/ADRA2C | WNT10B | FZD8_LRP6 | 7.2093E-05 | 0 | WNT10B_FZD8_LRP6 | WNT10B - (FZD8+LRP6) | WNT | Secreted Signaling | KEGG: hsa04310; PMID: 23209147 |
| C-PEP.TAC1/CACNG5 | WNT10B | FZD8_LRP6 | 3.80163E-05 | 0.02 | WNT10B_FZD8_LRP6 | WNT10B - (FZD8+LRP6) | WNT | Secreted Signaling | KEGG: hsa04310; PMID: 23209147 |
| A-PEP.NTRK3/S100A16 | WNT2B | FZD3_LRP6 | 0.000127695 | 0 | WNT2B_FZD3_LRP6 | WNT2B - (FZD3+LRP6) | WNT | Secreted Signaling | KEGG: hsa04310; PMID: 23209147 |
| A-PEP.NTRK3/S100A16 | WNT2B | FZD4_LRP6 | 5.9138E-05 | 0 | WNT2B_FZD4_LRP6 | WNT2B - (FZD4+LRP6) | WNT | Secreted Signaling | KEGG: hsa04310; PMID: 23209147 |
| A-PEP.SCGN/ADRA2C | WNT2B | FZD4_LRP6 | 3.20741E-05 | 0.03 | WNT2B_FZD4_LRP6 | WNT2B - (FZD4+LRP6) | WNT | Secreted Signaling | KEGG: hsa04310; PMID: 23209147 |
| A-PEP.NTRK3/S100A16 | WNT2B | FZD7_LRP6 | 9.28131E-06 | 0.02 | WNT2B_FZD7_LRP6 | WNT2B - (FZD7+LRP6) | WNT | Secreted Signaling | KEGG: hsa04310; PMID: 23209147 |
| A-PEP.NTRK3/S100A16 | WNT2B | FZD8_LRP6 | 2.22925E-05 | 0 | WNT2B_FZD8_LRP6 | WNT2B - (FZD8+LRP6) | WNT | Secreted Signaling | KEGG: hsa04310; PMID: 23209147 |
| A-PEP.KIT | WNT3 | FZD3_LRP6 | 0.000622646 | 0 | WNT3_FZD3_LRP6 | WNT3 - (FZD3+LRP6) | WNT | Secreted Signaling | KEGG: hsa04310; PMID: 23209147 |
| A-PEP.SCGN/ADRA2C | WNT3 | FZD3_LRP6 | 0.000181187 | 0.01 | WNT3_FZD3_LRP6 | WNT3 - (FZD3+LRP6) | WNT | Secreted Signaling | KEGG: hsa04310; PMID: 23209147 |
| A-PEP.KIT | WNT3 | FZD4_LRP6 | 0.000288432 | 0 | WNT3_FZD4_LRP6 | WNT3 - (FZD4+LRP6) | WNT | Secreted Signaling | KEGG: hsa04310; PMID: 23209147 |

| | | | | | | | | | |
|---|---|---|---|---|---|---|---|---|---|
| A-PEP.SCGN/ADRA2C | WNT3 | FZD4_LRP6 | 8.39134 E-05 | 0.01 | WNT3_FZD4_LRP6 | WNT3 - (FZD4+LRP6) | WNT | Secreted Signaling | KEGG: hsa04310; PMID: 23209147 |
| A-PEP.KIT | WNT3 | FZD7_LRP6 | 4.52758 E-05 | 0.02 | WNT3_FZD7_LRP6 | WNT3 - (FZD7+LRP6) | WNT | Secreted Signaling | KEGG: hsa04310; PMID: 23209147 |
| A-PEP.SCGN/ADRA2C | WNT3 | FZD7_LRP6 | 1.31699 E-05 | 0.02 | WNT3_FZD7_LRP6 | WNT3 - (FZD7+LRP6) | WNT | Secreted Signaling | KEGG: hsa04310; PMID: 23209147 |
| A-PEP.KIT | WNT3 | FZD8_LRP6 | 0.00010 8741 | 0 | WNT3_FZD8_LRP6 | WNT3 - (FZD8+LRP6) | WNT | Secreted Signaling | KEGG: hsa04310; PMID: 23209147 |
| A-PEP.SCGN/ADRA2C | WNT3 | FZD8_LRP6 | 3.16322 E-05 | 0.02 | WNT3_FZD8_LRP6 | WNT3 - (FZD8+LRP6) | WNT | Secreted Signaling | KEGG: hsa04310; PMID: 23209147 |
| A-PEP.CHRNA7/SLC18 A3 | WNT7B | FZD3_LRP6 | 0.00018 5504 | 0 | WNT7B_FZD3_LRP 6 | WNT7B - (FZD3+LRP6) | WNT | Secreted Signaling | KEGG: hsa04310; PMID: 23209147 |
| A-PEP.KIT | WNT7B | FZD3_LRP6 | 0.00207 4922 | 0 | WNT7B_FZD3_LRP 6 | WNT7B - (FZD3+LRP6) | WNT | Secreted Signaling | KEGG: hsa04310; PMID: 23209147 |
| A-PEP.NTRK3/S100A16 | WNT7B | FZD3_LRP6 | 0.00013 0834 | 0.02 | WNT7B_FZD3_LRP 6 | WNT7B - (FZD3+LRP6) | WNT | Secreted Signaling | KEGG: hsa04310; PMID: 23209147 |
| A-PEP.SCGN/ADRA2C | WNT7B | FZD3_LRP6 | 0.00431 5934 | 0 | WNT7B_FZD3_LRP 6 | WNT7B - (FZD3+LRP6) | WNT | Secreted Signaling | KEGG: hsa04310; PMID: 23209147 |
| C-PEP.ADORA2B | WNT7B | FZD3_LRP6 | 0.00117 8011 | 0 | WNT7B_FZD3_LRP 6 | WNT7B - (FZD3+LRP6) | WNT | Secreted Signaling | KEGG: hsa04310; PMID: 23209147 |
| A-PEP.CHRNA7/SLC18 A3 | WNT7B | FZD4_LRP6 | 8.59148 E-05 | 0 | WNT7B_FZD4_LRP 6 | WNT7B - (FZD4+LRP6) | WNT | Secreted Signaling | KEGG: hsa04310; PMID: 23209147 |
| A-PEP.KIT | WNT7B | FZD4_LRP6 | 0.00096 1891 | 0 | WNT7B_FZD4_LRP 6 | WNT7B - (FZD4+LRP6) | WNT | Secreted Signaling | KEGG: hsa04310; PMID: 23209147 |
| A-PEP.NTRK3/S100A16 | WNT7B | FZD4_LRP6 | 6.05921 E-05 | 0.02 | WNT7B_FZD4_LRP 6 | WNT7B - (FZD4+LRP6) | WNT | Secreted Signaling | KEGG: hsa04310; PMID: 23209147 |
| A-PEP.SCGN/ADRA2C | WNT7B | FZD4_LRP6 | 0.00200 3043 | 0 | WNT7B_FZD4_LRP 6 | WNT7B - (FZD4+LRP6) | WNT | Secreted Signaling | KEGG: hsa04310; PMID: 23209147 |
| C-PEP.ADORA2B | WNT7B | FZD4_LRP6 | 0.00054 5867 | 0 | WNT7B_FZD4_LRP 6 | WNT7B - (FZD4+LRP6) | WNT | Secreted Signaling | KEGG: hsa04310; PMID: 23209147 |
| A-PEP.CHRNA7/SLC18 A3 | WNT7B | FZD7_LRP6 | 1.34842 E-05 | 0.04 | WNT7B_FZD7_LRP 6 | WNT7B - (FZD7+LRP6) | WNT | Secreted Signaling | KEGG: hsa04310; PMID: 23209147 |
| A-PEP.KIT | WNT7B | FZD7_LRP6 | 0.00015 1072 | 0 | WNT7B_FZD7_LRP 6 | WNT7B - (FZD7+LRP6) | WNT | Secreted Signaling | KEGG: hsa04310; PMID: 23209147 |
| A-PEP.SCGN/ADRA2C | WNT7B | FZD7_LRP6 | 0.00031 4851 | 0 | WNT7B_FZD7_LRP 6 | WNT7B - (FZD7+LRP6) | WNT | Secreted Signaling | KEGG: hsa04310; PMID: 23209147 |
| C-PEP.ADORA2B | WNT7B | FZD7_LRP6 | 8.57053 E-05 | 0 | WNT7B_FZD7_LRP 6 | WNT7B - (FZD7+LRP6) | WNT | Secreted Signaling | KEGG: hsa04310; PMID: 23209147 |
| A-PEP.CHRNA7/SLC18 A3 | WNT7B | FZD8_LRP6 | 3.2387 E-05 | 0 | WNT7B_FZD8_LRP 6 | WNT7B - (FZD8+LRP6) | WNT | Secreted Signaling | KEGG: hsa04310; PMID: 23209147 |
| A-PEP.KIT | WNT7B | FZD8_LRP6 | 0.00036 2785 | 0 | WNT7B_FZD8_LRP 6 | WNT7B - (FZD8+LRP6) | WNT | Secreted Signaling | KEGG: hsa04310; PMID: 23209147 |
| A-PEP.NTRK3/S100A16 | WNT7B | FZD8_LRP6 | 2.28406 E-05 | 0.03 | WNT7B_FZD8_LRP 6 | WNT7B - (FZD8+LRP6) | WNT | Secreted Signaling | KEGG: hsa04310; PMID: 23209147 |

| | | | | | | | | | |
|---|---|---|---|---|---|---|---|---|---|
| A-PEP.SCGN/ADRA2C | WNT7B | FZD8_LRP6 | 0.000755925 | 0 | WNT7B_FZD8_LRP6 | WNT7B - (FZD8+LRP6) | WNT | Secreted Signaling | KEGG: hsa04310; PMID: 23209147 |
| C-PEP.ADORA2B | WNT7B | FZD8_LRP6 | 0.000205831 | 0 | WNT7B_FZD8_LRP6 | WNT7B - (FZD8+LRP6) | WNT | Secreted Signaling | KEGG: hsa04310; PMID: 23209147 |
| A-PEP.SCGN/ADRA2C | WNT9A | FZD3_LRP6 | 0.001405682 | 0 | WNT9A_FZD3_LRP6 | WNT9A - (FZD3+LRP6) | WNT | Secreted Signaling | KEGG: hsa04310; PMID: 23209147 |
| C-NP.MRGPRX1/MRGPRX4 | WNT9A | FZD3_LRP6 | 0.000444941 | 0 | WNT9A_FZD3_LRP6 | WNT9A - (FZD3+LRP6) | WNT | Secreted Signaling | KEGG: hsa04310; PMID: 23209147 |
| C-PEP.ADORA2B | WNT9A | FZD3_LRP6 | 0.000791908 | 0.01 | WNT9A_FZD3_LRP6 | WNT9A - (FZD3+LRP6) | WNT | Secreted Signaling | KEGG: hsa04310; PMID: 23209147 |
| A-PEP.SCGN/ADRA2C | WNT9A | FZD4_LRP6 | 0.00065142 | 0 | WNT9A_FZD4_LRP6 | WNT9A - (FZD4+LRP6) | WNT | Secreted Signaling | KEGG: hsa04310; PMID: 23209147 |
| C-NP.MRGPRX1/MRGPRX4 | WNT9A | FZD4_LRP6 | 0.000206104 | 0 | WNT9A_FZD4_LRP6 | WNT9A - (FZD4+LRP6) | WNT | Secreted Signaling | KEGG: hsa04310; PMID: 23209147 |
| C-PEP.ADORA2B | WNT9A | FZD4_LRP6 | 0.000366879 | 0.01 | WNT9A_FZD4_LRP6 | WNT9A - (FZD4+LRP6) | WNT | Secreted Signaling | KEGG: hsa04310; PMID: 23209147 |
| A-PEP.SCGN/ADRA2C | WNT9A | FZD8_LRP6 | 0.000245643 | 0 | WNT9A_FZD8_LRP6 | WNT9A - (FZD8+LRP6) | WNT | Secreted Signaling | KEGG: hsa04310; PMID: 23209147 |
| C-PEP.ADORA2B | WNT9A | FZD8_LRP6 | 0.000138324 | 0.02 | WNT9A_FZD8_LRP6 | WNT9A - (FZD8+LRP6) | WNT | Secreted Signaling | KEGG: hsa04310; PMID: 23209147 |
| C-NP.MRGPRX1/MRGPRX4 | WNT5A | FZD3 | 0.000877222 | 0 | WNT5A_FZD3 | WNT5A - FZD3 | ncWNT | Secreted Signaling | KEGG: hsa04310 |
| A-PEP.SCGN/ADRA2C | WNT5A | FZD4 | 0.000171035 | 0.02 | WNT5A_FZD4 | WNT5A - FZD4 | ncWNT | Secreted Signaling | KEGG: hsa04310 |
| C-LTMR.CDH9 | WNT5A | FZD4 | 0.000337709 | 0.02 | WNT5A_FZD4 | WNT5A - FZD4 | ncWNT | Secreted Signaling | KEGG: hsa04310 |
| C-NP.MRGPRX1/MRGPRX4 | WNT5A | FZD4 | 0.000188252 | 0.01 | WNT5A_FZD4 | WNT5A - FZD4 | ncWNT | Secreted Signaling | KEGG: hsa04310 |
| A-PEP.SCGN/ADRA2C | WNT5A | MCAM | 0.000499033 | 0 | WNT5A_MCAM | WNT5A - MCAM | ncWNT | Secreted Signaling | PMID: 24335906 |
| C-LTMR.CDH9 | WNT5A | MCAM | 0.000985029 | 0 | WNT5A_MCAM | WNT5A - MCAM | ncWNT | Secreted Signaling | PMID: 24335906 |
| C-NP.MRGPRX1/MRGPRX4 | WNT5A | MCAM | 0.00054925 | 0 | WNT5A_MCAM | WNT5A - MCAM | ncWNT | Secreted Signaling | PMID: 24335906 |
| A-PEP.CHRNA7/SLC18A3 | WNT11 | FZD3 | 0.000213835 | 0 | WNT11_FZD3 | WNT11 - FZD3 | ncWNT | Secreted Signaling | KEGG: hsa04310 |
| A-PEP.SCGN/ADRA2C | WNT11 | FZD3 | 0.001939581 | 0 | WNT11_FZD3 | WNT11 - FZD3 | ncWNT | Secreted Signaling | KEGG: hsa04310 |
| A-PEP.CHRNA7/SLC18A3 | WNT11 | FZD4 | 4.5865E-05 | 0 | WNT11_FZD4 | WNT11 - FZD4 | ncWNT | Secreted Signaling | KEGG: hsa04310 |
| A-PEP.KIT | WNT11 | FZD4 | 5.24885E-05 | 0.03 | WNT11_FZD4 | WNT11 - FZD4 | ncWNT | Secreted Signaling | KEGG: hsa04310 |

| | | | | | | | | | |
|---|---|---|---|---|---|---|---|---|---|
| A-PEP.NTRK3/S100A16 | WNT11 | FZD4 | 6.07766E-06 | 0.04 | WNT11_FZD4 | WNT11 - FZD4 | ncWNT | Secreted Signaling | KEGG: hsa04310 |
| A-PEP.SCGN/ADRA2C | WNT11 | FZD4 | 0.000416582 | 0 | WNT11_FZD4 | WNT11 - FZD4 | ncWNT | Secreted Signaling | KEGG: hsa04310 |
| A-PEP.CHRNA7/SLC18A3 | WNT11 | FZD7 | 1.12966E-06 | 0.03 | WNT11_FZD7 | WNT11 - FZD7 | ncWNT | Secreted Signaling | KEGG: hsa04310 |
| A-PEP.SCGN/ADRA2C | WNT11 | FZD7 | 1.02641E-05 | 0.01 | WNT11_FZD7 | WNT11 - FZD7 | ncWNT | Secreted Signaling | KEGG: hsa04310 |
| C-NP.SST/CCK | FGF1 | FGFR1 | 0.047892117 | 0 | FGF1_FGFR1 | FGF1 - FGFR1 | FGF | Secreted Signaling | PMC: 4393358 |
| A-PEP.KIT | FGF5 | FGFR1 | 0.009155068 | 0 | FGF5_FGFR1 | FGF5 - FGFR1 | FGF | Secreted Signaling | PMC: 4393358 |
| A-PEP.SCGN/ADRA2C | FGF5 | FGFR1 | 0.007395602 | 0 | FGF5_FGFR1 | FGF5 - FGFR1 | FGF | Secreted Signaling | PMC: 4393358 |
| C-LTMR.CDH9 | FGF5 | FGFR1 | 0.013814284 | 0 | FGF5_FGFR1 | FGF5 - FGFR1 | FGF | Secreted Signaling | PMC: 4393358 |
| C-PEP.TAC1/CACNG5 | FGF5 | FGFR1 | 0.008871052 | 0 | FGF5_FGFR1 | FGF5 - FGFR1 | FGF | Secreted Signaling | PMC: 4393358 |
| C-LTMR.CDH9 | FGF10 | FGFR1 | 0.035598393 | 0 | FGF10_FGFR1 | FGF10 - FGFR1 | FGF | Secreted Signaling | PMC: 4393358 |
| C-NP.MRGPRX1/MRGPRX4 | FGF10 | FGFR1 | 0.013004743 | 0 | FGF10_FGFR1 | FGF10 - FGFR1 | FGF | Secreted Signaling | PMC: 4393358 |
| C-NP.SST/CCK | FGF10 | FGFR1 | 0.009250863 | 0.02 | FGF10_FGFR1 | FGF10 - FGFR1 | FGF | Secreted Signaling | PMC: 4393358 |
| A-PEP.NTRK3/S100A16 | FGF18 | FGFR1 | 0.000723778 | 0 | FGF18_FGFR1 | FGF18 - FGFR1 | FGF | Secreted Signaling | PMC: 4393358 |
| A-PEP.SCGN/ADRA2C | FGF18 | FGFR1 | 0.000605923 | 0.01 | FGF18_FGFR1 | FGF18 - FGFR1 | FGF | Secreted Signaling | PMC: 4393358 |
| A-PEP.KIT | FGF9 | FGFR1 | 0.005947646 | 0.03 | FGF9_FGFR1 | FGF9 - FGFR1 | FGF | Secreted Signaling | PMC: 4393358 |
| A-PEP.NTRK3/S100A16 | FGF9 | FGFR1 | 0.011390542 | 0 | FGF9_FGFR1 | FGF9 - FGFR1 | FGF | Secreted Signaling | PMC: 4393358 |
| A-PEP.SCGN/ADRA2C | FGF9 | FGFR1 | 0.010100689 | 0 | FGF9_FGFR1 | FGF9 - FGFR1 | FGF | Secreted Signaling | PMC: 4393358 |
| C-LTMR.CDH9 | FGF9 | FGFR1 | 0.01561052 | 0 | FGF9_FGFR1 | FGF9 - FGFR1 | FGF | Secreted Signaling | PMC: 4393358 |
| A-PEP.SCGN/ADRA2C | PDGFA | PDGFRA | 0.00048802 | 0.04 | PDGFA_PDGFRA | PDGFA - PDGFRA | PDGF | Secreted Signaling | PMID: 15207812 |
| A-PEP.KIT | PDGFB | PDGFRA | 7.64171E-05 | 0.03 | PDGFB_PDGFRA | PDGFB - PDGFRA | PDGF | Secreted Signaling | PMID: 15207812 |
| A-PEP.SCGN/ADRA2C | PDGFB | PDGFRA | 6.87433E-05 | 0.01 | PDGFB_PDGFRA | PDGFB - PDGFRA | PDGF | Secreted Signaling | PMID: 15207812 |
| C-PEP.ADORA2B | PDGFB | PDGFRA | 9.3522E-05 | 0.02 | PDGFB_PDGFRA | PDGFB - PDGFRA | PDGF | Secreted Signaling | PMID: 15207812 |
| C-PEP.TAC1/CHRNA3 | PDGFB | PDGFRA | 7.2885E-05 | 0.03 | PDGFB_PDGFRA | PDGFB - PDGFRA | PDGF | Secreted Signaling | PMID: 15207812 |

| | | | | | | | | | |
|---|---|---|---|---|---|---|---|---|---|
| C-PEP.ADORA2B | PDGFB | PDGFRB | 0.000215614 | 0.02 | PDGFB_PDGFRB | PDGFB - PDGFRB | PDGF | Secreted Signaling | PMID: 15207812 |
| C-LTMR.CDH9 | PDGFC | PDGFRA | 9.60306E-06 | 0 | PDGFC_PDGFRA | PDGFC - PDGFRA | PDGF | Secreted Signaling | PMID: 15207812 |
| A-PEP.SCGN/ADRA2C | PDGFD | PDGFRB | 2.06086E-05 | 0 | PDGFD_PDGFRB | PDGFD - PDGFRB | PDGF | Secreted Signaling | PMID: 15207812 |
| C-PEP.ADORA2B | PDGFD | PDGFRB | 5.7683E-05 | 0 | PDGFD_PDGFRB | PDGFD - PDGFRB | PDGF | Secreted Signaling | PMID: 15207812 |
| A-PEP.CHRNA7/SLC18A3 | VEGFA | KDR | 0.000233103 | 0 | VEGFA_VEGFR2 | VEGFA - VEGFR2 | VEGF | Secreted Signaling | KEGG: hsa04370; PMID: 16633338 |
| A-PEP.KIT | VEGFA | KDR | 0.000404065 | 0 | VEGFA_VEGFR2 | VEGFA - VEGFR2 | VEGF | Secreted Signaling | KEGG: hsa04370; PMID: 16633338 |
| A-PEP.NTRK3/S100A16 | VEGFA | KDR | 0.000122368 | 0 | VEGFA_VEGFR2 | VEGFA - VEGFR2 | VEGF | Secreted Signaling | KEGG: hsa04370; PMID: 16633338 |
| A-PEP.SCGN/ADRA2C | VEGFA | KDR | 0.000598131 | 0 | VEGFA_VEGFR2 | VEGFA - VEGFR2 | VEGF | Secreted Signaling | KEGG: hsa04370; PMID: 16633338 |
| C-PEP.ADORA2B | VEGFA | KDR | 0.000789118 | 0 | VEGFA_VEGFR2 | VEGFA - VEGFR2 | VEGF | Secreted Signaling | KEGG: hsa04370; PMID: 16633338 |
| C-PEP.TAC1/CACNG5 | VEGFA | KDR | 0.00063345 | 0 | VEGFA_VEGFR2 | VEGFA - VEGFR2 | VEGF | Secreted Signaling | KEGG: hsa04370; PMID: 16633338 |
| C-PEP.TAC1/CHRNA3 | VEGFA | KDR | 0.000489603 | 0 | VEGFA_VEGFR2 | VEGFA - VEGFR2 | VEGF | Secreted Signaling | KEGG: hsa04370; PMID: 16633338 |
| A-PEP.KIT | VEGFC | KDR | 1.13837E-05 | 0 | VEGFC_VEGFR2 | VEGFC - VEGFR2 | VEGF | Secreted Signaling | KEGG: hsa04370; PMID: 16633338 |
| A-PEP.SCGN/ADRA2C | VEGFC | KDR | 0.000217253 | 0 | VEGFC_VEGFR2 | VEGFC - VEGFR2 | VEGF | Secreted Signaling | KEGG: hsa04370; PMID: 16633338 |
| C-PEP.ADORA2B | VEGFC | KDR | 0.000699004 | 0 | VEGFC_VEGFR2 | VEGFC - VEGFR2 | VEGF | Secreted Signaling | KEGG: hsa04370; PMID: 16633338 |
| A-PEP.NTRK3/S100A16 | CCL2 | ACKR1 | 0.001759537 | 0 | CCL2_ACKR1 | CCL2 - ACKR1 | CCL | Secreted Signaling | PMID: 26740381 |
| C-LTMR.CDH9 | CCL2 | ACKR1 | 0.001872795 | 0.04 | CCL2_ACKR1 | CCL2 - ACKR1 | CCL | Secreted Signaling | PMID: 26740381 |
| C-PEP.ADORA2B | MIF | CD74_CD44 | 0.017403934 | 0 | MIF_CD74_CD44 | MIF - (CD74+CD44) | MIF | Secreted Signaling | PMID: 29637711; PMID: 26175090 |
| A-PEP.SCGN/ADRA2C | CX3CL1 | CX3CR1 | 0.000358976 | 0 | CX3CL1_CX3CR1 | CX3CL1 - CX3CR1 | CX3C | Secreted Signaling | KEGG: hsa04060 |
| A-PEP.SCGN/ADRA2C | TNF | TNFRSF1A | 4.17711E-05 | 0 | TNF_TNFRSF1A | TNF - TNFRSF1A | TNF | Secreted Signaling | KEGG: hsa04060 |
| C-PEP.TAC1/CACNG5 | TNF | TNFRSF1A | 4.64145E-05 | 0 | TNF_TNFRSF1A | TNF - TNFRSF1A | TNF | Secreted Signaling | KEGG: hsa04060 |
| A-PEP.NTRK3/S100A16 | SPP1 | CD44 | 0.094508586 | 0 | SPP1_CD44 | SPP1 - CD44 | SPP1 | Secreted Signaling | PMID: 21907263 |
| A-PEP.NTRK3/S100A16 | SPP1 | ITGAV_ITGB1 | 0.042783047 | 0 | SPP1_ITGAV_ITGB1 | SPP1 - (ITGAV+ITGB1) | SPP1 | Secreted Signaling | PMID: 21907263 |
| C-NP.MRGPRX1/MRGPRX4 | SPP1 | ITGAV_ITGB1 | 0.017887865 | 0.01 | SPP1_ITGAV_ITGB1 | SPP1 - (ITGAV+ITGB1) | SPP1 | Secreted Signaling | PMID: 21907263 |

| | | | | | | | | | |
|---|---|---|---|---|---|---|---|---|---|
| C-NP.SST/CCK | SPP1 | ITGAV_ITGB1 | 0.020596086 | 0 | SPP1_ITGAV_ITGB1 | SPP1 - (ITGAV+ITGB1) | SPP1 | Secreted Signaling | PMID: 21907263 |
| A-PEP.NTRK3/S100A16 | SPP1 | ITGAV_ITGB5 | 0.004774877 | 0 | SPP1_ITGAV_ITGB5 | SPP1 - (ITGAV+ITGB5) | SPP1 | Secreted Signaling | PMID: 21907263 |
| C-LTMR.CDH9 | SPP1 | ITGAV_ITGB5 | 0.002026802 | 0.03 | SPP1_ITGAV_ITGB5 | SPP1 - (ITGAV+ITGB5) | SPP1 | Secreted Signaling | PMID: 21907263 |
| C-NP.MRGPRX1/MRGPRX4 | SPP1 | ITGAV_ITGB5 | 0.001951321 | 0.03 | SPP1_ITGAV_ITGB5 | SPP1 - (ITGAV+ITGB5) | SPP1 | Secreted Signaling | PMID: 21907263 |
| C-NP.SST/CCK | SPP1 | ITGAV_ITGB5 | 0.002252284 | 0.01 | SPP1_ITGAV_ITGB5 | SPP1 - (ITGAV+ITGB5) | SPP1 | Secreted Signaling | PMID: 21907263 |
| C-PEP.ADORA2B | PTN | PTPRZ1 | 0.001269889 | 0 | PTN_PTPRZ1 | PTN - PTPRZ1 | PTN | Secreted Signaling | PMID: 28356350; PMID: 25620911 |
| A-PEP.CHRNA7/SLC18A3 | PTN | SDC2 | 4.53105E-05 | 0 | PTN_SDC2 | PTN - SDC2 | PTN | Secreted Signaling | PMID: 28356350; PMID: 25620911 |
| A-PEP.SCGN/ADRA2C | PTN | SDC2 | 1.97342E-05 | 0.04 | PTN_SDC2 | PTN - SDC2 | PTN | Secreted Signaling | PMID: 28356350; PMID: 25620911 |
| C-LTMR.CDH9 | PTN | SDC2 | 6.20857E-05 | 0 | PTN_SDC2 | PTN - SDC2 | PTN | Secreted Signaling | PMID: 28356350; PMID: 25620911 |
| C-PEP.ADORA2B | PTN | SDC2 | 0.000169873 | 0 | PTN_SDC2 | PTN - SDC2 | PTN | Secreted Signaling | PMID: 28356350; PMID: 25620911 |
| C-PEP.TAC1/CHRNA3 | PTN | SDC2 | 4.08038E-05 | 0.01 | PTN_SDC2 | PTN - SDC2 | PTN | Secreted Signaling | PMID: 28356350; PMID: 25620911 |
| A-PEP.CHRNA7/SLC18A3 | PTN | SDC3 | 0.000595926 | 0.01 | PTN_SDC3 | PTN - SDC3 | PTN | Secreted Signaling | PMID: 28356350; PMID: 25620911 |
| C-LTMR.CDH9 | PTN | SDC3 | 0.000816389 | 0.04 | PTN_SDC3 | PTN - SDC3 | PTN | Secreted Signaling | PMID: 28356350; PMID: 25620911 |
| C-PEP.ADORA2B | PTN | SDC3 | 0.002230798 | 0 | PTN_SDC3 | PTN - SDC3 | PTN | Secreted Signaling | PMID: 28356350; PMID: 25620911 |
| C-PEP.TAC1/CHRNA3 | PTN | SDC3 | 0.000536683 | 0.04 | PTN_SDC3 | PTN - SDC3 | PTN | Secreted Signaling | PMID: 28356350; PMID: 25620911 |
| C-PEP.ADORA2B | PTN | SDC4 | 8.45441E-05 | 0 | PTN_SDC4 | PTN - SDC4 | PTN | Secreted Signaling | PMID: 28356350; PMID: 25620911 |
| A-PEP.CHRNA7/SLC18A3 | PTN | NCL | 0.001801157 | 0.01 | PTN_NCL | PTN - NCL | PTN | Secreted Signaling | PMID: 28356350; PMID: 25620911 |
| C-PEP.ADORA2B | PTN | NCL | 0.006720242 | 0 | PTN_NCL | PTN - NCL | PTN | Secreted Signaling | PMID: 28356350; PMID: 25620911 |
| C-NP.SST/CCK | CCK | CCKAR | 0.002326524 | 0 | CCK_CCKAR | CCK - CCKAR | CCK | Secreted Signaling | KEGG: hsa04080 |
| C-NP.MRGPRX1/MRGPRX4 | PENK | OPRK1 | 5.28254E-07 | 0 | PENK_OPRK1 | PENK - OPRK1 | OPIOID | Secreted Signaling | KEGG: hsa04080 |
| C-NP.MRGPRX1/MRGPRX4 | PENK | OPRM1 | 0.000289106 | 0 | PENK_OPRM1 | PENK - OPRM1 | OPIOID | Secreted Signaling | KEGG: hsa04080 |

| | | | | | | | | | |
|---|---|---|---|---|---|---|---|---|---|
| A-PEP.SCGN/ADRA2C | POMC | OPRK1 | 4.66117E-07 | 0 | POMC_OPRK1 | POMC - OPRK1 | OPIOID | Secreted Signaling | KEGG: hsa04080 |
| A-PEP.SCGN/ADRA2C | POMC | OPRM1 | 0.000255108 | 0 | POMC_OPRM1 | POMC - OPRM1 | OPIOID | Secreted Signaling | KEGG: hsa04080 |
| A-PEP.CHRNA7/SLC18A3 | TAC1 | TACR2 | 0.00270149 | 0 | TAC1_TACR2 | TAC1 - TACR2 | TAC | Secreted Signaling | KEGG: hsa04080 |
| C-PEP.ADORA2B | TAC1 | TACR2 | 0.006138333 | 0 | TAC1_TACR2 | TAC1 - TACR2 | TAC | Secreted Signaling | KEGG: hsa04080 |
| C-PEP.TAC1/CACNG5 | TAC1 | TACR2 | 0.003951982 | 0 | TAC1_TACR2 | TAC1 - TACR2 | TAC | Secreted Signaling | KEGG: hsa04080 |
| C-PEP.TAC1/CHRNA3 | TAC1 | TACR2 | 0.003757506 | 0 | TAC1_TACR2 | TAC1 - TACR2 | TAC | Secreted Signaling | KEGG: hsa04080 |
| A-PEP.CHRNA7/SLC18A3 | KITLG | KIT | 0.003890444 | 0 | KITL_KIT | KITL - KIT | KIT | Secreted Signaling | KEGG: hsa04080 |
| A-PEP.KIT | KITLG | KIT | 0.007119123 | 0 | KITL_KIT | KITL - KIT | KIT | Secreted Signaling | KEGG: hsa04080 |
| A-PEP.NTRK3/S100A16 | KITLG | KIT | 0.013221321 | 0 | KITL_KIT | KITL - KIT | KIT | Secreted Signaling | KEGG: hsa04080 |
| A-PEP.SCGN/ADRA2C | KITLG | KIT | 0.007387684 | 0 | KITL_KIT | KITL - KIT | KIT | Secreted Signaling | KEGG: hsa04080 |
| C-NP.MRGPRX1/MRGPRX4 | SEMA3B | NRP1_PLXNA1 | 0.000337092 | 0.02 | SEMA3B_NRP1_PLXNA1 | SEMA3B - (NRP1+PLXNA1) | SEMA3 | Secreted Signaling | PMID: 27533782 |
| C-NP.SST/CCK | SEMA3B | NRP1_PLXNA1 | 0.000520666 | 0.03 | SEMA3B_NRP1_PLXNA1 | SEMA3B - (NRP1+PLXNA1) | SEMA3 | Secreted Signaling | PMID: 27533782 |
| C-PEP.TAC1/CACNG5 | SEMA3B | NRP1_PLXNA1 | 0.000856383 | 0.02 | SEMA3B_NRP1_PLXNA1 | SEMA3B - (NRP1+PLXNA1) | SEMA3 | Secreted Signaling | PMID: 27533782 |
| C-NP.MRGPRX1/MRGPRX4 | SEMA3B | NRP1_PLXNA2 | 0.000551229 | 0.01 | SEMA3B_NRP1_PLXNA2 | SEMA3B - (NRP1+PLXNA2) | SEMA3 | Secreted Signaling | PMID: 27533782 |
| C-NP.SST/CCK | SEMA3B | NRP1_PLXNA2 | 0.00085132 | 0.01 | SEMA3B_NRP1_PLXNA2 | SEMA3B - (NRP1+PLXNA2) | SEMA3 | Secreted Signaling | PMID: 27533782 |
| C-PEP.TAC1/CACNG5 | SEMA3B | NRP1_PLXNA2 | 0.001399936 | 0.01 | SEMA3B_NRP1_PLXNA2 | SEMA3B - (NRP1+PLXNA2) | SEMA3 | Secreted Signaling | PMID: 27533782 |
| C-NP.MRGPRX1/MRGPRX4 | SEMA3B | NRP1_PLXNA3 | 0.000747681 | 0 | SEMA3B_NRP1_PLXNA3 | SEMA3B - (NRP1+PLXNA3) | SEMA3 | Secreted Signaling | PMID: 27533782 |
| C-NP.SST/CCK | SEMA3B | NRP1_PLXNA3 | 0.001154597 | 0 | SEMA3B_NRP1_PLXNA3 | SEMA3B - (NRP1+PLXNA3) | SEMA3 | Secreted Signaling | PMID: 27533782 |
| C-PEP.TAC1/CACNG5 | SEMA3B | NRP1_PLXNA3 | 0.001898283 | 0.01 | SEMA3B_NRP1_PLXNA3 | SEMA3B - (NRP1+PLXNA3) | SEMA3 | Secreted Signaling | PMID: 27533782 |
| C-NP.MRGPRX1/MRGPRX4 | SEMA3B | NRP1_PLXNA4 | 0.00085204 | 0 | SEMA3B_NRP1_PLXNA4 | SEMA3B - (NRP1+PLXNA4) | SEMA3 | Secreted Signaling | PMID: 27533782 |
| C-NP.SST/CCK | SEMA3B | NRP1_PLXNA4 | 0.001315677 | 0 | SEMA3B_NRP1_PLXNA4 | SEMA3B - (NRP1+PLXNA4) | SEMA3 | Secreted Signaling | PMID: 27533782 |

| | | | | | | | | | |
|---|---|---|---|---|---|---|---|---|---|
| C-PEP.TAC1/CACNG5 | SEMA3B | NRP1_PLXNA4 | 0.002162892 | 0 | SEMA3B_NRP1_PLXNA4 | SEMA3B - (NRP1+PLXNA4) | SEMA3 | Secreted Signaling | PMID: 27533782 |
| C-NP.MRGPRX1/MRGPRX4 | SEMA3C | NRP1_PLXNA1 | 0.000582584 | 0 | SEMA3C_NRP1_PLXNA1 | SEMA3C - (NRP1+PLXNA1) | SEMA3 | Secreted Signaling | PMID: 27533782 |
| C-NP.MRGPRX1/MRGPRX4 | SEMA3C | NRP1_PLXNA2 | 0.000952521 | 0 | SEMA3C_NRP1_PLXNA2 | SEMA3C - (NRP1+PLXNA2) | SEMA3 | Secreted Signaling | PMID: 27533782 |
| C-NP.MRGPRX1/MRGPRX4 | SEMA3C | NRP1_PLXNA3 | 0.001291804 | 0 | SEMA3C_NRP1_PLXNA3 | SEMA3C - (NRP1+PLXNA3) | SEMA3 | Secreted Signaling | PMID: 27533782 |
| C-NP.MRGPRX1/MRGPRX4 | SEMA3C | NRP1_PLXNA4 | 0.001471999 | 0 | SEMA3C_NRP1_PLXNA4 | SEMA3C - (NRP1+PLXNA4) | SEMA3 | Secreted Signaling | PMID: 27533782 |
| A-PEP.KIT | SEMA3D | NRP1_PLXNA1 | 0.000344818 | 0 | SEMA3D_NRP1_PLXNA1 | SEMA3D - (NRP1+PLXNA1) | SEMA3 | Secreted Signaling | PMID: 27533782 |
| C-LTMR.CDH9 | SEMA3D | NRP1_PLXNA1 | 0.00053525 | 0 | SEMA3D_NRP1_PLXNA1 | SEMA3D - (NRP1+PLXNA1) | SEMA3 | Secreted Signaling | PMID: 27533782 |
| A-PEP.KIT | SEMA3D | NRP1_PLXNA2 | 0.00056386 | 0 | SEMA3D_NRP1_PLXNA2 | SEMA3D - (NRP1+PLXNA2) | SEMA3 | Secreted Signaling | PMID: 27533782 |
| C-LTMR.CDH9 | SEMA3D | NRP1_PLXNA2 | 0.000875157 | 0 | SEMA3D_NRP1_PLXNA2 | SEMA3D - (NRP1+PLXNA2) | SEMA3 | Secreted Signaling | PMID: 27533782 |
| A-PEP.KIT | SEMA3D | NRP1_PLXNA3 | 0.00076481 | 0 | SEMA3D_NRP1_PLXNA3 | SEMA3D - (NRP1+PLXNA3) | SEMA3 | Secreted Signaling | PMID: 27533782 |
| C-LTMR.CDH9 | SEMA3D | NRP1_PLXNA3 | 0.001186916 | 0 | SEMA3D_NRP1_PLXNA3 | SEMA3D - (NRP1+PLXNA3) | SEMA3 | Secreted Signaling | PMID: 27533782 |
| A-PEP.KIT | SEMA3D | NRP1_PLXNA4 | 0.000871557 | 0 | SEMA3D_NRP1_PLXNA4 | SEMA3D - (NRP1+PLXNA4) | SEMA3 | Secreted Signaling | PMID: 27533782 |
| C-LTMR.CDH9 | SEMA3D | NRP1_PLXNA4 | 0.001352499 | 0 | SEMA3D_NRP1_PLXNA4 | SEMA3D - (NRP1+PLXNA4) | SEMA3 | Secreted Signaling | PMID: 27533782 |
| C-NP.MRGPRX1/MRGPRX4 | SEMA3B | NRP2_PLXNA1 | 0.000638177 | 0 | SEMA3B_NRP2_PLXNA1 | SEMA3B - (NRP2+PLXNA1) | SEMA3 | Secreted Signaling | PMID: 27533782 |
| C-NP.SST/CCK | SEMA3B | NRP2_PLXNA1 | 0.000985555 | 0 | SEMA3B_NRP2_PLXNA1 | SEMA3B - (NRP2+PLXNA1) | SEMA3 | Secreted Signaling | PMID: 27533782 |
| C-PEP.ADORA2B | SEMA3B | NRP2_PLXNA1 | 0.000482693 | 0.03 | SEMA3B_NRP2_PLXNA1 | SEMA3B - (NRP2+PLXNA1) | SEMA3 | Secreted Signaling | PMID: 27533782 |
| C-PEP.TAC1/CACNG5 | SEMA3B | NRP2_PLXNA1 | 0.001620536 | 0 | SEMA3B_NRP2_PLXNA1 | SEMA3B - (NRP2+PLXNA1) | SEMA3 | Secreted Signaling | PMID: 27533782 |
| C-NP.MRGPRX1/MRGPRX4 | SEMA3B | NRP2_PLXNA2 | 0.001043377 | 0 | SEMA3B_NRP2_PLXNA2 | SEMA3B - (NRP2+PLXNA2) | SEMA3 | Secreted Signaling | PMID: 27533782 |
| C-NP.SST/CCK | SEMA3B | NRP2_PLXNA2 | 0.001610963 | 0 | SEMA3B_NRP2_PLXNA2 | SEMA3B - (NRP2+PLXNA2) | SEMA3 | Secreted Signaling | PMID: 27533782 |
| C-PEP.ADORA2B | SEMA3B | NRP2_PLXNA2 | 0.00078925 | 0 | SEMA3B_NRP2_PLXNA2 | SEMA3B - (NRP2+PLXNA2) | SEMA3 | Secreted Signaling | PMID: 27533782 |
| C-PEP.TAC1/CACNG5 | SEMA3B | NRP2_PLXNA2 | 0.00264782 | 0 | SEMA3B_NRP2_PLXNA2 | SEMA3B - (NRP2+PLXNA2) | SEMA3 | Secreted Signaling | PMID: 27533782 |

| | | | | | | | | | |
|---|---|---|---|---|---|---|---|---|---|
| C-NP.MRGPRX1/MRGPRX4 | SEMA3B | NRP2_PLXNA3 | 0.001414977 | 0 | SEMA3B_NRP2_PLXNA3 | SEMA3B - (NRP2+PLXNA3) | SEMA3 | Secreted Signaling | PMID: 27533782 |
| C-NP.SST/CCK | SEMA3B | NRP2_PLXNA3 | 0.002184267 | 0 | SEMA3B_NRP2_PLXNA3 | SEMA3B - (NRP2+PLXNA3) | SEMA3 | Secreted Signaling | PMID: 27533782 |
| C-PEP.ADORA2B | SEMA3B | NRP2_PLXNA3 | 0.001070439 | 0 | SEMA3B_NRP2_PLXNA3 | SEMA3B - (NRP2+PLXNA3) | SEMA3 | Secreted Signaling | PMID: 27533782 |
| C-PEP.TAC1/CACNG5 | SEMA3B | NRP2_PLXNA3 | 0.00358879 | 0 | SEMA3B_NRP2_PLXNA3 | SEMA3B - (NRP2+PLXNA3) | SEMA3 | Secreted Signaling | PMID: 27533782 |
| C-LTMR.CDH9 | SEMA3B | NRP2_PLXNA4 | 0.001444684 | 0.04 | SEMA3B_NRP2_PLXNA4 | SEMA3B - (NRP2+PLXNA4) | SEMA3 | Secreted Signaling | PMID: 27533782 |
| C-NP.MRGPRX1/MRGPRX4 | SEMA3B | NRP2_PLXNA4 | 0.001612325 | 0 | SEMA3B_NRP2_PLXNA4 | SEMA3B - (NRP2+PLXNA4) | SEMA3 | Secreted Signaling | PMID: 27533782 |
| C-NP.SST/CCK | SEMA3B | NRP2_PLXNA4 | 0.002488641 | 0 | SEMA3B_NRP2_PLXNA4 | SEMA3B - (NRP2+PLXNA4) | SEMA3 | Secreted Signaling | PMID: 27533782 |
| C-PEP.ADORA2B | SEMA3B | NRP2_PLXNA4 | 0.001219793 | 0 | SEMA3B_NRP2_PLXNA4 | SEMA3B - (NRP2+PLXNA4) | SEMA3 | Secreted Signaling | PMID: 27533782 |
| C-PEP.TAC1/CACNG5 | SEMA3B | NRP2_PLXNA4 | 0.004088081 | 0 | SEMA3B_NRP2_PLXNA4 | SEMA3B - (NRP2+PLXNA4) | SEMA3 | Secreted Signaling | PMID: 27533782 |
| C-NP.MRGPRX1/MRGPRX4 | SEMA3C | NRP2_PLXNA1 | 0.001102696 | 0 | SEMA3C_NRP2_PLXNA1 | SEMA3C - (NRP2+PLXNA1) | SEMA3 | Secreted Signaling | PMID: 27533782 |
| C-PEP.TAC1/CACNG5 | SEMA3C | NRP2_PLXNA1 | 0.000433757 | 0.04 | SEMA3C_NRP2_PLXNA1 | SEMA3C - (NRP2+PLXNA1) | SEMA3 | Secreted Signaling | PMID: 27533782 |
| C-NP.MRGPRX1/MRGPRX4 | SEMA3C | NRP2_PLXNA2 | 0.001802305 | 0 | SEMA3C_NRP2_PLXNA2 | SEMA3C - (NRP2+PLXNA2) | SEMA3 | Secreted Signaling | PMID: 27533782 |
| C-NP.SST/CCK | SEMA3C | NRP2_PLXNA2 | 0.000553139 | 0.02 | SEMA3C_NRP2_PLXNA2 | SEMA3C - (NRP2+PLXNA2) | SEMA3 | Secreted Signaling | PMID: 27533782 |
| C-PEP.TAC1/CACNG5 | SEMA3C | NRP2_PLXNA2 | 0.000709257 | 0 | SEMA3C_NRP2_PLXNA2 | SEMA3C - (NRP2+PLXNA2) | SEMA3 | Secreted Signaling | PMID: 27533782 |
| C-NP.MRGPRX1/MRGPRX4 | SEMA3C | NRP2_PLXNA3 | 0.002443536 | 0 | SEMA3C_NRP2_PLXNA3 | SEMA3C - (NRP2+PLXNA3) | SEMA3 | Secreted Signaling | PMID: 27533782 |
| C-NP.SST/CCK | SEMA3C | NRP2_PLXNA3 | 0.000750271 | 0.01 | SEMA3C_NRP2_PLXNA3 | SEMA3C - (NRP2+PLXNA3) | SEMA3 | Secreted Signaling | PMID: 27533782 |
| C-PEP.TAC1/CACNG5 | SEMA3C | NRP2_PLXNA3 | 0.000961975 | 0.01 | SEMA3C_NRP2_PLXNA3 | SEMA3C - (NRP2+PLXNA3) | SEMA3 | Secreted Signaling | PMID: 27533782 |
| C-NP.MRGPRX1/MRGPRX4 | SEMA3C | NRP2_PLXNA4 | 0.002783938 | 0 | SEMA3C_NRP2_PLXNA4 | SEMA3C - (NRP2+PLXNA4) | SEMA3 | Secreted Signaling | PMID: 27533782 |
| C-NP.SST/CCK | SEMA3C | NRP2_PLXNA4 | 0.000854991 | 0.01 | SEMA3C_NRP2_PLXNA4 | SEMA3C - (NRP2+PLXNA4) | SEMA3 | Secreted Signaling | PMID: 27533782 |
| C-PEP.TAC1/CACNG5 | SEMA3C | NRP2_PLXNA4 | 0.001096211 | 0 | SEMA3C_NRP2_PLXNA4 | SEMA3C - (NRP2+PLXNA4) | SEMA3 | Secreted Signaling | PMID: 27533782 |
| A-PEP.KIT | SEMA3D | NRP2_PLXNA1 | 0.000652798 | 0 | SEMA3D_NRP2_PLXNA1 | SEMA3D - (NRP2+PLXNA1) | SEMA3 | Secreted Signaling | PMID: 27533782 |

| | | | | | | | | | |
|---|---|---|---|---|---|---|---|---|---|
| C-LTMR.CDH9 | SEMA3D | NRP2_PLXNA1 | 0.001013147 | 0 | SEMA3D_NRP2_PLXNA1 | SEMA3D - (NRP2+PLXNA1) | SEMA3 | Secreted Signaling | PMID: 27533782 |
| A-PEP.KIT | SEMA3D | NRP2_PLXNA2 | 0.001067273 | 0 | SEMA3D_NRP2_PLXNA2 | SEMA3D - (NRP2+PLXNA2) | SEMA3 | Secreted Signaling | PMID: 27533782 |
| C-LTMR.CDH9 | SEMA3D | NRP2_PLXNA2 | 0.001656035 | 0 | SEMA3D_NRP2_PLXNA2 | SEMA3D - (NRP2+PLXNA2) | SEMA3 | Secreted Signaling | PMID: 27533782 |
| A-PEP.KIT | SEMA3D | NRP2_PLXNA3 | 0.001447371 | 0 | SEMA3D_NRP2_PLXNA3 | SEMA3D - (NRP2+PLXNA3) | SEMA3 | Secreted Signaling | PMID: 27533782 |
| C-LTMR.CDH9 | SEMA3D | NRP2_PLXNA3 | 0.002245343 | 0 | SEMA3D_NRP2_PLXNA3 | SEMA3D - (NRP2+PLXNA3) | SEMA3 | Secreted Signaling | PMID: 27533782 |
| A-PEP.KIT | SEMA3D | NRP2_PLXNA4 | 0.001649229 | 0 | SEMA3D_NRP2_PLXNA4 | SEMA3D - (NRP2+PLXNA4) | SEMA3 | Secreted Signaling | PMID: 27533782 |
| C-LTMR.CDH9 | SEMA3D | NRP2_PLXNA4 | 0.002558206 | 0 | SEMA3D_NRP2_PLXNA4 | SEMA3D - (NRP2+PLXNA4) | SEMA3 | Secreted Signaling | PMID: 27533782 |
| A-PEP.CHRNA7/SLC18A3 | SEMA3F | NRP2_PLXNA1 | 0.000592146 | 0 | SEMA3F_NRP2_PLXNA1 | SEMA3F - (NRP2+PLXNA1) | SEMA3 | Secreted Signaling | PMID: 27533782 |
| A-PEP.KIT | SEMA3F | NRP2_PLXNA1 | 0.00060156 | 0 | SEMA3F_NRP2_PLXNA1 | SEMA3F - (NRP2+PLXNA1) | SEMA3 | Secreted Signaling | PMID: 27533782 |
| A-PEP.SCGN/ADRA2C | SEMA3F | NRP2_PLXNA1 | 0.00101426 | 0 | SEMA3F_NRP2_PLXNA1 | SEMA3F - (NRP2+PLXNA1) | SEMA3 | Secreted Signaling | PMID: 27533782 |
| A-PEP.CHRNA7/SLC18A3 | SEMA3F | NRP2_PLXNA2 | 0.000968149 | 0 | SEMA3F_NRP2_PLXNA2 | SEMA3F - (NRP2+PLXNA2) | SEMA3 | Secreted Signaling | PMID: 27533782 |
| A-PEP.KIT | SEMA3F | NRP2_PLXNA2 | 0.000983535 | 0 | SEMA3F_NRP2_PLXNA2 | SEMA3F - (NRP2+PLXNA2) | SEMA3 | Secreted Signaling | PMID: 27533782 |
| A-PEP.NTRK3/S100A16 | SEMA3F | NRP2_PLXNA2 | 0.000200027 | 0.02 | SEMA3F_NRP2_PLXNA2 | SEMA3F - (NRP2+PLXNA2) | SEMA3 | Secreted Signaling | PMID: 27533782 |
| A-PEP.SCGN/ADRA2C | SEMA3F | NRP2_PLXNA2 | 0.001657853 | 0 | SEMA3F_NRP2_PLXNA2 | SEMA3F - (NRP2+PLXNA2) | SEMA3 | Secreted Signaling | PMID: 27533782 |
| C-LTMR.CDH9 | SEMA3F | NRP2_PLXNA2 | 0.000486454 | 0.02 | SEMA3F_NRP2_PLXNA2 | SEMA3F - (NRP2+PLXNA2) | SEMA3 | Secreted Signaling | PMID: 27533782 |
| A-PEP.CHRNA7/SLC18A3 | SEMA3F | NRP2_PLXNA3 | 0.001312991 | 0 | SEMA3F_NRP2_PLXNA3 | SEMA3F - (NRP2+PLXNA3) | SEMA3 | Secreted Signaling | PMID: 27533782 |
| A-PEP.KIT | SEMA3F | NRP2_PLXNA3 | 0.001333851 | 0 | SEMA3F_NRP2_PLXNA3 | SEMA3F - (NRP2+PLXNA3) | SEMA3 | Secreted Signaling | PMID: 27533782 |
| A-PEP.NTRK3/S100A16 | SEMA3F | NRP2_PLXNA3 | 0.000271348 | 0.02 | SEMA3F_NRP2_PLXNA3 | SEMA3F - (NRP2+PLXNA3) | SEMA3 | Secreted Signaling | PMID: 27533782 |
| A-PEP.SCGN/ADRA2C | SEMA3F | NRP2_PLXNA3 | 0.002247806 | 0 | SEMA3F_NRP2_PLXNA3 | SEMA3F - (NRP2+PLXNA3) | SEMA3 | Secreted Signaling | PMID: 27533782 |
| C-LTMR.CDH9 | SEMA3F | NRP2_PLXNA3 | 0.000659836 | 0.02 | SEMA3F_NRP2_PLXNA3 | SEMA3F - (NRP2+PLXNA3) | SEMA3 | Secreted Signaling | PMID: 27533782 |
| A-PEP.CHRNA7/SLC18A3 | SEMA3F | NRP2_PLXNA4 | 0.001496136 | 0 | SEMA3F_NRP2_PLXNA4 | SEMA3F - (NRP2+PLXNA4) | SEMA3 | Secreted Signaling | PMID: 27533782 |
| A-PEP.KIT | SEMA3F | NRP2_PLXNA4 | 0.001519901 | 0 | SEMA3F_NRP2_PLXNA4 | SEMA3F - (NRP2+PLXNA4) | SEMA3 | Secreted Signaling | PMID: 27533782 |

| | | | | | | | | | |
|---|---|---|---|---|---|---|---|---|---|
| A-PEP.NTRK3/S100A16 | SEMA3F | NRP2_PLXNA4 | 0.000309243 | 0 | SEMA3F_NRP2_PLXNA4 | SEMA3F - (NRP2+PLXNA4) | SEMA3 | Secreted Signaling | PMID: 27533782 |
| A-PEP.SCGN/ADRA2C | SEMA3F | NRP2_PLXNA4 | 0.002561011 | 0 | SEMA3F_NRP2_PLXNA4 | SEMA3F - (NRP2+PLXNA4) | SEMA3 | Secreted Signaling | PMID: 27533782 |
| C-LTMR.CDH9 | SEMA3F | NRP2_PLXNA4 | 0.000751943 | 0.02 | SEMA3F_NRP2_PLXNA4 | SEMA3F - (NRP2+PLXNA4) | SEMA3 | Secreted Signaling | PMID: 27533782 |
| C-NP.MRGPRX1/MRGPRX4 | SEMA3G | NRP2_PLXNA1 | 0.006321496 | 0 | SEMA3G_NRP2_PLXNA1 | SEMA3G - (NRP2+PLXNA1) | SEMA3 | Secreted Signaling | PMID: 27533782 |
| C-NP.SST/CCK | SEMA3G | NRP2_PLXNA1 | 0.007568468 | 0 | SEMA3G_NRP2_PLXNA1 | SEMA3G - (NRP2+PLXNA1) | SEMA3 | Secreted Signaling | PMID: 27533782 |
| C-NP.MRGPRX1/MRGPRX4 | SEMA3G | NRP2_PLXNA2 | 0.010298053 | 0 | SEMA3G_NRP2_PLXNA2 | SEMA3G - (NRP2+PLXNA2) | SEMA3 | Secreted Signaling | PMID: 27533782 |
| C-NP.SST/CCK | SEMA3G | NRP2_PLXNA2 | 0.012319711 | 0 | SEMA3G_NRP2_PLXNA2 | SEMA3G - (NRP2+PLXNA2) | SEMA3 | Secreted Signaling | PMID: 27533782 |
| C-NP.MRGPRX1/MRGPRX4 | SEMA3G | NRP2_PLXNA3 | 0.013919785 | 0 | SEMA3G_NRP2_PLXNA3 | SEMA3G - (NRP2+PLXNA3) | SEMA3 | Secreted Signaling | PMID: 27533782 |
| C-NP.SST/CCK | SEMA3G | NRP2_PLXNA3 | 0.016640487 | 0 | SEMA3G_NRP2_PLXNA3 | SEMA3G - (NRP2+PLXNA3) | SEMA3 | Secreted Signaling | PMID: 27533782 |
| C-NP.MRGPRX1/MRGPRX4 | SEMA3G | NRP2_PLXNA4 | 0.015833536 | 0 | SEMA3G_NRP2_PLXNA4 | SEMA3G - (NRP2+PLXNA4) | SEMA3 | Secreted Signaling | PMID: 27533782 |
| C-NP.SST/CCK | SEMA3G | NRP2_PLXNA4 | 0.018921114 | 0 | SEMA3G_NRP2_PLXNA4 | SEMA3G - (NRP2+PLXNA4) | SEMA3 | Secreted Signaling | PMID: 27533782 |
| C-NP.MRGPRX1/MRGPRX4 | SEMA3C | NRP1_NRP2 | 0.001292038 | 0 | SEMA3C_NRP1_NRP2 | SEMA3C - (NRP1+NRP2) | SEMA3 | Secreted Signaling | PMID: 27533782 |
| C-PEP.TAC1/CACNG5 | SEMA3C | NRP1_NRP2 | 0.000508295 | 0.02 | SEMA3C_NRP1_NRP2 | SEMA3C - (NRP1+NRP2) | SEMA3 | Secreted Signaling | PMID: 27533782 |
| A-PEP.NTRK3/S100A16 | SEMA3E | PLXND1 | 0.002727554 | 0.03 | SEMA3E_PLXND1 | SEMA3E - PLXND1 | SEMA3 | Secreted Signaling | PMID: 22325954 |
| C-LTMR.CDH9 | SEMA3E | PLXND1 | 0.00704376 | 0.03 | SEMA3E_PLXND1 | SEMA3E - PLXND1 | SEMA3 | Secreted Signaling | PMID: 22325954 |
| C-NP.MRGPRX1/MRGPRX4 | ANXA1 | FPR1 | 0.000566602 | 0 | ANXA1_FPR1 | ANXA1 - FPR1 | ANNEXIN | Secreted Signaling | PMID: 23230437 |
| C-NP.SST/CCK | ANXA1 | FPR1 | 0.000219815 | 0.03 | ANXA1_FPR1 | ANXA1 - FPR1 | ANNEXIN | Secreted Signaling | PMID: 23230437 |
| A-PEP.CHRNA7/SLC18A3 | GAS6 | TYRO3 | 0.007177057 | 0.02 | GAS6_TYRO3 | GAS6 - TYRO3 | GAS | Secreted Signaling | PMID: 27801848 |
| A-PEP.KIT | GAS6 | TYRO3 | 0.009758481 | 0 | GAS6_TYRO3 | GAS6 - TYRO3 | GAS | Secreted Signaling | PMID: 27801848 |
| A-PEP.NTRK3/S100A16 | GAS6 | TYRO3 | 0.011427436 | 0 | GAS6_TYRO3 | GAS6 - TYRO3 | GAS | Secreted Signaling | PMID: 27801848 |
| A-PEP.SCGN/ADRA2C | GAS6 | TYRO3 | 0.010911414 | 0 | GAS6_TYRO3 | GAS6 - TYRO3 | GAS | Secreted Signaling | PMID: 27801848 |

| | | | | | | | | | |
|---|---|---|---|---|---|---|---|---|---|
| C-NP.MRGPRX1/MRGPRX4 | GAS6 | TYRO3 | 0.009655697 | 0 | GAS6_TYRO3 | GAS6 - TYRO3 | GAS | Secreted Signaling | PMID: 27801848 |
| C-NP.SST/CCK | GAS6 | TYRO3 | 0.010523758 | 0 | GAS6_TYRO3 | GAS6 - TYRO3 | GAS | Secreted Signaling | PMID: 27801848 |
| C-PEP.ADORA2B | GAS6 | TYRO3 | 0.007638793 | 0.03 | GAS6_TYRO3 | GAS6 - TYRO3 | GAS | Secreted Signaling | PMID: 27801848 |
| C-PEP.TAC1/CACNG5 | GAS6 | TYRO3 | 0.006934268 | 0.03 | GAS6_TYRO3 | GAS6 - TYRO3 | GAS | Secreted Signaling | PMID: 27801848 |
| C-PEP.TAC1/CHRNA3 | GAS6 | TYRO3 | 0.007831599 | 0.01 | GAS6_TYRO3 | GAS6 - TYRO3 | GAS | Secreted Signaling | PMID: 27801848 |
| C-PEP.ADORA2B | PROS1 | AXL | 5.33506E-05 | 0 | PROS1_AXL | PROS1 - AXL | PROS | Secreted Signaling | PMID: 29531161 |
| C-PEP.TAC1/CACNG5 | PROS1 | AXL | 4.43008E-05 | 0.01 | PROS1_AXL | PROS1 - AXL | PROS | Secreted Signaling | PMID: 29531161 |
| C-NP.SST/CCK | PROS1 | TYRO3 | 0.000183675 | 0 | PROS1_TYRO3 | PROS1 - TYRO3 | PROS | Secreted Signaling | PMID: 30501104 |
| C-PEP.ADORA2B | PROS1 | TYRO3 | 0.001364441 | 0 | PROS1_TYRO3 | PROS1 - TYRO3 | PROS | Secreted Signaling | PMID: 30501104 |
| C-PEP.TAC1/CACNG5 | PROS1 | TYRO3 | 0.001133243 | 0 | PROS1_TYRO3 | PROS1 - TYRO3 | PROS | Secreted Signaling | PMID: 30501104 |
| C-PEP.TAC1/CHRNA3 | PROS1 | TYRO3 | 0.00023986 | 0 | PROS1_TYRO3 | PROS1 - TYRO3 | PROS | Secreted Signaling | PMID: 30501104 |
| C-NP.MRGPRX1/MRGPRX4 | LGALS9 | CD44 | 0.003691675 | 0.02 | LGALS9_CD44 | LGALS9 - CD44 | GALECTIN | Secreted Signaling | PMID: 25065622 |
| C-LTMR.CDH9 | TAFA4 | FPR1 | 6.57609E-05 | 0 | TAFA4_FPR1 | TAFA4 - FPR1 | TAFA | Secreted Signaling | PMID: 35712659;PMID: 35712659 |
| A-PEP.CHRNA7/SLC18A3 | PTPRS | NTRK3 | 0.036340889 | 0.01 | PTPRS_NTRK3 | PTPRS - NTRK3 | PTPR | Secreted Signaling | PMID:21262467;PMID:25385546; |
| A-PEP.KIT | PTPRS | NTRK3 | 0.038235047 | 0 | PTPRS_NTRK3 | PTPRS - NTRK3 | PTPR | Secreted Signaling | PMID:21262467;PMID:25385546; |
| A-PEP.NTRK3/S100A16 | PTPRS | NTRK3 | 0.040222351 | 0 | PTPRS_NTRK3 | PTPRS - NTRK3 | PTPR | Secreted Signaling | PMID:21262467;PMID:25385546; |
| A-PEP.SCGN/ADRA2C | PTPRS | NTRK3 | 0.041174248 | 0 | PTPRS_NTRK3 | PTPRS - NTRK3 | PTPR | Secreted Signaling | PMID:21262467;PMID:25385546; |
| C-LTMR.CDH9 | PTPRS | NTRK3 | 0.039969481 | 0 | PTPRS_NTRK3 | PTPRS - NTRK3 | PTPR | Secreted Signaling | PMID:21262467;PMID:25385546; |
| C-NP.MRGPRX1/MRGPRX4 | PTPRS | NTRK3 | 0.033073687 | 0.01 | PTPRS_NTRK3 | PTPRS - NTRK3 | PTPR | Secreted Signaling | PMID:21262467;PMID:25385546; |
| C-NP.SST/CCK | PTPRS | NTRK3 | 0.056495139 | 0 | PTPRS_NTRK3 | PTPRS - NTRK3 | PTPR | Secreted Signaling | PMID:21262467;PMID:25385546; |
| C-PEP.ADORA2B | PTPRS | NTRK3 | 0.035541188 | 0.01 | PTPRS_NTRK3 | PTPRS - NTRK3 | PTPR | Secreted Signaling | PMID:21262467;PMID:25385546; |
| C-PEP.TAC1/CACNG5 | PTPRS | NTRK3 | 0.042364087 | 0 | PTPRS_NTRK3 | PTPRS - NTRK3 | PTPR | Secreted Signaling | PMID:21262467;PMID:25385546; |

| | | | | | | | | | |
|---|---|---|---|---|---|---|---|---|---|
| C-PEP.TAC1/CHRNA3 | PTPRS | NTRK3 | 0.044569022 | 0 | PTPRS_NTRK3 | PTPRS - NTRK3 | PTPR | Secreted Signaling | PMID:21262467;PMID:25385546; |
| C-NP.MRGPRX1/MRGPRX4 | SLITRK1 | PTPRD | 0.001453186 | 0 | SLITRK1_PTPRD | SLITRK1 - PTPRD | SLITRK | Secreted Signaling | PMID: 30648269;PMID: 30822649 |
| C-LTMR.CDH9 | SLITRK1 | PTPRS | 0.033479726 | 0 | SLITRK1_PTPRS | SLITRK1 - PTPRS | SLITRK | Secreted Signaling | PMID: 30648269;PMID: 30822649 |
| C-NP.MRGPRX1/MRGPRX4 | SLITRK1 | PTPRS | 0.018813321 | 0 | SLITRK1_PTPRS | SLITRK1 - PTPRS | SLITRK | Secreted Signaling | PMID: 30648269;PMID: 30822649 |
| C-PEP.TAC1/CACNG5 | SLITRK2 | PTPRS | 0.012156524 | 0.01 | SLITRK2_PTPRS | SLITRK2 - PTPRS | SLITRK | Secreted Signaling | PMID: 30648269;PMID: 30822649 |
| C-PEP.TAC1/CHRNA3 | SLITRK2 | PTPRS | 0.011684286 | 0.02 | SLITRK2_PTPRS | SLITRK2 - PTPRS | SLITRK | Secreted Signaling | PMID: 30648269;PMID: 30822649 |
| A-PEP.KIT | SLITRK4 | PTPRS | 0.024833343 | 0 | SLITRK4_PTPRS | SLITRK4 - PTPRS | SLITRK | Secreted Signaling | PMID: 30648269;PMID: 30822649 |
| A-PEP.NTRK3/S100A16 | SLITRK4 | PTPRS | 0.011901276 | 0.04 | SLITRK4_PTPRS | SLITRK4 - PTPRS | SLITRK | Secreted Signaling | PMID: 30648269;PMID: 30822649 |
| A-PEP.SCGN/ADRA2C | SLITRK4 | PTPRS | 0.01816898 | 0 | SLITRK4_PTPRS | SLITRK4 - PTPRS | SLITRK | Secreted Signaling | PMID: 30648269;PMID: 30822649 |
| A-PEP.SCGN/ADRA2C | SLITRK5 | PTPRS | 0.015821065 | 0.03 | SLITRK5_PTPRS | SLITRK5 - PTPRS | SLITRK | Secreted Signaling | PMID: 30648269;PMID: 30822649 |
| C-NP.MRGPRX1/MRGPRX4 | SLITRK5 | PTPRS | 0.015582364 | 0 | SLITRK5_PTPRS | SLITRK5 - PTPRS | SLITRK | Secreted Signaling | PMID: 30648269;PMID: 30822649 |
| A-PEP.SCGN/ADRA2C | BDNF | NGFR | 0.01266276 | 0.01 | BDNF_NGFR | BDNF - NGFR | NT | Secreted Signaling | uniprot |
| C-NP.MRGPRX1/MRGPRX4 | BDNF | NGFR | 0.016487409 | 0 | BDNF_NGFR | BDNF - NGFR | NT | Secreted Signaling | uniprot |
| C-NP.MRGPRX1/MRGPRX4 | LGALS9 | P4HB | 0.000773263 | 0.02 | LGALS9_P4HB | LGALS9 - P4HB | GALECTIN | Secreted Signaling | PMID:21670307;uniprot |
| A-PEP.CHRNA7/SLC18A3 | PPIA | BSG | 0.177190527 | 0 | PPIA_BSG | PPIA - BSG | CypA | Secreted Signaling | PMID: 36012604 |
| A-PEP.NTRK3/S100A16 | PPIA | BSG | 0.173610301 | 0.01 | PPIA_BSG | PPIA - BSG | CypA | Secreted Signaling | PMID: 36012604 |
| C-LTMR.CDH9 | SLIT1 | ROBO1 | 0.004907165 | 0.01 | SLIT1_ROBO1 | SLIT1 - ROBO1 | SLIT | Secreted Signaling | PMID:10102268 |
| C-NP.MRGPRX1/MRGPRX4 | SLIT1 | ROBO1 | 0.002650044 | 0 | SLIT1_ROBO1 | SLIT1 - ROBO1 | SLIT | Secreted Signaling | PMID:10102268 |
| C-LTMR.CDH9 | SLIT2 | ROBO1 | 0.001265774 | 0 | SLIT2_ROBO1 | SLIT2 - ROBO1 | SLIT | Secreted Signaling | PMID:10102268 |
| C-NP.SST/CCK | SLIT2 | ROBO1 | 8.56372E-05 | 0.01 | SLIT2_ROBO1 | SLIT2 - ROBO1 | SLIT | Secreted Signaling | PMID:10102268 |
| C-PEP.ADORA2B | SLIT2 | ROBO1 | 0.000130975 | 0.02 | SLIT2_ROBO1 | SLIT2 - ROBO1 | SLIT | Secreted Signaling | PMID:10102268 |

| | | | | | | | | | |
|---|---|---|---|---|---|---|---|---|---|
| C-LTMR.CDH9 | SLIT2 | ROBO2 | 0.01148193 | 0 | SLIT2_ROBO2 | SLIT2 - ROBO2 | SLIT | Secreted Signaling | PMID:10102268 |
| C-NP.SST/CCK | SLIT2 | ROBO2 | 0.000784301 | 0 | SLIT2_ROBO2 | SLIT2 - ROBO2 | SLIT | Secreted Signaling | PMID:10102268 |
| C-PEP.ADORA2B | SLIT2 | ROBO2 | 0.001199083 | 0 | SLIT2_ROBO2 | SLIT2 - ROBO2 | SLIT | Secreted Signaling | PMID:10102268 |
| A-PEP.SCGN/ADRA2C | TUB | MERTK | 5.69996E-05 | 0 | TUB_MERTK | TUB - MERTK | TULP | Secreted Signaling | PMID: 20978475 |
| C-PEP.ADORA2B | TUB | MERTK | 0.000137591 | 0 | TUB_MERTK | TUB - MERTK | TULP | Secreted Signaling | PMID: 20978475 |
| C-PEP.ADORA2B | PROS1 | MERTK | 7.08919E-05 | 0 | PROS1_MERTK | PROS1 - MERTK | PROS | Secreted Signaling | PMID: 34631419 |
| C-PEP.TAC1/CACNG5 | PROS1 | MERTK | 5.88667E-05 | 0 | PROS1_MERTK | PROS1 - MERTK | PROS | Secreted Signaling | PMID: 34631419 |
| C-PEP.TAC1/CHRNA3 | PROS1 | MERTK | 1.24491E-05 | 0.03 | PROS1_MERTK | PROS1 - MERTK | PROS | Secreted Signaling | PMID: 34631419 |
| A-PEP.CHRNA7/SLC18A3 | Ach-CHAT_SLC18A3 | CHRNA7 | 0.000159764 | 0 | Acetylcholine-Ach-CHAT_SLC18A3_CHRNA7 | Ach-(CHAT+SLC18A3) - CHRNA7 | Ach | Non-protein Signaling | PMID: 33674322;uniprot;reactome;IUPHAR |
| A-PEP.SCGN/ADRA2C | ADO-NT5E_SLC29A4 | ADORA1 | 2.40917E-05 | 0 | Adenosine-ADO-NT5E_SLC29A4_ADORA1 | ADO-(NT5E+SLC29A4) - ADORA1 | Adenosine | Non-protein Signaling | PMID:32514148 |
| C-NP.SST/CCK | ADO-NT5E_SLC29A4 | ADORA1 | 4.98279E-06 | 0.04 | Adenosine-ADO-NT5E_SLC29A4_ADORA1 | ADO-(NT5E+SLC29A4) - ADORA1 | Adenosine | Non-protein Signaling | PMID:32514148 |
| C-PEP.ADORA2B | ADO-NT5E_SLC29A4 | ADORA1 | 5.44208E-05 | 0 | Adenosine-ADO-NT5E_SLC29A4_ADORA1 | ADO-(NT5E+SLC29A4) - ADORA1 | Adenosine | Non-protein Signaling | PMID:32514148 |
| C-PEP.TAC1/CACNG5 | ADO-NT5E_SLC29A4 | ADORA1 | 1.57621E-05 | 0 | Adenosine-ADO-NT5E_SLC29A4_ADORA1 | ADO-(NT5E+SLC29A4) - ADORA1 | Adenosine | Non-protein Signaling | PMID:32514148 |
| A-PEP.CHRNA7/SLC18A3 | Ach-CHAT_SLC18A3 | CHRM4 | 0.001631514 | 0 | Acetylcholine-Ach-CHAT_SLC18A3_CHRM4 | Ach-(CHAT+SLC18A3) - CHRM4 | Ach | Non-protein Signaling | PMID: 33674322;PMID: 25176177 |
| A-PEP.KIT | Glu-SLC17A7_GLS | GRIA2 | 0.002989357 | 0 | Glutamate-Glu-SLC17A7_GLS_GRIA2 | Glu-(SLC17A7+GLS) - GRIA2 | Glutamate | Non-protein Signaling | PMID: 24155668;PMID: 14612154 |
| A-PEP.NTRK3/S100A16 | Glu-SLC17A7_GLS | GRIA2 | 0.002855661 | 0 | Glutamate-Glu-SLC17A7_GLS_GRIA2 | Glu-(SLC17A7+GLS) - GRIA2 | Glutamate | Non-protein Signaling | PMID: 24155668;PMID: 14612154 |
| A-PEP.SCGN/ADRA2C | Glu-SLC17A7_GLS | GRIA2 | 0.003269992 | 0 | Glutamate-Glu-SLC17A7_GLS_GRIA2 | Glu-(SLC17A7+GLS) - GRIA2 | Glutamate | Non-protein Signaling | PMID: 24155668;PMID: 14612154 |
| A-PEP.SCGN/ADRA2C | Glu-SLC1A1_GLS | GRIA2 | 0.00038468 | 0 | Glutamate-Glu-SLC1A1_GLS_GRIA2 | Glu-(SLC1A1+GLS) - GRIA2 | Glutamate | Non-protein Signaling | PMID: 24155668;PMID: 14612154 |
| A-PEP.SCGN/ADRA2C | Glu-SLC1A6_GLS | GRIA2 | 0.000739277 | 0 | Glutamate-Glu-SLC1A6_GLS_GRIA2 | Glu-(SLC1A6+GLS) - GRIA2 | Glutamate | Non-protein Signaling | PMID: 24155668;PMID: 14612154 |

| | | | | | | | | | |
|---|---|---|---|---|---|---|---|---|---|
| C-LTMR.CDH9 | Glu-SLC1A6_GLS | GRIA2 | 0.000852594 | 0 | Glutamate-Glu-SLC1A6_GLS_GRIA2 | Glu-(SLC1A6+GLS) - GRIA2 | Glutamate | Non-protein Signaling | PMID: 24155668;PMID: 14612154 |
| A-PEP.KIT | Glu-SLC17A7_GLS | GRIA4 | 0.001603707 | 0 | Glutamate-Glu-SLC17A7_GLS_GRIA4 | Glu-(SLC17A7+GLS) - GRIA4 | Glutamate | Non-protein Signaling | PMID: 24155668;PMID: 14612154 |
| A-PEP.NTRK3/S100A16 | Glu-SLC17A7_GLS | GRIA4 | 0.001531887 | 0 | Glutamate-Glu-SLC17A7_GLS_GRIA4 | Glu-(SLC17A7+GLS) - GRIA4 | Glutamate | Non-protein Signaling | PMID: 24155668;PMID: 14612154 |
| A-PEP.SCGN/ADRA2C | Glu-SLC17A7_GLS | GRIA4 | 0.001754489 | 0 | Glutamate-Glu-SLC17A7_GLS_GRIA4 | Glu-(SLC17A7+GLS) - GRIA4 | Glutamate | Non-protein Signaling | PMID: 24155668;PMID: 14612154 |
| A-PEP.SCGN/ADRA2C | Glu-SLC1A1_GLS | GRIA4 | 0.000206121 | 0 | Glutamate-Glu-SLC1A1_GLS_GRIA4 | Glu-(SLC1A1+GLS) - GRIA4 | Glutamate | Non-protein Signaling | PMID: 24155668;PMID: 14612154 |
| A-PEP.SCGN/ADRA2C | Glu-SLC1A6_GLS | GRIA4 | 0.000396187 | 0 | Glutamate-Glu-SLC1A6_GLS_GRIA4 | Glu-(SLC1A6+GLS) - GRIA4 | Glutamate | Non-protein Signaling | PMID: 24155668;PMID: 14612154 |
| C-LTMR.CDH9 | Glu-SLC1A6_GLS | GRIA4 | 0.000456939 | 0.01 | Glutamate-Glu-SLC1A6_GLS_GRIA4 | Glu-(SLC1A6+GLS) - GRIA4 | Glutamate | Non-protein Signaling | PMID: 24155668;PMID: 14612154 |
| A-PEP.SCGN/ADRA2C | Glu-SLC1A1_GLS | GRIK1 | 0.000171995 | 0 | Glutamate-Glu-SLC1A1_GLS_GRIK1 | Glu-(SLC1A1+GLS) - GRIK1 | Glutamate | Non-protein Signaling | PMID: 24155668;PMID: 14612154 |
| A-PEP.SCGN/ADRA2C | Glu-SLC1A6_GLS | GRIK1 | 0.000330603 | 0.01 | Glutamate-Glu-SLC1A6_GLS_GRIK1 | Glu-(SLC1A6+GLS) - GRIK1 | Glutamate | Non-protein Signaling | PMID: 24155668;PMID: 14612154 |
| A-PEP.NTRK3/S100A16 | Glu-SLC17A7_GLS | GRIK2 | 0.004742181 | 0.02 | Glutamate-Glu-SLC17A7_GLS_GRIK2 | Glu-(SLC17A7+GLS) - GRIK2 | Glutamate | Non-protein Signaling | PMID: 24155668;PMID: 14612154 |
| A-PEP.SCGN/ADRA2C | Glu-SLC17A7_GLS | GRIK2 | 0.00542874 | 0.03 | Glutamate-Glu-SLC17A7_GLS_GRIK2 | Glu-(SLC17A7+GLS) - GRIK2 | Glutamate | Non-protein Signaling | PMID: 24155668;PMID: 14612154 |
| A-PEP.SCGN/ADRA2C | Glu-SLC1A1_GLS | GRIK2 | 0.000639857 | 0 | Glutamate-Glu-SLC1A1_GLS_GRIK2 | Glu-(SLC1A1+GLS) - GRIK2 | Glutamate | Non-protein Signaling | PMID: 24155668;PMID: 14612154 |
| A-PEP.SCGN/ADRA2C | Glu-SLC1A6_GLS | GRIK2 | 0.001229386 | 0.01 | Glutamate-Glu-SLC1A6_GLS_GRIK2 | Glu-(SLC1A6+GLS) - GRIK2 | Glutamate | Non-protein Signaling | PMID: 24155668;PMID: 14612154 |
| A-PEP.KIT | Glu-SLC17A7_GLS | GRIK3 | 0.00632632 | 0 | Glutamate-Glu-SLC17A7_GLS_GRIK3 | Glu-(SLC17A7+GLS) - GRIK3 | Glutamate | Non-protein Signaling | PMID: 24155668;PMID: 14612154 |
| A-PEP.NTRK3/S100A16 | Glu-SLC17A7_GLS | GRIK3 | 0.006044286 | 0 | Glutamate-Glu-SLC17A7_GLS_GRIK3 | Glu-(SLC17A7+GLS) - GRIK3 | Glutamate | Non-protein Signaling | PMID: 24155668;PMID: 14612154 |
| A-PEP.SCGN/ADRA2C | Glu-SLC17A7_GLS | GRIK3 | 0.006918049 | 0 | Glutamate-Glu-SLC17A7_GLS_GRIK3 | Glu-(SLC17A7+GLS) - GRIK3 | Glutamate | Non-protein Signaling | PMID: 24155668;PMID: 14612154 |

| | | | | | | | | | |
|---|---|---|---|---|---|---|---|---|---|
| A-PEP.SCGN/ADRA2C | Glu-SLC1A1_GLS | GRIK3 | 0.000816473 | 0 | Glutamate-Glu-SLC1A1_GLS_GRIK3 | Glu-(SLC1A1+GLS) - GRIK3 | Glutamate | Non-protein Signaling | PMID: 24155668;PMID: 14612154 |
| C-PEP.ADORA2B | Glu-SLC1A2_GLS | GRIK3 | 0.001511975 | 0.03 | Glutamate-Glu-SLC1A2_GLS_GRIK3 | Glu-(SLC1A2+GLS) - GRIK3 | Glutamate | Non-protein Signaling | PMID: 24155668;PMID: 14612154 |
| C-PEP.TAC1/CHRNA3 | Glu-SLC1A2_GLS | GRIK3 | 0.001568007 | 0.04 | Glutamate-Glu-SLC1A2_GLS_GRIK3 | Glu-(SLC1A2+GLS) - GRIK3 | Glutamate | Non-protein Signaling | PMID: 24155668;PMID: 14612154 |
| A-PEP.SCGN/ADRA2C | Glu-SLC1A6_GLS | GRIK3 | 0.00156847 | 0 | Glutamate-Glu-SLC1A6_GLS_GRIK3 | Glu-(SLC1A6+GLS) - GRIK3 | Glutamate | Non-protein Signaling | PMID: 24155668;PMID: 14612154 |
| C-LTMR.CDH9 | Glu-SLC1A6_GLS | GRIK3 | 0.001808657 | 0 | Glutamate-Glu-SLC1A6_GLS_GRIK3 | Glu-(SLC1A6+GLS) - GRIK3 | Glutamate | Non-protein Signaling | PMID: 24155668;PMID: 14612154 |
| A-PEP.CHRNA7/SLC18A3 | Glu-SLC17A6_GLS | GRM4 | 0.024839176 | 0 | Glutamate-Glu-SLC17A6_GLS_GRM4 | Glu-(SLC17A6+GLS) - GRM4 | Glutamate | Non-protein Signaling | PMID: 24155668;PMID: 14612154 |
| A-PEP.KIT | Glu-SLC17A6_GLS | GRM4 | 0.026260273 | 0 | Glutamate-Glu-SLC17A6_GLS_GRM4 | Glu-(SLC17A6+GLS) - GRM4 | Glutamate | Non-protein Signaling | PMID: 24155668;PMID: 14612154 |
| A-PEP.NTRK3/S100A16 | Glu-SLC17A6_GLS | GRM4 | 0.024793991 | 0 | Glutamate-Glu-SLC17A6_GLS_GRM4 | Glu-(SLC17A6+GLS) - GRM4 | Glutamate | Non-protein Signaling | PMID: 24155668;PMID: 14612154 |
| A-PEP.SCGN/ADRA2C | Glu-SLC17A6_GLS | GRM4 | 0.023960497 | 0 | Glutamate-Glu-SLC17A6_GLS_GRM4 | Glu-(SLC17A6+GLS) - GRM4 | Glutamate | Non-protein Signaling | PMID: 24155668;PMID: 14612154 |
| C-LTMR.CDH9 | Glu-SLC17A6_GLS | GRM4 | 0.027254157 | 0 | Glutamate-Glu-SLC17A6_GLS_GRM4 | Glu-(SLC17A6+GLS) - GRM4 | Glutamate | Non-protein Signaling | PMID: 24155668;PMID: 14612154 |
| C-NP.MRGPRX1/MRGPRX4 | Glu-SLC17A6_GLS | GRM4 | 0.018755236 | 0 | Glutamate-Glu-SLC17A6_GLS_GRM4 | Glu-(SLC17A6+GLS) - GRM4 | Glutamate | Non-protein Signaling | PMID: 24155668;PMID: 14612154 |
| C-NP.SST/CCK | Glu-SLC17A6_GLS | GRM4 | 0.030419736 | 0 | Glutamate-Glu-SLC17A6_GLS_GRM4 | Glu-(SLC17A6+GLS) - GRM4 | Glutamate | Non-protein Signaling | PMID: 24155668;PMID: 14612154 |
| C-PEP.ADORA2B | Glu-SLC17A6_GLS | GRM4 | 0.029329997 | 0 | Glutamate-Glu-SLC17A6_GLS_GRM4 | Glu-(SLC17A6+GLS) - GRM4 | Glutamate | Non-protein Signaling | PMID: 24155668;PMID: 14612154 |
| C-PEP.TAC1/CACNG5 | Glu-SLC17A6_GLS | GRM4 | 0.027588409 | 0 | Glutamate-Glu-SLC17A6_GLS_GRM4 | Glu-(SLC17A6+GLS) - GRM4 | Glutamate | Non-protein Signaling | PMID: 24155668;PMID: 14612154 |
| C-PEP.TAC1/CHRNA3 | Glu-SLC17A6_GLS | GRM4 | 0.032170417 | 0 | Glutamate-Glu-SLC17A6_GLS_GRM4 | Glu-(SLC17A6+GLS) - GRM4 | Glutamate | Non-protein Signaling | PMID: 24155668;PMID: 14612154 |
| A-PEP.CHRNA7/SLC18A3 | Glu-SLC17A7_GLS | GRM4 | 0.007719483 | 0.01 | Glutamate-Glu-SLC17A7_GLS_GRM4 | Glu-(SLC17A7+GLS) - GRM4 | Glutamate | Non-protein Signaling | PMID: 24155668;PMID: 14612154 |

| | | | | | | | | | |
|---|---|---|---|---|---|---|---|---|---|
| A-PEP.KIT | Glu-SLC17A7_GLS | GRM4 | 0.031107919 | 0 | Glutamate-Glu-SLC17A7_GLS_GRM4 | Glu-(SLC17A7+GLS) - GRM4 | Glutamate | Non-protein Signaling | PMID: 24155668;PMID: 14612154 |
| A-PEP.NTRK3/S100A16 | Glu-SLC17A7_GLS | GRM4 | 0.029754174 | 0 | Glutamate-Glu-SLC17A7_GLS_GRM4 | Glu-(SLC17A7+GLS) - GRM4 | Glutamate | Non-protein Signaling | PMID: 24155668;PMID: 14612154 |
| A-PEP.SCGN/ADRA2C | Glu-SLC17A7_GLS | GRM4 | 0.033938412 | 0 | Glutamate-Glu-SLC17A7_GLS_GRM4 | Glu-(SLC17A7+GLS) - GRM4 | Glutamate | Non-protein Signaling | PMID: 24155668;PMID: 14612154 |
| C-PEP.TAC1/CACNG5 | Glu-SLC17A7_GLS | GRM4 | 0.007721119 | 0.02 | Glutamate-Glu-SLC17A7_GLS_GRM4 | Glu-(SLC17A7+GLS) - GRM4 | Glutamate | Non-protein Signaling | PMID: 24155668;PMID: 14612154 |
| C-PEP.TAC1/CHRNA3 | Glu-SLC17A7_GLS | GRM4 | 0.007172197 | 0 | Glutamate-Glu-SLC17A7_GLS_GRM4 | Glu-(SLC17A7+GLS) - GRM4 | Glutamate | Non-protein Signaling | PMID: 24155668;PMID: 14612154 |
| A-PEP.SCGN/ADRA2C | Glu-SLC1A1_GLS | GRM4 | 0.004103918 | 0 | Glutamate-Glu-SLC1A1_GLS_GRM4 | Glu-(SLC1A1+GLS) - GRM4 | Glutamate | Non-protein Signaling | PMID: 24155668;PMID: 14612154 |
| A-PEP.CHRNA7/SLC18A3 | Glu-SLC1A2_GLS | GRM4 | 0.003969858 | 0.01 | Glutamate-Glu-SLC1A2_GLS_GRM4 | Glu-(SLC1A2+GLS) - GRM4 | Glutamate | Non-protein Signaling | PMID: 24155668;PMID: 14612154 |
| A-PEP.KIT | Glu-SLC1A2_GLS | GRM4 | 0.007155456 | 0 | Glutamate-Glu-SLC1A2_GLS_GRM4 | Glu-(SLC1A2+GLS) - GRM4 | Glutamate | Non-protein Signaling | PMID: 24155668;PMID: 14612154 |
| A-PEP.SCGN/ADRA2C | Glu-SLC1A2_GLS | GRM4 | 0.006257569 | 0 | Glutamate-Glu-SLC1A2_GLS_GRM4 | Glu-(SLC1A2+GLS) - GRM4 | Glutamate | Non-protein Signaling | PMID: 24155668;PMID: 14612154 |
| C-LTMR.CDH9 | Glu-SLC1A2_GLS | GRM4 | 0.004916828 | 0.01 | Glutamate-Glu-SLC1A2_GLS_GRM4 | Glu-(SLC1A2+GLS) - GRM4 | Glutamate | Non-protein Signaling | PMID: 24155668;PMID: 14612154 |
| C-NP.SST/CCK | Glu-SLC1A2_GLS | GRM4 | 0.007117207 | 0 | Glutamate-Glu-SLC1A2_GLS_GRM4 | Glu-(SLC1A2+GLS) - GRM4 | Glutamate | Non-protein Signaling | PMID: 24155668;PMID: 14612154 |
| C-PEP.ADORA2B | Glu-SLC1A2_GLS | GRM4 | 0.007578549 | 0 | Glutamate-Glu-SLC1A2_GLS_GRM4 | Glu-(SLC1A2+GLS) - GRM4 | Glutamate | Non-protein Signaling | PMID: 24155668;PMID: 14612154 |
| C-PEP.TAC1/CACNG5 | Glu-SLC1A2_GLS | GRM4 | 0.007983627 | 0 | Glutamate-Glu-SLC1A2_GLS_GRM4 | Glu-(SLC1A2+GLS) - GRM4 | Glutamate | Non-protein Signaling | PMID: 24155668;PMID: 14612154 |
| C-PEP.TAC1/CHRNA3 | Glu-SLC1A2_GLS | GRM4 | 0.007857631 | 0 | Glutamate-Glu-SLC1A2_GLS_GRM4 | Glu-(SLC1A2+GLS) - GRM4 | Glutamate | Non-protein Signaling | PMID: 24155668;PMID: 14612154 |
| A-PEP.CHRNA7/SLC18A3 | Glu-SLC1A3_GLS | GRM4 | 0.005144116 | 0 | Glutamate-Glu-SLC1A3_GLS_GRM4 | Glu-(SLC1A3+GLS) - GRM4 | Glutamate | Non-protein Signaling | PMID: 24155668;PMID: 14612154 |
| A-PEP.KIT | Glu-SLC1A3_GLS | GRM4 | 0.006162722 | 0 | Glutamate-Glu-SLC1A3_GLS_GRM4 | Glu-(SLC1A3+GLS) - GRM4 | Glutamate | Non-protein Signaling | PMID: 24155668;PMID: 14612154 |

| | | | | | | | | | |
|---|---|---|---|---|---|---|---|---|---|
| A-PEP.SCGN/ADRA2C | Glu-SLC1A3_GLS | GRM4 | 0.005166457 | 0.01 | Glutamate-Glu-SLC1A3_GLS_GRM4 | Glu-(SLC1A3+GLS) - GRM4 | Glutamate | Non-protein Signaling | PMID: 24155668;PMID: 14612154 |
| C-LTMR.CDH9 | Glu-SLC1A3_GLS | GRM4 | 0.007718465 | 0 | Glutamate-Glu-SLC1A3_GLS_GRM4 | Glu-(SLC1A3+GLS) - GRM4 | Glutamate | Non-protein Signaling | PMID: 24155668;PMID: 14612154 |
| C-NP.MRGPRX1/MRGPRX4 | Glu-SLC1A3_GLS | GRM4 | 0.005996131 | 0 | Glutamate-Glu-SLC1A3_GLS_GRM4 | Glu-(SLC1A3+GLS) - GRM4 | Glutamate | Non-protein Signaling | PMID: 24155668;PMID: 14612154 |
| C-PEP.TAC1/CACNG5 | Glu-SLC1A3_GLS | GRM4 | 0.00751118 | 0 | Glutamate-Glu-SLC1A3_GLS_GRM4 | Glu-(SLC1A3+GLS) - GRM4 | Glutamate | Non-protein Signaling | PMID: 24155668;PMID: 14612154 |
| C-PEP.TAC1/CHRNA3 | Glu-SLC1A3_GLS | GRM4 | 0.00682513 | 0 | Glutamate-Glu-SLC1A3_GLS_GRM4 | Glu-(SLC1A3+GLS) - GRM4 | Glutamate | Non-protein Signaling | PMID: 24155668;PMID: 14612154 |
| A-PEP.SCGN/ADRA2C | Glu-SLC1A6_GLS | GRM4 | 0.007859938 | 0 | Glutamate-Glu-SLC1A6_GLS_GRM4 | Glu-(SLC1A6+GLS) - GRM4 | Glutamate | Non-protein Signaling | PMID: 24155668;PMID: 14612154 |
| C-LTMR.CDH9 | Glu-SLC1A6_GLS | GRM4 | 0.009054825 | 0 | Glutamate-Glu-SLC1A6_GLS_GRM4 | Glu-(SLC1A6+GLS) - GRM4 | Glutamate | Non-protein Signaling | PMID: 24155668;PMID: 14612154 |
| A-PEP.SCGN/ADRA2C | Glu-SLC1A1_GLS | GRM7 | 0.000417626 | 0 | Glutamate-Glu-SLC1A1_GLS_GRM7 | Glu-(SLC1A1+GLS) - GRM7 | Glutamate | Non-protein Signaling | PMID: 24155668;PMID: 14612154 |
| A-PEP.SCGN/ADRA2C | Glu-SLC1A6_GLS | GRM7 | 0.000802567 | 0.01 | Glutamate-Glu-SLC1A6_GLS_GRM7 | Glu-(SLC1A6+GLS) - GRM7 | Glutamate | Non-protein Signaling | PMID: 24155668;PMID: 14612154 |
| C-LTMR.CDH9 | 5-HT-TPH2_SLC18A2 | HTR1D | 2.25629E-05 | 0 | 5-HT-TPH2_SLC18A2_HTR1D | 5-HT-(TPH2+SLC18A2) - HTR1D | 5-HT | Non-protein Signaling | PMID: 23328530 |
| C-PEP.ADORA2B | 5-HT-TPH2_SLC18A2 | HTR1D | 2.59266E-05 | 0 | 5-HT-TPH2_SLC18A2_HTR1D | 5-HT-(TPH2+SLC18A2) - HTR1D | 5-HT | Non-protein Signaling | PMID: 23328530 |
| C-LTMR.CDH9 | 5-HT-TPH2_SLC18A2 | HTR2A | 3.72568E-05 | 0 | 5-HT-TPH2_SLC18A2_HTR2A | 5-HT-(TPH2+SLC18A2) - HTR2A | 5-HT | Non-protein Signaling | PMID: 23328530 |
| C-PEP.ADORA2B | 5-HT-TPH2_SLC18A2 | HTR2A | 4.2811E-05 | 0 | 5-HT-TPH2_SLC18A2_HTR2A | 5-HT-(TPH2+SLC18A2) - HTR2A | 5-HT | Non-protein Signaling | PMID: 23328530 |
| C-LTMR.CDH9 | 5-HT-TPH2_SLC18A2 | HTR5A | 2.6377E-05 | 0 | 5-HT-TPH2_SLC18A2_HTR5A | 5-HT-(TPH2+SLC18A2) - HTR5A | 5-HT | Non-protein Signaling | PMID: 23328530 |
| C-PEP.ADORA2B | 5-HT-TPH2_SLC18A2 | HTR5A | 3.03093E-05 | 0 | 5-HT-TPH2_SLC18A2_HTR5A | 5-HT-(TPH2+SLC18A2) - HTR5A | 5-HT | Non-protein Signaling | PMID: 23328530 |
| C-LTMR.CDH9 | 5-HT-TPH2_SLC18A2 | HTR6 | 1.93052E-05 | 0 | 5-HT-TPH2_SLC18A2_HTR6 | 5-HT-(TPH2+SLC18A2) - HTR6 | 5-HT | Non-protein Signaling | PMID: 23328530 |

| | | | | | | | | | |
|---|---|---|---|---|---|---|---|---|---|
| C-PEP.ADORA2B | 5-HT-TPH2_SLC18A2 | HTR6 | 2.21833E-05 | 0 | 5-HT-TPH2_SLC18A2_HTR6 | 5-HT-(TPH2+SLC18A2) - HTR6 | 5-HT | Non-protein Signaling | PMID: 23328530 |
| C-LTMR.CDH9 | 5-HT-TPH2_SLC18A2 | HTR7 | 6.35178E-05 | 0 | 5-HT-TPH2_SLC18A2_HTR7 | 5-HT-(TPH2+SLC18A2) - HTR7 | 5-HT | Non-protein Signaling | PMID: 23328530 |
| C-PEP.ADORA2B | 5-HT-TPH2_SLC18A2 | HTR7 | 7.29868E-05 | 0 | 5-HT-TPH2_SLC18A2_HTR7 | 5-HT-(TPH2+SLC18A2) - HTR7 | 5-HT | Non-protein Signaling | PMID: 23328530 |
| A-PEP.SCGN/ADRA2C | b-Endorphin-POMC | OPRM1 | 0.000255108 | 0 | b-Endorphin-POMC_OPRM1 | b-ENDORPHIN-POMC - OPRM1 | OPIOID | Non-protein Signaling | PMID: 29934561;PMID: 34724150 |
| A-PEP.SCGN/ADRA2C | b-Endorphin-POMC | OPRK1 | 4.66117E-07 | 0 | b-Endorphin-POMC_OPRK1 | b-ENDORPHIN-POMC - OPRK1 | OPIOID | Non-protein Signaling | PMID: 29934561;PMID: 34724150 |
| A-PEP.KIT | PGE2-PTGES | PTGER3 | 0.007078072 | 0 | PGE2-PTGES_PTGER3 | PGE2-PTGES - PTGER3 | Prostaglandin | Non-protein Signaling | PMID: 34949672;PMID: 21508345 |
| A-PEP.NTRK3/S100A16 | PGE2-PTGES | PTGER3 | 0.006144327 | 0 | PGE2-PTGES_PTGER3 | PGE2-PTGES - PTGER3 | Prostaglandin | Non-protein Signaling | PMID: 34949672;PMID: 21508345 |
| A-PEP.SCGN/ADRA2C | PGE2-PTGES | PTGER3 | 0.008383541 | 0 | PGE2-PTGES_PTGER3 | PGE2-PTGES - PTGER3 | Prostaglandin | Non-protein Signaling | PMID: 34949672;PMID: 21508345 |
| C-LTMR.CDH9 | PGE2-PTGES | PTGER3 | 0.009683408 | 0 | PGE2-PTGES_PTGER3 | PGE2-PTGES - PTGER3 | Prostaglandin | Non-protein Signaling | PMID: 34949672;PMID: 21508345 |
| A-PEP.KIT | PGE2-PTGES2 | PTGER3 | 0.005623337 | 0.04 | PGE2-PTGES2_PTGER3 | PGE2-PTGES2 - PTGER3 | Prostaglandin | Non-protein Signaling | PMID: 34949672;PMID: 21508345 |
| A-PEP.SCGN/ADRA2C | PGE2-PTGES2 | PTGER3 | 0.007813927 | 0 | PGE2-PTGES2_PTGER3 | PGE2-PTGES2 - PTGER3 | Prostaglandin | Non-protein Signaling | PMID: 34949672;PMID: 21508345 |
| C-PEP.ADORA2B | PGE2-PTGES2 | PTGER3 | 0.008084084 | 0 | PGE2-PTGES2_PTGER3 | PGE2-PTGES2 - PTGER3 | Prostaglandin | Non-protein Signaling | PMID: 34949672;PMID: 21508345 |
| A-PEP.CHRNA7/SLC18A3 | PGE2-PTGES3 | PTGER3 | 0.014178829 | 0.03 | PGE2-PTGES3_PTGER3 | PGE2-PTGES3 - PTGER3 | Prostaglandin | Non-protein Signaling | PMID: 34949672;PMID: 21508345 |
| A-PEP.KIT | PGE2-PTGES3 | PTGER3 | 0.013975229 | 0.04 | PGE2-PTGES3_PTGER3 | PGE2-PTGES3 - PTGER3 | Prostaglandin | Non-protein Signaling | PMID: 34949672;PMID: 21508345 |
| A-PEP.NTRK3/S100A16 | PGE2-PTGES3 | PTGER3 | 0.014696078 | 0.02 | PGE2-PTGES3_PTGER3 | PGE2-PTGES3 - PTGER3 | Prostaglandin | Non-protein Signaling | PMID: 34949672;PMID: 21508345 |
| A-PEP.SCGN/ADRA2C | PGE2-PTGES3 | PTGER3 | 0.014473642 | 0.03 | PGE2-PTGES3_PTGER3 | PGE2-PTGES3 - PTGER3 | Prostaglandin | Non-protein Signaling | PMID: 34949672;PMID: 21508345 |

| | | | | | | | | | |
|---|---|---|---|---|---|---|---|---|---|
| C-NP.SST/CCK | PGE2-PTGES3 | PTGER3 | 0.014429338 | 0.04 | PGE2-PTGES3_PTGER3 | PGE2-PTGES3 - PTGER3 | Prostaglandin | Non-protein Signaling | PMID: 34949672;PMID: 21508345 |
| A-PEP.CHRNA7/SLC18A3 | PGE2-PTGES | PTGER4 | 0.002665922 | 0.03 | PGE2-PTGES_PTGER4 | PGE2-PTGES - PTGER4 | Prostaglandin | Non-protein Signaling | PMID: 34949672;PMID: 21508345 |
| A-PEP.KIT | PGE2-PTGES | PTGER4 | 0.005637883 | 0 | PGE2-PTGES_PTGER4 | PGE2-PTGES - PTGER4 | Prostaglandin | Non-protein Signaling | PMID: 34949672;PMID: 21508345 |
| A-PEP.NTRK3/S100A16 | PGE2-PTGES | PTGER4 | 0.004893193 | 0 | PGE2-PTGES_PTGER4 | PGE2-PTGES - PTGER4 | Prostaglandin | Non-protein Signaling | PMID: 34949672;PMID: 21508345 |
| A-PEP.SCGN/ADRA2C | PGE2-PTGES | PTGER4 | 0.006679512 | 0 | PGE2-PTGES_PTGER4 | PGE2-PTGES - PTGER4 | Prostaglandin | Non-protein Signaling | PMID: 34949672;PMID: 21508345 |
| C-LTMR.CDH9 | PGE2-PTGES | PTGER4 | 0.007717226 | 0 | PGE2-PTGES_PTGER4 | PGE2-PTGES - PTGER4 | Prostaglandin | Non-protein Signaling | PMID: 34949672;PMID: 21508345 |
| C-NP.MRGPRX1/MRGPRX4 | PGE2-PTGES | PTGER4 | 0.002716444 | 0.04 | PGE2-PTGES_PTGER4 | PGE2-PTGES - PTGER4 | Prostaglandin | Non-protein Signaling | PMID: 34949672;PMID: 21508345 |
| A-PEP.CHRNA7/SLC18A3 | PGE2-PTGES2 | PTGER4 | 0.00404839 | 0.04 | PGE2-PTGES2_PTGER4 | PGE2-PTGES2 - PTGER4 | Prostaglandin | Non-protein Signaling | PMID: 34949672;PMID: 21508345 |
| A-PEP.KIT | PGE2-PTGES2 | PTGER4 | 0.004477811 | 0.03 | PGE2-PTGES2_PTGER4 | PGE2-PTGES2 - PTGER4 | Prostaglandin | Non-protein Signaling | PMID: 34949672;PMID: 21508345 |
| A-PEP.NTRK3/S100A16 | PGE2-PTGES2 | PTGER4 | 0.004056993 | 0.02 | PGE2-PTGES2_PTGER4 | PGE2-PTGES2 - PTGER4 | Prostaglandin | Non-protein Signaling | PMID: 34949672;PMID: 21508345 |
| A-PEP.SCGN/ADRA2C | PGE2-PTGES2 | PTGER4 | 0.00622495 | 0 | PGE2-PTGES2_PTGER4 | PGE2-PTGES2 - PTGER4 | Prostaglandin | Non-protein Signaling | PMID: 34949672;PMID: 21508345 |
| C-NP.SST/CCK | PGE2-PTGES2 | PTGER4 | 0.004347386 | 0.01 | PGE2-PTGES2_PTGER4 | PGE2-PTGES2 - PTGER4 | Prostaglandin | Non-protein Signaling | PMID: 34949672;PMID: 21508345 |
| C-PEP.ADORA2B | PGE2-PTGES2 | PTGER4 | 0.006440528 | 0 | PGE2-PTGES2_PTGER4 | PGE2-PTGES2 - PTGER4 | Prostaglandin | Non-protein Signaling | PMID: 34949672;PMID: 21508345 |
| C-PEP.TAC1/CACNG5 | PGE2-PTGES2 | PTGER4 | 0.004597699 | 0.01 | PGE2-PTGES2_PTGER4 | PGE2-PTGES2 - PTGER4 | Prostaglandin | Non-protein Signaling | PMID: 34949672;PMID: 21508345 |
| C-PEP.TAC1/CHRNA3 | PGE2-PTGES2 | PTGER4 | 0.004690104 | 0.02 | PGE2-PTGES2_PTGER4 | PGE2-PTGES2 - PTGER4 | Prostaglandin | Non-protein Signaling | PMID: 34949672;PMID: 21508345 |
| A-PEP.CHRNA7/SLC18A3 | PGE2-PTGES3 | PTGER4 | 0.011310293 | 0.01 | PGE2-PTGES3_PTGER4 | PGE2-PTGES3 - PTGER4 | Prostaglandin | Non-protein Signaling | PMID: 34949672;PMID: 21508345 |

| | | | | | | | | | |
|---|---|---|---|---|---|---|---|---|---|
| A-PEP.KIT | PGE2-PTGES3 | PTGER4 | 0.011147418 | 0 | PGE2-PTGES3_PTGER4 | PGE2-PTGES3 - PTGER4 | Prostaglandin | Non-protein Signaling | PMID: 34949672;PMID: 21508345 |
| A-PEP.NTRK3/S100A16 | PGE2-PTGES3 | PTGER4 | 0.011724141 | 0 | PGE2-PTGES3_PTGER4 | PGE2-PTGES3 - PTGER4 | Prostaglandin | Non-protein Signaling | PMID: 34949672;PMID: 21508345 |
| A-PEP.SCGN/ADRA2C | PGE2-PTGES3 | PTGER4 | 0.01154616 | 0.01 | PGE2-PTGES3_PTGER4 | PGE2-PTGES3 - PTGER4 | Prostaglandin | Non-protein Signaling | PMID: 34949672;PMID: 21508345 |
| C-NP.MRGPRX1/MRGPRX4 | PGE2-PTGES3 | PTGER4 | 0.010161021 | 0.04 | PGE2-PTGES3_PTGER4 | PGE2-PTGES3 - PTGER4 | Prostaglandin | Non-protein Signaling | PMID: 34949672;PMID: 21508345 |
| C-NP.SST/CCK | PGE2-PTGES3 | PTGER4 | 0.011510713 | 0.01 | PGE2-PTGES3_PTGER4 | PGE2-PTGES3 - PTGER4 | Prostaglandin | Non-protein Signaling | PMID: 34949672;PMID: 21508345 |
| C-PEP.TAC1/CHRNA3 | PGE2-PTGES3 | PTGER4 | 0.010421012 | 0.02 | PGE2-PTGES3_PTGER4 | PGE2-PTGES3 - PTGER4 | Prostaglandin | Non-protein Signaling | PMID: 34949672;PMID: 21508345 |
| A-PEP.KIT | Glu-SLC17A7_GLS | GRIK2_GRIK5 | 0.005745458 | 0.03 | Glutamate-Glu-SLC17A7_GLS_GRIK2_GRIK5 | Glu-(SLC17A7+GLS) - (GRIK2+GRIK5) | Glutamate | Non-protein Signaling | PMID: 24155668;PMID: 31776169;PMID: 32594024 |
| A-PEP.NTRK3/S100A16 | Glu-SLC17A7_GLS | GRIK2_GRIK5 | 0.005489176 | 0 | Glutamate-Glu-SLC17A7_GLS_GRIK2_GRIK5 | Glu-(SLC17A7+GLS) - (GRIK2+GRIK5) | Glutamate | Non-protein Signaling | PMID: 24155668;PMID: 31776169;PMID: 32594024 |
| A-PEP.SCGN/ADRA2C | Glu-SLC17A7_GLS | GRIK2_GRIK5 | 0.0062832 | 0 | Glutamate-Glu-SLC17A7_GLS_GRIK2_GRIK5 | Glu-(SLC17A7+GLS) - (GRIK2+GRIK5) | Glutamate | Non-protein Signaling | PMID: 24155668;PMID: 31776169;PMID: 32594024 |
| A-PEP.KIT | Glu-SLC17A7_GLS | GRIK3_GRIK5 | 0.006485879 | 0 | Glutamate-Glu-SLC17A7_GLS_GRIK3_GRIK5 | Glu-(SLC17A7+GLS) - (GRIK3+GRIK5) | Glutamate | Non-protein Signaling | PMID: 24155668;PMID: 31776169;PMID: 32594024 |
| A-PEP.NTRK3/S100A16 | Glu-SLC17A7_GLS | GRIK3_GRIK5 | 0.006196776 | 0 | Glutamate-Glu-SLC17A7_GLS_GRIK3_GRIK5 | Glu-(SLC17A7+GLS) - (GRIK3+GRIK5) | Glutamate | Non-protein Signaling | PMID: 24155668;PMID: 31776169;PMID: 32594024 |
| A-PEP.SCGN/ADRA2C | Glu-SLC17A7_GLS | GRIK3_GRIK5 | 0.007092426 | 0 | Glutamate-Glu-SLC17A7_GLS_GRIK3_GRIK5 | Glu-(SLC17A7+GLS) - (GRIK3+GRIK5) | Glutamate | Non-protein Signaling | PMID: 24155668;PMID: 31776169;PMID: 32594024 |
| A-PEP.SCGN/ADRA2C | Glu-SLC1A1_GLS | GRIK1_GRIK5 | 0.000384294 | 0 | Glutamate-Glu-SLC1A1_GLS_GRIK1_GRIK5 | Glu-(SLC1A1+GLS) - (GRIK1+GRIK5) | Glutamate | Non-protein Signaling | PMID: 24155668;PMID: 31776169;PMID: 32594024 |
| A-PEP.SCGN/ADRA2C | Glu-SLC1A1_GLS | GRIK2_GRIK5 | 0.00074113 | 0 | Glutamate-Glu-SLC1A1_GLS_GRIK2_GRIK5 | Glu-(SLC1A1+GLS) - (GRIK2+GRIK5) | Glutamate | Non-protein Signaling | PMID: 24155668;PMID: 31776169;PMID: 32594024 |
| A-PEP.SCGN/ADRA2C | Glu-SLC1A1_GLS | GRIK3_GRIK5 | 0.000837183 | 0 | Glutamate-Glu-SLC1A1_GLS_GRIK3_GRIK5 | Glu-(SLC1A1+GLS) - (GRIK3+GRIK5) | Glutamate | Non-protein Signaling | PMID: 24155668;PMID: 31776169;PMID: 32594024 |
| A-PEP.SCGN/ADRA2C | Glu-SLC1A6_GLS | GRIK1_GRIK5 | 0.000738535 | 0.01 | Glutamate-Glu-SLC1A6_GLS_GRIK1_GRIK5 | Glu-(SLC1A6+GLS) - (GRIK1+GRIK5) | Glutamate | Non-protein Signaling | PMID: 24155668;PMID: 31776169;PMID: 32594024 |

| | | | | | | | | | |
|---|---|---|---|---|---|---|---|---|---|
| A-PEP.SCGN/ADRA2C | Glu-SLC1A6_GLS | GRIK2_GRIK5 | 0.001423832 | 0.01 | Glutamate-Glu-SLC1A6_GLS_GRIK2_GRIK5 | Glu-(SLC1A6+GLS) - (GRIK2+GRIK5) | Glutamate | Non-protein Signaling | PMID: 24155668;PMID: 31776169;PMID: 32594024 |
| A-PEP.SCGN/ADRA2C | Glu-SLC1A6_GLS | GRIK3_GRIK5 | 0.001608223 | 0 | Glutamate-Glu-SLC1A6_GLS_GRIK3_GRIK5 | Glu-(SLC1A6+GLS) - (GRIK3+GRIK5) | Glutamate | Non-protein Signaling | PMID: 24155668;PMID: 31776169;PMID: 32594024 |
| C-LTMR.CDH9 | Glu-SLC1A6_GLS | GRIK3_GRIK5 | 0.001854486 | 0.01 | Glutamate-Glu-SLC1A6_GLS_GRIK3_GRIK5 | Glu-(SLC1A6+GLS) - (GRIK3+GRIK5) | Glutamate | Non-protein Signaling | PMID: 24155668;PMID: 31776169;PMID: 32594024 |
| A-PEP.SCGN/ADRA2C | DHEAS-SULT2B | PPARA | 6.62358E-05 | 0.03 | DHEAsulfate-DHEAS-SULT2B_PPARA | DHEAS-SULT2B - PPARA | DHEAS | Non-protein Signaling | HMRbase |
| A-PEP.NTRK3/S100A16 | DHT-SRD5A1 | AR | 0.008759072 | 0 | Dihydrotestosterone-DHT-SRD5A1_AR | DHT-SRD5A1 - AR | DHT | Non-protein Signaling | HMRbase;uniprot |
| A-PEP.SCGN/ADRA2C | DHT-SRD5A1 | AR | 0.008577062 | 0 | Dihydrotestosterone-DHT-SRD5A1_AR | DHT-SRD5A1 - AR | DHT | Non-protein Signaling | HMRbase;uniprot |
| C-LTMR.CDH9 | DHT-SRD5A1 | AR | 0.009390258 | 0.04 | Dihydrotestosterone-DHT-SRD5A1_AR | DHT-SRD5A1 - AR | DHT | Non-protein Signaling | HMRbase;uniprot |
| C-NP.MRGPRX1/MRGPRX4 | DHT-SRD5A1 | AR | 0.008267121 | 0 | Dihydrotestosterone-DHT-SRD5A1_AR | DHT-SRD5A1 - AR | DHT | Non-protein Signaling | HMRbase;uniprot |
| A-PEP.KIT | DHT-SRD5A3 | AR | 0.000102019 | 0 | Dihydrotestosterone-DHT-SRD5A3_AR | DHT-SRD5A3 - AR | DHT | Non-protein Signaling | HMRbase;uniprot |
| A-PEP.SCGN/ADRA2C | DHT-SRD5A3 | AR | 0.000157863 | 0 | Dihydrotestosterone-DHT-SRD5A3_AR | DHT-SRD5A3 - AR | DHT | Non-protein Signaling | HMRbase;uniprot |
| C-PEP.ADORA2B | DHT-SRD5A3 | AR | 0.001623588 | 0 | Dihydrotestosterone-DHT-SRD5A3_AR | DHT-SRD5A3 - AR | DHT | Non-protein Signaling | HMRbase;uniprot |
| A-PEP.KIT | Testosterone-HSD17B12 | AR | 0.003848416 | 0.01 | Testosterone-Testosterone-HSD17B12_AR | TESTOSTERONE-HSD17B12 - AR | Testosterone | Non-protein Signaling | HMRbase;uniprot |
| A-PEP.NTRK3/S100A16 | Testosterone-HSD17B12 | AR | 0.003788878 | 0.02 | Testosterone-Testosterone-HSD17B12_AR | TESTOSTERONE-HSD17B12 - AR | Testosterone | Non-protein Signaling | HMRbase;uniprot |
| A-PEP.SCGN/ADRA2C | Testosterone-HSD17B12 | AR | 0.004443588 | 0 | Testosterone-Testosterone-HSD17B12_AR | TESTOSTERONE-HSD17B12 - AR | Testosterone | Non-protein Signaling | HMRbase;uniprot |
| C-LTMR.CDH9 | Testosterone-HSD17B12 | AR | 0.004151321 | 0.01 | Testosterone-Testosterone-HSD17B12_AR | TESTOSTERONE-HSD17B12 - AR | Testosterone | Non-protein Signaling | HMRbase;uniprot |
| C-NP.SST/CCK | Testosterone-HSD17B12 | AR | 0.003861904 | 0.01 | Testosterone-Testosterone-HSD17B12_AR | TESTOSTERONE-HSD17B12 - AR | Testosterone | Non-protein Signaling | HMRbase;uniprot |

| | | | | | | | | | |
|---|---|---|---|---|---|---|---|---|---|
| C-PEP.ADORA2B | Testosterone-HSD17B12 | AR | 0.016183172 | 0 | Testosterone-Testosterone-HSD17B12_AR | TESTOSTERONE-HSD17B12 - AR | Testosterone | Non-protein Signaling | HMRbase;uniprot |
| C-PEP.TAC1/CACNG5 | Testosterone-HSD17B12 | AR | 0.005767644 | 0 | Testosterone-Testosterone-HSD17B12_AR | TESTOSTERONE-HSD17B12 - AR | Testosterone | Non-protein Signaling | HMRbase;uniprot |
| C-PEP.TAC1/CHRNA3 | Testosterone-HSD17B12 | AR | 0.006897973 | 0 | Testosterone-Testosterone-HSD17B12_AR | TESTOSTERONE-HSD17B12 - AR | Testosterone | Non-protein Signaling | HMRbase;uniprot |
